# Gene dosage imbalance disrupts systemic metabolism in the Dp16 Down syndrome mouse model

**DOI:** 10.64898/2026.01.13.699318

**Authors:** Fangluo Chen, Muzna Saqib, Christy M. Nguyen, Dylan C. Sarver, Y. Eugene Yu, Susan Aja, Marcus M. Seldin, G. William Wong

## Abstract

Gene dosage imbalance resulting from an extra copy of human chromosome 21 (Hsa21) contributes to numerous clinical features in Down syndrome (DS). While dysregulated metabolism has long been noted in DS, the underlying cause is poorly understood and vastly understudied. To fill this critical knowledge gap, we conducted a comprehensive metabolic analysis of Dp(16)1Yey/+ mice (abbreviated Dp16), a segmental duplication model carrying ∼58% of the triplicated Hsa21 gene orthologs. Our multi-tissue transcriptomic analyses reveal shared and sex-specific increases in expression dosage of the triplicated genes in white and brown adipose tissues, liver, skeletal muscle, and hypothalamus. Despite sexual dimorphism in body weight, body temperature, food intake, and physical activity, Dp16 males and females share striking core phenotypes of pronounced insulin resistance, glucose intolerance, impaired lipid clearance, and dyslipidemia. Functional assessments, combined with biochemical, transcriptomic, and metabolomic analyses reveal tissue signatures of immune activation and a pro-inflammatory state, ER and oxidative stress, fibrosis, impaired glucose and fatty acid catabolism, altered lipid and bile acid profiles, and reduced mitochondrial respiratory capacity in Dp16 mice. These concerted changes disrupt homeostatic mechanisms that underpin metabolic health, contributing to systemic metabolic dysfunction. An obesogenic diet further exacerbates insulin resistance in Dp16 males and females despite divergent weight gain. The collective phenotypes broadly reflect the metabolic profile of DS. Our extensive molecular, biochemical, and physiological data provide an essential foundation for genetic dissection of dosage-sensitive genes affecting glucose and lipid metabolism, and for testing therapeutic strategies to improve metabolic outcomes in DS.

## INTRODUCTION

Down syndrome (DS) is the most common aneuploidy compatible with postnatal survival and it affects ∼1/700 live births (1, 2). The presence of an extra human chromosome 21 (Hsa21) in DS alters the expression dosage of a large number of triplicated genes, which has major functional impacts across many organ systems (3–7). The core clinical features of DS include cognitive deficits, craniofacial dysmorphology, hypotonia, and the development of Alzheimer (AD)-like pathology in mid-life (2, 8). With variable degrees of penetrance and expressivity, individuals with DS also frequently exhibit congenital heart defects, hearing and vision loss, leukemia, reduced bone mass, and gastrointestinal diseases (2, 8).

In the last few decades, DS research has largely been centered on understanding how trisomy 21 affects brain development and its consequences on learning and cognitive function (9). However, a major clinical feature that has frequently been overlooked and neglected, although increasingly appreciated, is that adolescents and adults with DS also have a much higher incidence of obesity, insulin resistance, type 2 diabetes, and dyslipidemia (10–19). A recent large retrospective study in the UK, spanning three decades, found that the median age at diabetes diagnosis was 15 years earlier in individuals with DS and diabetes was more than four times more common in children and young adults with DS than in individuals without DS (13). There was also an increased incidence of obesity in children and young adults with DS, with rates increasing over time (13). Despite the fact that metabolic dysfunction in DS was first noted in the 1960s (20) and well documented (10–12, 20, 21), the underlying cause is largely unknown.

Beyond clinical and epidemiological observations (10–13), only limited studies have been conducted to determine the mechanistic underpinnings of DS-associated metabolic dysfunction. Most human studies involving adolescents or adults with DS assessed the impact of trisomy 21 on food intake, adiposity, physical activity level, and energy expenditure in adolescents or adults with DS (22–30), with one study documenting a deficit in mitochondrial function in the skeletal muscle (31). However, the relative contribution of genetics versus lifestyle to altered metabolic parameters seen in DS remains challenging to untangle. At the cellular level, altered mitochondrial morphology, dynamics, and function have been well documented *in vitro* in cultured cells derived from DS (32–42); it is unclear, however, whether this translates into changes in systemic metabolism *in vivo*.

To fill this critical knowledge gap, we have recently conducted a comprehensive and in-depth analysis of changes in systemic metabolism in two trisomic DS mouse models (Ts65Dn and TcMAC21) (43, 44). Ts65Dn mice were the workhorse of DS models for over two decades (45). Due to a translocation event between mouse chromosome 16 (Mmu16) and Mmu17, Ts65Dn mice carry a freely segregating marker chromosome, Ts(17^16^), that contains ∼59% of the gene orthologs found on Hsa21 (45–47); they lack ∼70 Hsa21 gene orthologs (48). Under the basal state, chow-fed Ts65Dn mice of both sexes are glucose intolerant (44). Deterioration in metabolic homeostasis becomes much more apparent when mice are challenged with a high-fat diet (HFD). While obese Ts65Dn mice of both sexes exhibit dyslipidemia, male mice also show impaired systemic insulin sensitivity, reduced mitochondrial activity, and elevated fibrotic and inflammatory gene signatures in the liver and adipose tissue. Our systems-level analysis also reveals major changes in gene connectivity and pathways in liver and adipose tissues that contribute to dysregulated glucose and lipid metabolism seen in Ts65Dn mice (44). The metabolic phenotypes of Ts65Dn mice are largely consistent with the clinical and epidemiological findings of DS, namely the proclivity of individuals with DS to develop diabetes (10–13), dyslipidemia (16, 18, 19, 49), and reduced mitochondrial function (32). However, Ts65Dn has one major limitation. In addition to the 103 triplicated Hsa21 gene orthologs, Ts65Dn mice also carry an additional 41 triplicated protein-coding genes (from the sub-centromeric region of Mmu17) unrelated to Hsa21 (50, 51). This confounds and complicates the genotype-phenotype correlations in this mouse model (52, 53).

TcMAC21 is a recently generated transchromosomic mouse model carrying a non-mosaic Hsa21q (54). Due to multiple deletions, TcMAC21 mice are missing ∼7% of the Hsa21 genes located on the long arm.

Our systematic analysis of TcMAC21 mice has led to completely unexpected findings—the TcMAC21 mice are hypermetabolic, showing a dramatic increase in mitochondrial respiration and energy expenditure, and are markedly leaner despite consuming greater amounts of food, and are strikingly more insulin sensitive (43). These phenotypes are inconsistent with the clinical profile of DS. Thus, TcMAC21 mice do not model the metabolic phenotypes seen in DS. This phenomenon may be partly caused by abnormal interactions between human and mouse proteins, including the orthologous proteins of these two species, as well as the presence of over 400 non-coding human genes with uncertain effects on the mouse transcriptome (43).

To overcome the caveats and limitations associated with the metabolic studies of Ts65Dn and TcMAC21, and to help inform the selection of mouse model that best reflects the metabolic profile of DS, we undertook a comprehensive metabolic analysis of Dp(16)1Yey/+ (abbreviated Dp16), another widely used mouse model of DS (55). The Dp16 mice carry a duplicated segment of Mmu16 syntenic to Hsa21, with 115 triplicated Hsa21 gene orthologs (55). Unlike the Ts65Dn mice generated from chromosomal translocation (46, 47), the Dp16 mice were generated by chromosomal engineering based on precise Cre/LoxP-mediated recombineering (55) and therefore carry no extra non-Hsa21 gene orthologs. Also, unlike the TcMAC21 mice, all the triplicated Hsa21 gene orthologs in Dp16 mice are of mouse origin. The Dp16 mouse model thus allows us to address the contribution of gene dosage imbalance to changes in systemic metabolism.

While we observed striking sex differences in various metabolic parameters, Dp16 male and female mice also shared important metabolic deficits that are hallmarks of dysregulated systemic metabolism; both sexes developed pronounced glucose intolerance, insulin resistance, impaired lipid clearance, and dyslipidemia, and these phenotypes were further exacerbated by a high-fat diet. Using a multi-omics approach, we showed that Dp16 mice displayed transcriptomic, metabolomic, and biochemical signatures associated with tissue fibrosis, oxidative stress, a pro-inflammatory state, impaired metabolism and mitochondrial function, and dyslipidemia. Our pathway enrichment analyses revealed global changes in gene expression and biological pathways across major metabolic tissues in Dp16 mice. These collective changes underpin and contribute to the metabolic dysfunction seen in Dp16 mice. Collectively, our data suggest that dosage imbalance arising from the triplication of Hsa21 gene orthologs, along with its cascading effects across tissue transcriptomes and metabolomes, are causally linked to insulin resistance and impaired systemic glucose and lipid metabolism. Our present study lays critical and essential groundwork for genetic dissection of dosage-sensitive genes underpinning widespread metabolic deficits in DS.

## MATERIALS AND METHODS

### Mouse model

Dp(16)1Yey/+ (abbreviated Dp16) mice and wild-type (WT) littermate controls, on C57BL/6J genetic background, were obtained from the Jackson Laboratory (Strain # 013530). Mice were fed a standard chow (Envigo; 2018SX) or a high-fat diet (HFD; 60% kcal derived from fat, #D12492, Research Diets, New Brunswick, NJ). Mice were housed in polyethylene terephthalate (PET) cages on a 12h:12h light-dark photocycle (lights on at 6 am, lights off at 6 pm) with *ad libitum* access to water and food. For the HFD-fed group, HFD was provided beginning for 26-34 weeks. At termination of the study, all mice were fasted for 2 h and euthanized. The age of mice at the time of tissue harvest: male and female mice fed a standard chow were 27.5 weeks old; male mice fed an HFD were 50 weeks old (on HFD for 34.5 weeks); female mice fed an HFD were 45 weeks old (on HFD for 26 weeks). Tissues were collected, snap-frozen in liquid nitrogen, and kept at 80°C until analysis.

All mouse protocols were approved by the Institutional Animal Care and Use Committee of the Johns Hopkins University School of Medicine (animal protocol # MO22M367). All animal experiments were conducted in accordance with the National Institute of Health guidelines and followed the standards established by the Animal Welfare Acts.

### Body composition analysis

Body composition analyses for total fat mass, lean mass, and water content were determined using a quantitative magnetic resonance instrument (Echo-MRI-100, Echo Medical Systems, Waco, TX) at the Mouse Phenotyping Core facility at Johns Hopkins University School of Medicine.

### Indirect calorimetry

Chow- or HFD-fed Dp16 male and female mice and WT littermates were used for simultaneous assessments of daily body weight change, food intake (corrected for spillage), physical activity, and whole-body metabolic profile in an open flow indirect calorimeter (Comprehensive Laboratory Animal Monitoring System, CLAMS; Columbus Instruments, Columbus, OH) as previously described (56). In brief, data were collected for three days to confirm mice were acclimatized to the calorimetry chambers (indicated by stable body weights, food intakes, and diurnal metabolic patterns), then data were analyzed for the subsequent three days. Mice were observed with *ad libitum* access to food, throughout the fasting process, and in response to refeeding. Rates of oxygen consumption (*V͘*_O2_; mL·kg^-1^·h^-1^) and carbon dioxide production (*V͘*_CO2_; mL·kg^-1·^h^-1^) in each chamber were measured every 24 min. Respiratory exchange ratio (RER =

*V͘*_CO2_/*V͘*_O2_) was calculated by CLAMS software (version 5.18) to estimate relative oxidation of carbohydrates (RER = 1.0) versus fats (RER = 0.7), not accounting for protein oxidation. Energy expenditure (EE) was calculated as EE= *V͘*_O2_× [3.815 + (1.232 × RER)] and normalized to lean mass. We also performed ANCOVA analysis on EE using body weight as a covariate (57). Physical activities (total and ambulatory) were measured by infrared beam breaks in the metabolic chamber. Average metabolic values and summed intake and activity values were calculated per subject and averaged across subjects for statistical analysis by Student’s t-test.

### Body Temperature

Deep colonic temperature was measured by inserting a lubricated (Medline, water soluble lubricating jelly, MDS032280) probe (Physitemp, BAT-12 Microprobe Thermometer) into the anus of mice at a depth of 2 cm. Stable numbers were recorded in both the dark and light cycle for each mouse.

### Fecal bomb calorimetry and assessment of fecal parameters

Fecal pellet frequency and average fecal pellet weight were monitored by housing each mouse singly in clean cages and counting the number of fecal pellets and recording their weight at the end of a 24 h period. Fecal pellets were shipped to the University of Michigan Animal Phenotyping Core for fecal bomb calorimetry. Briefly, fecal samples were dried overnight at 50°C prior to weighing and grinding them to powder. Each sample was mixed with wheat flour (90% wheat flour, 10% sample) and formed into 1.0 g pellet, which was then secured into the firing platform and surrounded by 100% oxygen. The bomb was lowered into a water reservoir and ignited to release heat into the surrounding water. Together these data were used to calculate fecal pellet frequency (bowel movements/day), average fecal pellet weight (g/bowel movement), fecal energy (cal/g feces), and total fecal energy (kcal/day).

### Glucose, insulin, and lipid tolerance tests

All tolerance tests were conducted as previously described (58–60). For glucose tolerance tests (GTTs), mice were fasted for 6 h before glucose injection. Glucose (Sigma, St. Louis, MO) was reconstituted in saline (0.9 g NaCl/L) to a final concentration of 1 g/10 mL (for the chow-fed mice) or 2g/10 mL (for the HFD-fed mice), sterile-filtered, and injected intraperitoneally (i.p.) at 1 mg/g body weight (i.e., 10 μL/g body weight for chow-fed mice or 5 μL/g body weight of HFD-fed mice). Blood glucose was measured at 0, 15, 30, 60, and 120 min after glucose injection using a glucometer (NovaMax Plus, Billerica, MA). For insulin tolerance tests (ITTs), food was removed 2 h before insulin injection. 6.5 μL of insulin stock (4 mg/mL; Gibco) was diluted in 10 mL of saline, sterile-filtered, and injected i.p. at 0.75 U/kg body weight (i.e., 10 uL/g body weight). Blood glucose was measured at 0, 15, 30, 60, and 90 min after insulin injection using a glucometer (NovaMax Plus). For lipid tolerance tests (LTTs), mice were fasted for 12 h and then injected i.p. with 20% emulsified Intralipid (soybean oil; Sigma; 10 μL/g of body weight). Sera were collected via tail bleed using a Microvette® CB 300 (Sarstedt) at 0, 1, 2, 3, and 4 h post-injection. Serum triglyceride levels were quantified using kits from Infinity Triglycerides (Thermo Scientific).

### Fasting glucose, insulin, and lipid profile

Mice were fasted overnight (∼16 h), beginning at 1 h before the dark cycle (around 5 pm). Clean cages were provided before food withdrawal. Overnight fasting blood glucose levels from tail bleed were measured using a glucometer. Serum was collected at the 16 h fast (around 10 am in the morning) for insulin ELISA, as well as for the quantification of triglyceride, cholesterol, non-esterified free fatty acids (NEFA), and β-hydroxybutyrate concentrations.

### Blood and tissue chemistry analysis

Tail vein blood samples were allowed to clot on ice and then centrifuged for 10 min at 10,000 x *g*. Serum samples were stored at -80°C until analyzed. Serum triglycerides (TG) and cholesterol were measured according to manufacturer’s instructions using an Infinity kit (Thermo Fisher Scientific, Middletown, VA). Non-esterified free fatty acids (NEFA) were measured using a Wako kit (Wako Chemicals, Richmond, VA). Serum β-hydroxybutyrate (ketone) concentrations were measured with a StanBio Liquicolor kit (StanBio Laboratory, Boerne, TX). Serum insulin (Crystal Chem, 90080), T3 (Calbiotech, T3043T-100), testosterone (Cayman, 582701), estradiol (Cayman, 501890), corticosterone (Cayman, 501320), and alanine aminotransferase (ALT; abcam, ab282882) levels were measured using commercial kits according to manufacturer’s instructions.

Hydroxyproline assay (Sigma Aldrich, MAK569) was used to quantify total collagen content in liver, hypothalamus, and adipose tissues according to the manufacturer’s instructions, with specific optimizations for each tissue type. Tissues were homogenized in deionized water using a bead mill homogenizer. Liver, iWAT, and gWAT were homogenized to a concentration of 0.1mg tissue/µL. Due to the small and variable tissue mass, hypothalami were homogenized in a fixed volume of 110–120µL to yield sufficient volume for processing. All samples were hydrolyzed with an equal volume of HCl at 120°C for 3 hours. The volume of hydrolyzed tissue samples plated for the dehydration step was optimized for each tissue type and experimental group to ensure all measurements fell within the linear range of the standard curve. Hydroxyproline content was calculated based on a standard curve and normalized to tissue weight.

Lipid peroxidation levels (marker of oxidative stress) in liver, hypothalamus, and adipose tissues were assessed by the quantification of malondialdehyde (MDA) levels via the Thiobarbituric Acid Reactive Substances (TBARS) assay (Cayman Chemical, 700870) according to the manufacturer’s instructions.

### Serum lipoprotein-triglyceride and cholesterol analysis by FPLC

Food was removed for ∼2 h (in the light cycle) prior to blood collection. Sera collected from mice were pooled (*n* = 10-14/group) and sent to the Mouse Metabolism Core at Baylor College of Medicine for fast protein liquid chromatography (FPLC) separation. A total of 45 fractions were collected, and TG and cholesterol in each fraction were quantified.

### Extraction of hepatic lipids

Lipid extraction from frozen liver samples was performed using a modified Folch method (61). Briefly, approximately 25 mg of frozen liver tissue was weighed and homogenized in 400 µL of cold sucrose buffer (250 mM sucrose, 10 mM Tris, 1 mM EDTA) using a bead beater (FastPrep-24, MP Biomedical). The samples underwent three rounds of 20-second bead beating cycles. Lipids were then extracted from the liver homogenate by adding 1.5 mL of a chloroform:methanol (2:1, v/v) solution to the mixture. The sample was vortexed thoroughly and centrifuged at 1700 rpm for 5 minutes at 4℃ to separate the phases. The lower chloroform phase, containing lipids, was carefully transferred to a new tube and split equally into two separate tubes. Each tube was then dried in a speed vacuum to dehydrate samples. The dried lipid extracts were reconstituted in 50 µL of chloroform for lipid analysis and quantification by thin-layer chromatography (TLC) or in 50 µL of tert-butanol:methanol:TritonX-100 solution (3:1:1, v/v/v) for cholesterol quantification. Reconstituted samples were either used immediately or stored at -80°C until analysis.

### Separation of lipid classes and quantification by thin-layer chromatography (TLC)

TLC was performed as described by Ring et al. (62) to separate specific classes of lipids. To separate and quantify lipid classes from liver tissue, silica gel (Analtech, Preadsorbent Silica Gel G UNIPLATES Channeled, 20x20cm, 250 µm, Cat #P31911) plates were pre-washed in methanol, air-dried, and then subsequently equilibrated by pre-washing in hexane:diethyl ether:acetic acid (H:D:A) (80:20:1, v/v/v) solvent system. The plates were allowed to air-dry completely before lipid spotting. Lipid extracts (2 µL) and lipid standards (2 µL, 5mg/mL in chloroform, Millipore Sigma, Cat# 1787-1AMP, Supelco) were spotted onto the plates, and lipids were separated in H:D:A (80:20:1, v/v/v) to resolve triacylglycerols (TAG) and diacylglycerols (DAG). After development, the plates were air-dried and exposed to iodine vapor in a pre-equilibrated tank overnight to visualize lipid spots. For quantification, the plates were imaged, and lipid spots were analyzed using ImageJ software (https://imagej.net/) to determine the intensity of each lipid spot relative to the known lipid standards and normalized to tissue weight.

### Hepatic cholesterol quantification

Following tissue extraction, total cholesterol content was quantified using the Infinity Cholesterol Reagent kit (Thermo Fisher Scientific, Middletown, VA) according to the manufacturer’s instructions. Quantified cholesterol values were normalized to tissue weight.

### Untargeted serum and liver metabolomic analyses

Serum and liver metabolites were extracted and subjected to LC-MS/MS detection on a Q Exactive HF-X Quadrupole-Orbitrap mass spectrometer system at Novogene (Sacramento, CA) (*n* = 6 mice per genotype per sex). Data were processed using Novogene in-house analysis pipeline. In brief, the raw mass spectrometry data were first converted to mzXML format using ProteoWizard (63). Peak extraction, alignment, and retention time correction were then performed with XCMS software (64). The total peak area within each sample was normalized, and peaks with a missing rate greater than 50% across sample groups were filtered out. The corrected and filtered results were matched with the Novogene local database to obtain metabolite identification information. A total of 4182 metabolites were identified from the 48 samples. Multivariate statistical analysis was conducted on the metabolites, including Principal Component Analysis (PCA) and Partial Least Squares Discriminant Analysis (PLS-DA). PLS-DA is a supervised discriminant analysis statistical method. This method uses partial least squares regression (65) to establish the relationship model between the relative quantitative value of metabolites and the sample category to realize the prediction of the sample category. The PLS-DA model of each comparison group was established, and the model evaluation parameters (R2, Q2) obtained by 7-cycle cross-validation. Differential metabolites were screened according to the criteria: VIP > 1.0, fold change (FC) > 1.2 or FC < 0.833 and *P*-value < 0.05. VIP refers to the variable importance in the projection of the first principal component of the PLS-DA model, and the VIP value represents the contribution of the metabolites to the grouping. KEGG enrichment (FDR correction by Benjamini and Hochberg method) and GSEA analysis was performed on KEGG entries based on the changes in quantitative values of metabolites. All metabolomics data, raw spectral files, and details of experimental protocol and data analyses have been deposited in a public repository, the Metabolomics Workbench (66).

### Mitochondrial respirometry

Respirometry was conducted on frozen tissue samples to assay for mitochondrial activity as described previously (67, 68). Briefly, liver and brown adipose tissue (BAT) were dissected, snapped frozen in liquid nitrogen, and stored at -80°C for later analysis. Samples were thawed in MAS buffer (70mM sucrose, 220 mM mannitol, 5 mM KH_2_PO_4_, 5 mM MgCl_2_, 1 mM EGTA, 2 mM HEPES pH 7.4), finely minced with scissors, and then homogenized with a glass Dounce homogenizer. The resulting homogenate was spun at 1000 *g* for 10 min at 4°C. The supernatant was collected and immediately used for protein quantification by BCA assay (Thermo Scientific, 23225). Each well of the Seahorse microplate was loaded with 4 µg (BAT) or 8 µg (liver) homogenate protein. Each biological replicate consists of three technical replicates. Samples from all tissues were treated separately with NADH (1 mM) as a complex I substrate or Succinate (a complex II substrate, 5 mM) in the presence of rotenone (a complex I inhibitor, 2 µM), then with the inhibitors rotenone (2 µM) and Antimycin A (4 µM), followed by TMPD (0.45 mM) and Ascorbate (1 mM) to activate complex IV, and finally treated with Azide (40 mM) to assess non-mitochondrial respiration. All mitochondrial respiration data were normalized to mitochondrial content, quantified using MitoTracker Deep Red (MTDR, ThermoFisher, M22426) as described (67, 68). Briefly, lysates were incubated with MTDR (1 µM) for 10 min at 37°C, then centrifuged at 2000 *g* for 5 min at 4°C. The supernatant was carefully removed and replaced with 1x MAS solution and fluorescence was read with excitation and emission wavelengths of 625 and 670 nm, respectively. To minimize non-specific background signal contribution, control wells were loaded with MTDR and 1x MAS and subtracted from all sample values.

### RNA-sequencing and bioinformatics analysis

Bulk RNA sequencing of Dp16 (*n* = 6) and WT (*n* = 6) mouse liver, gWAT, iWAT, brown adipose tissue (BAT), skeletal muscle (gastrocnemius), pancreas, and hypothalamus were performed by Novogene (Sacramento, California, USA) on a NovaSeq X Plus platform and pair-end reads (2 x 150 bp) were generated, with 6 G raw data per sample. One gWAT sample from the Dp16 female mice failed the initial quality control test and was excluded from subsequent RNA sequencing. Sequencing data was analyzed using the standard Novogene Analysis Pipeline. Sequencing reads were aligned to Mus musculus reference genome (GRCm39/mm39). Data analysis was performed using a combination of programs, including Fastp, Hisat2, and FeatureCounts. Differential expressions were determined through DESeq2. The resulting *P*-values were adjusted using the Benjamini and Hochberg’s approach for controlling the false discovery rate. Genes with an adjusted *P*-value < 0.05 and log2(FC) > 0.5 were assigned as differentially expressed. Gene ontology (GO), Kyoto Encyclopedia of Genes and Genomes (KEGG), and Reactome (http://www.reactome.org) enrichment were implemented by ClusterProfiler. All volcano plots and heat maps were generated in Graphpad Prism 10 software. All statistics were performed on log transformed data. All heat maps were generated from column z-score transformed data. The z-score of each column was determined by taking the column average, subtracting each sample’s individual expression value by said average then dividing that difference by the column standard deviation. Z-score = (value – column average)/column standard deviation. High-throughput sequencing data from this study have been submitted to the NCBI Sequence Read Archive (SRA) under accession number # PRJNA1160420.

### Statistical analyses

All results are expressed as mean ± standard error of the mean (SEM). Statistical analysis was performed with GraphPad Prism 10 software (GraphPad Software, San Diego, CA). Data were analyzed with two-tailed Student’s *t*-tests or by repeated measures ANOVA. For two-way ANOVA, we performed Sidek or Bonferroni post hoc tests. *P* < 0.05 was considered statistically significant.

## RESULTS

### Increased gene expression dosage of the triplicated Hsa21 gene orthologs across major metabolic tissues

The Dp16 mice are triplicated for ∼58% of Hsa21 gene orthologs (55, 69) (**Fig. 1A**). First, we wanted to establish which among the 115 triplicated Hsa21 gene orthologs on Mmu16 are expressed in six major metabolic tissues: gonadal white adipose tissue (gWAT; visceral fat), inguinal white adipose tissue (iWAT; subcutaneous fat), brown adipose tissue (BAT), skeletal muscle (gastrocnemius), and hypothalamus. Bulk RNA-sequencing showed that the majority of the triplicated genes are indeed expressed at the expected higher dosage (1.5-fold or higher) across tissues and their expression are regulated in a tissue- and sex-specific manner (**Fig. 1B,C**). Interestingly, female gWAT is the only tissue with 8 triplicated Hsa21 gene orthologs (*Rbm11*, *Chodl*, *Cldn8*, *Sh3bgr*, *Igsf5*, *Itgb2l*, *Pcp4* and *Tmprss2*) whose expressions are significantly suppressed relative to WT controls (**Fig. 1B**). Except female gWAT, we observed little evidence of dosage compensation, consistent with recent findings (70). Of the six tissues examined, hypothalamus expresses the largest number of triplicated Hsa21 gene orthologs, as well as the largest number of shared triplicated genes between males and females (**Fig. 1C**). Visceral fat (gWAT) and skeletal muscle (gastrocnemius) have the lowest number of shared differentially expressed triplicated genes between sexes, with females expressing a significantly higher number of triplicated genes compared to males (**Fig. 1C**). Together, these data indicate increased expression dosage of many triplicated Hsa21 gene orthologs across major metabolic tissues in Dp16 mice, with sex differences noted.

**Figure 1.**
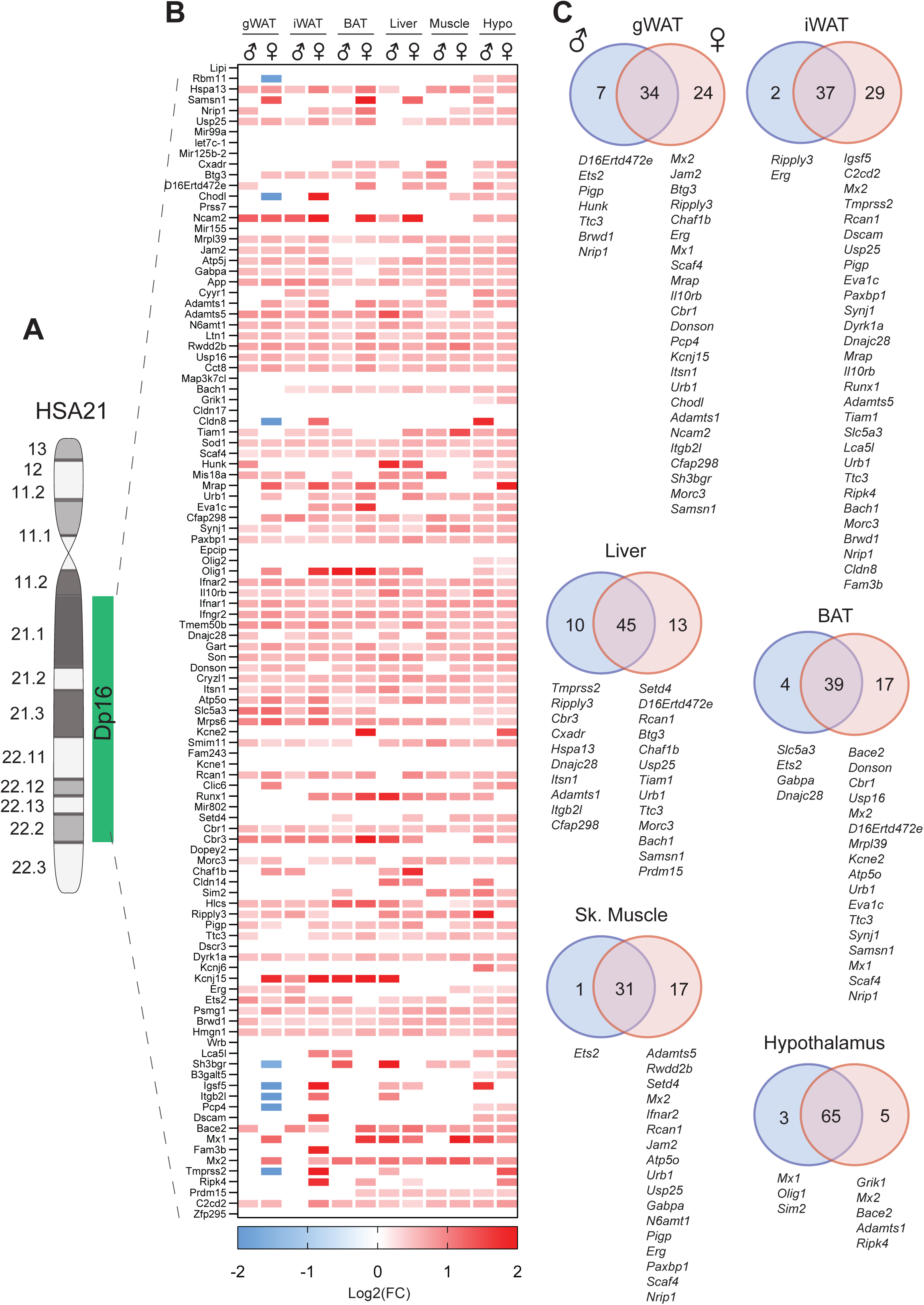
Increased gene expression dosage of the triplicated Hsa21 gene orthologs on mouse chromosome 16 (Mmu16) across tissues. **A)** Graphical representation of human chromosome 21 (Hsa21) and the syntenic Mmu16 segment that is duplicated in Dp16 mice. **B)** Global view of the expression of 108 triplicated Hsa21 gene orthologs on Mmu16 in gonadal white adipose tissue (gWAT), inguinal white adipose tissue (iWAT), interscapular brown adipose tissue (BAT), skeletal muscle (gastrocnemius), and hypothalamus. Red denotes transcript that is expressed at ≥1.5-fold the WT level, whereas blue denotes transcript that is expressed at significantly lower level compared to WT control. The Ktrap gene cluster (23 Ktrap genes) located between Cldn8 and Tiam1is not shown. **C)** Overlap analysis showing differentially expressed Hsa21 gene orthologs that are shared between males and females across six tissues. The criteria for differentially expressed genes (DEGs) is log2(FC) > 0 with padj ≤ 0.05. *n* = 6 RNA samples per genotype per sex per tissue-type. Chow-fed WT and Dp16 mice were at 27.5 weeks of age at the time of tissue collection.

### Sexual dimorphism in body weight, food intake, physical activity, and body temperature in Dp16 mice

We next determined the impact of gene dosage imbalance on systemic metabolism under the basal state when mice were fed a standard chow. The body weights of Dp16 male mice from 7 to 16 weeks old were not different from WT controls (**Fig. 2A**). Body composition analysis by NMR revealed a modest increase in fat mass in Dp16 male mice but no difference in lean mass (**Fig. 2B**). At 27.5 weeks of age, body weights and the absolute and relative (% of body weight) weights of visceral (gWAT) and subcutaneous (iWAT) fat, liver, and heart were not different between genotypes in male mice (**Figure 2 – figure supplement 1**). The absolute and relative weights of the kidney, however, were significantly higher in Dp16 male mice. Female body weights were similar between genotypes from 6 to 8 weeks of age; however, Dp16 females gained markedly more weight compared to WT controls from 9 weeks of age onward (**Fig. 2C**). Increased body weight in Dp16 females was due to increased fat and lean mass (**Fig. 2D**). At 27.5 weeks of age, body weights and the absolute weights of gWAT and iWAT were significantly higher in Dp16 females compared to WT controls (**Figure 2 – figure supplement 1**). The absolute and relative weights of liver, heart, and kidney were not different between genotypes in female mice.

**Figure 2.**
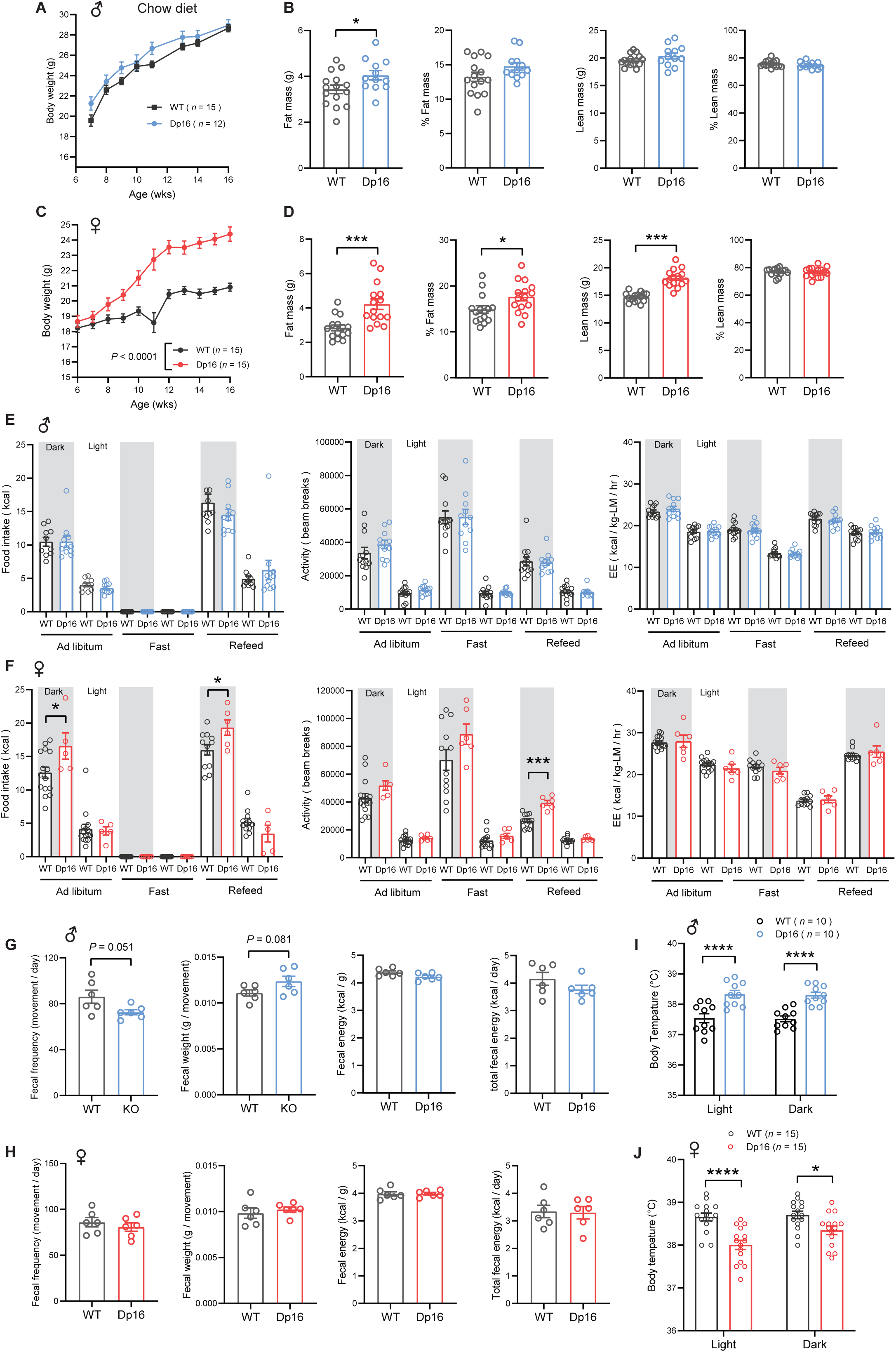
Sexually dimorphism in body weight, body temperature, food intake, and physical activity in chow-fed Dp16 mice. **A)** Body weight of chow-fed male Dp16 and WT mice over time. **B)** absolute and relative (% of body weight) fat and lean mass in male mice (WT = 15; Dp16 = 12). **C)** Body weight of chow-fed female Dp16 and WT mice over time. **D)** absolute and relative (% of body weight) fat and lean mass in female mice (WT = 15; Dp16 = 15). **E-F)** Food intake, total physical activity level, and energy expenditure of male (E) and female (F) Dp16 and WT mice across the circadian cycle (light and dark) and metabolic states (*ad libitum* fed, fast, refeed). Sample size for male (WT = 10-12; Dp16 = 11-12) and female (WT = 13-15; Dp16 = 5-6) mice. **G-H)** fecal frequency, average fecal weight, and fecal energy content (per gram and total) in male (G) and female (H) Dp16 and WT mice. Sample size for male (WT = 6; Dp16 = 6) and female (WT = 6; Dp16 = 6) mice. **I-J)** Body temperature in the light and dark cycle of male (I) and female (J) Dp16 and WT mice. Sample size for male (WT = 10; Dp16 = 10) and female (WT = 15; Dp16 = 15) mice. All data are presented as mean ± SEM. * *P*<0.05; *** *P*<0.001; **** *P*<0.0001. For body weight over time, data were analyzed by 2-way ANOVA with Sidek post hoc tests.

We performed indirect calorimetry analysis to determine food intake, physical activity level, metabolic rate (VO_2_), and energy expenditure across the circadian cycle (light and dark phase) and metabolic states (*ad libitum* fed, fasted, and refed). None of the parameters measured were different between genotypes in male mice (**Fig. 2E**). In females, however, food intake was significantly higher in Dp16 mice compared to WT controls in the *ad libitum* fed state and during the refeeding period following a fast (**Fig. 2F**). Physical activity levels were also higher in Dp16 females during the refed period. Energy expenditure (normalized to lean mass), however, was not different between Dp16 females and WT controls across the circadian cycles and metabolic states (**Fig. 2F**). Normalization of energy expenditure to lean mass can lead to an overestimation of energy expenditure (57). We therefore also performed ANCOVA analyses (using lean mass as a covariate of energy expenditure) (57). Both types of analyses indicated no differences in energy expenditure between genotypes of either sex across the circadian cycles and metabolic states (**Figure 2 – figure supplement 2**).

To account for any potential differences in nutrient absorption in the intestine, we measured fecal output, frequency, and weight, as well as the fecal energy content by fecal bomb calorimetry. None of the fecal parameters were significantly different between genotypes of either sex (**Fig. 2G-H**). Interestingly, Dp16 males had higher body temperature compared to WT controls in both the light and dark cycle, whereas Dp16 females had lower body temperature (**Fig. 2I-J**). This prompted us to assess whether Dp16 mice have altered mitochondrial function in brown adipose tissue (BAT), a major thermogenic tissue. Despite differences in body temperature, both Dp16 males and females had reduced maximal mitochondrial respiratory capacity in BAT, an effect more pronounced in males (**Figure 2 – figure supplement 3**). Circulating Triiodothyronine (T3) levels, a hormone that also controls body temperature, were not different between genotypes of either sex (**Figure 2 – figure supplement 4**). We measured serum levels of sex and stress hormones, as these could contribute to sex differences in metabolic outcomes. Corticosterone levels were not different between genotypes of either sex. Testosterone levels in males were variable and not significantly different between genotypes, whereas estradiol levels were higher in Dp16 females (**Figure 2 – figure supplement 4**), possibly contributing to greater physical activity (71, 72) and lower body temperature (73, 74). Our results suggest that lower body temperature, coupled with higher food intake, likely contributes to greater weight gain over time in Dp16 females. Together, these data reveal striking sex differences in body weight, food intake, physical activity, and body temperature in Dp16 mice.

**Glucose intolerance, insulin resistance, impaired lipid clearance, and dyslipidemia in Dp16 mice** Type 2 diabetes and altered lipid profile are well documented in DS population (10–19). To determine baseline insulin, glucose, and lipid profiles, we measured blood glucose, serum insulin, triglyceride (TG), cholesterol, non-esterified free fatty acids (NEFA), and β-hydroxybutyrate (BHB; ketone) in overnight fasted (16 h) mice. The Dp16 males fed a standard chow had significantly higher fasting insulin levels (**Fig. 3A**). Fasting hyperinsulinemia likely contributed to lower fasting blood glucose seen in Dp16 male mice. Serum TG and NEFA levels were also significantly higher in Dp16 males compared to WT controls, suggesting enhanced fat mobilization in the fasted state. Serum β-hydroxybutyrate (ketone) levels were not different between genotypes whereas serum cholesterol levels were markedly lower in Dp16 male mice (**Fig. 3A**). Like the males, Dp16 females also had significantly higher fasting serum insulin levels (**Fig. 3B**). Unlike the males, blood glucose, triglycerides, cholesterol, and ketone levels were not different between genotypes in females. Also, different from the males, Dp16 females had lower NEFA levels relative to WT controls.

**Figure 3.**
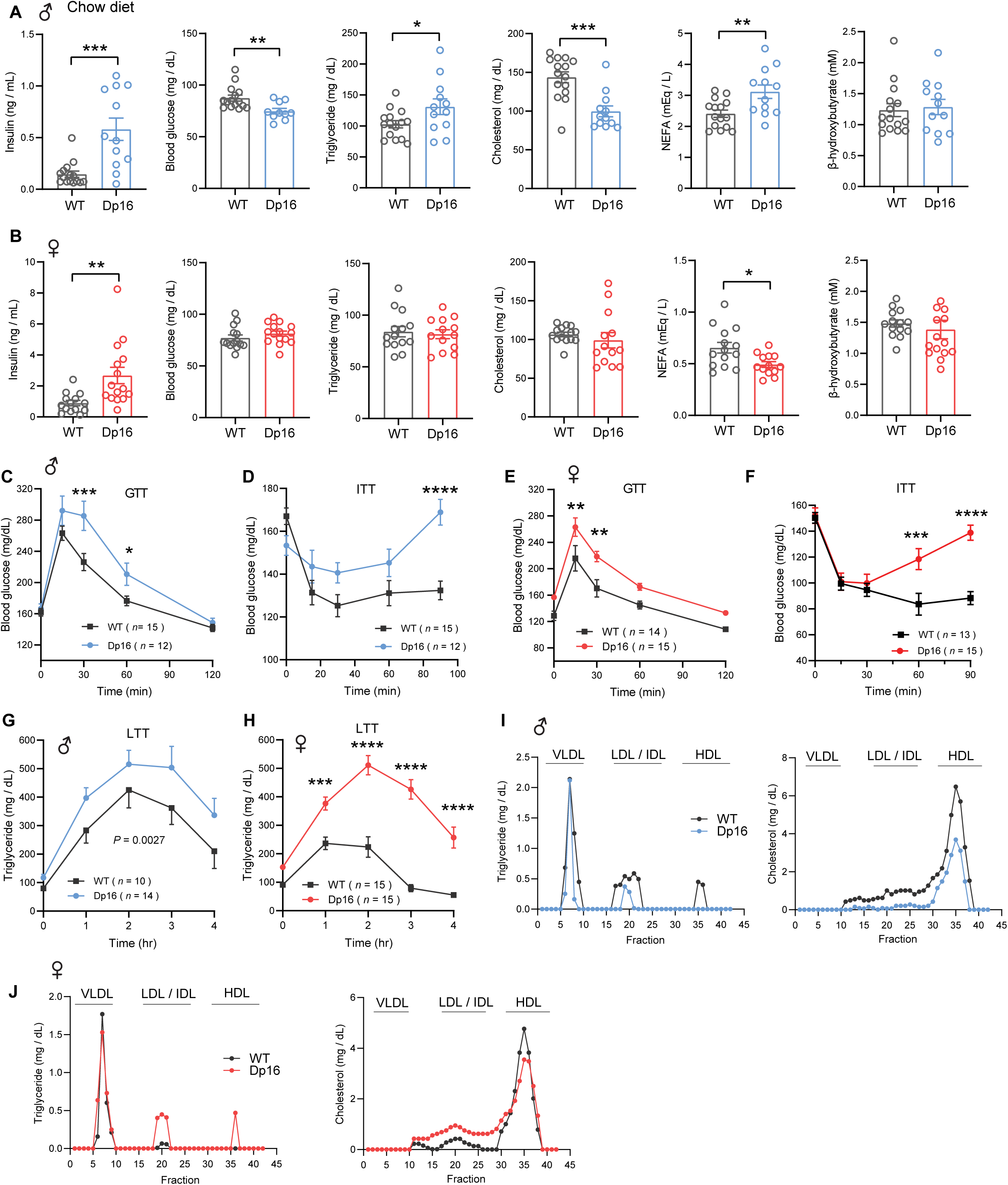
Glucose intolerance, insulin resistance, and impaired lipid clearance in chow-fed Dp16 mice. A-B) Overnight fasting insulin, blood glucose, serum triglyceride, cholesterol, non-esterified free fatty acids (NEFA), and β-hydroxybutyrate (ketone) in male (A) and female (B) Dp16 and WT mice. Sample size for male mice (WT = 15; Dp16 = 12) and female mice (WT = 14; Dp16 = 15). **C-F)** Impaired glucose tolerance as determined by the glucose tolerance test (GTT) in male (C) and female (E) Dp16 mice compared to WT controls. Impaired insulin sensitivity as determined by the insulin tolerance test (ITT) in male (D) and female (F) Dp16 compared to WT controls. Sample size for male mice (WT = 15; Dp16 = 12) and female mice (WT = 14; Dp16 = 15). **G-H)** Impaired triglyceride clearance in response to lipid gavage as determined by the lipid tolerance test (LTT) in male (G) and female (H) Dp16 relative to WT controls. Sample size for male mice (WT = 10; Dp16 = 14) and female mice (WT = 15; Dp16 = 15). **I-J)** Pooled mouse sera from male (I) and female (J) Dp16 and WT mice were fractionated by fast protein liquid chromatography (FPLC), and the triglyceride and cholesterol content of each fraction was quantified. Fractions corresponding to very-low density lipoprotein (VLDL), low-density lipoprotein (LDL), intermediate-density lipoprotein (IDL), and high-density lipoprotein (HDL) are indicated. All data are presented as mean ± SEM. * *P*<0.05; ** *P*<0.01; *** *P*<0.001; **** *P*<0.0001. For all tolerance tests, data were analyzed by 2-way ANOVA with Sidek post hoc tests.

Higher fasting insulin levels in Dp16 males and females suggest insulin resistance. To further assess glucose metabolism in these mice, we performed glucose and insulin tolerance tests to determine the rate of glucose clearance in response to glucose or insulin injection. Both Dp16 males and females showed impaired glucose clearance after glucose loading (**Fig. 3C,E**). To confirm that Dp16 mice have impaired insulin action, we directly assessed insulin sensitivity via insulin tolerance test (ITT). The rate of glucose clearance in response to insulin injection was markedly impaired in both Dp16 males and females (**Fig. 3D,F**). Impaired insulin action, glucose intolerance, and fasting hyperinsulinemia strongly indicate an insulin resistance phenotype in Dp16 mice.

We next assessed whether Dp16 mice have altered lipid handling capacity by performing a lipid tolerance test. The rate of triglyceride clearance in response to an acute lipid load was significantly impaired in Dp16 mice of either sex, with Dp16 females showing a much more striking deficit in lipid clearance (**Fig. 3G,H**). To determine whether Dp16 mice have altered lipoprotein profiles, we subjected pooled sera to FPLC fractionation followed by the quantification of triglyceride and cholesterol levels in each fraction.

Dp16 males had lower triglyceride and cholesterol levels in the LDL/IDL and HDL fractions (**Fig. 3I**). In contrast, Dp16 females had higher triglyceride and cholesterol levels in the LDL/IDL fractions, as well as higher triglyceride levels in the HDL fractions. Since liver is a key organ in lipid and lipoprotein synthesis and export, we measured hepatic triglyceride (TG), diacylglycerol (DAG), and cholesterol contents. No genotypic differences in either sex was observed (**Fig. 3 – figure supplement 1**). Dp16 males had lower maximal liver mitochondrial respiratory capacity, though not significant; and this was not different between genotypes in females (**Fig. 3 – figure supplement 2**). Altogether, our data indicate that Dp16 mice of either sex developed pronounced insulin resistance, glucose intolerance, dyslipidemia, and impaired lipid clearance.

### Altered hepatic and serum metabolome in Dp16 mice

Since Dp16 mice showed profound disturbances in systemic energy metabolism, we performed untargeted metabolomic analyses to assess possible changes in liver and serum metabolome. A total of 4182 metabolites were identified from the 48 serum and liver samples. Partial Least Square Discriminant Analysis (PLS-DA) indicated that liver and serum metabolome of Dp16 males and females are clearly distinguishable from that of their corresponding WT controls (**Fig. 4A,B**). In male and female liver, we observed 319 and 337 differential metabolites, respectively, in Dp16 mice vs WT controls (**Fig. 4 – figure supplement 1; Fig. 4 – source data 1-4**). In Dp16 male and female serum samples, we observed 422 and 835 differential metabolites, respectively (**Fig. 4 – figure supplement 1**). When comparing the differential metabolites found in liver and serum, there appeared to be limited overlap between the two compartments (**Fig. 4C**). Major sex differences were seen in the differential metabolites found in liver and serum. There were 32 differential metabolites shared between Dp16 male and female liver, and 155 differential serum metabolites shared between the sexes (**Fig. 4D**). The majority of differential metabolites found in liver and serum, however, were not shared between the sexes (**figure 4 – source data 5**).

**Figure 4.**
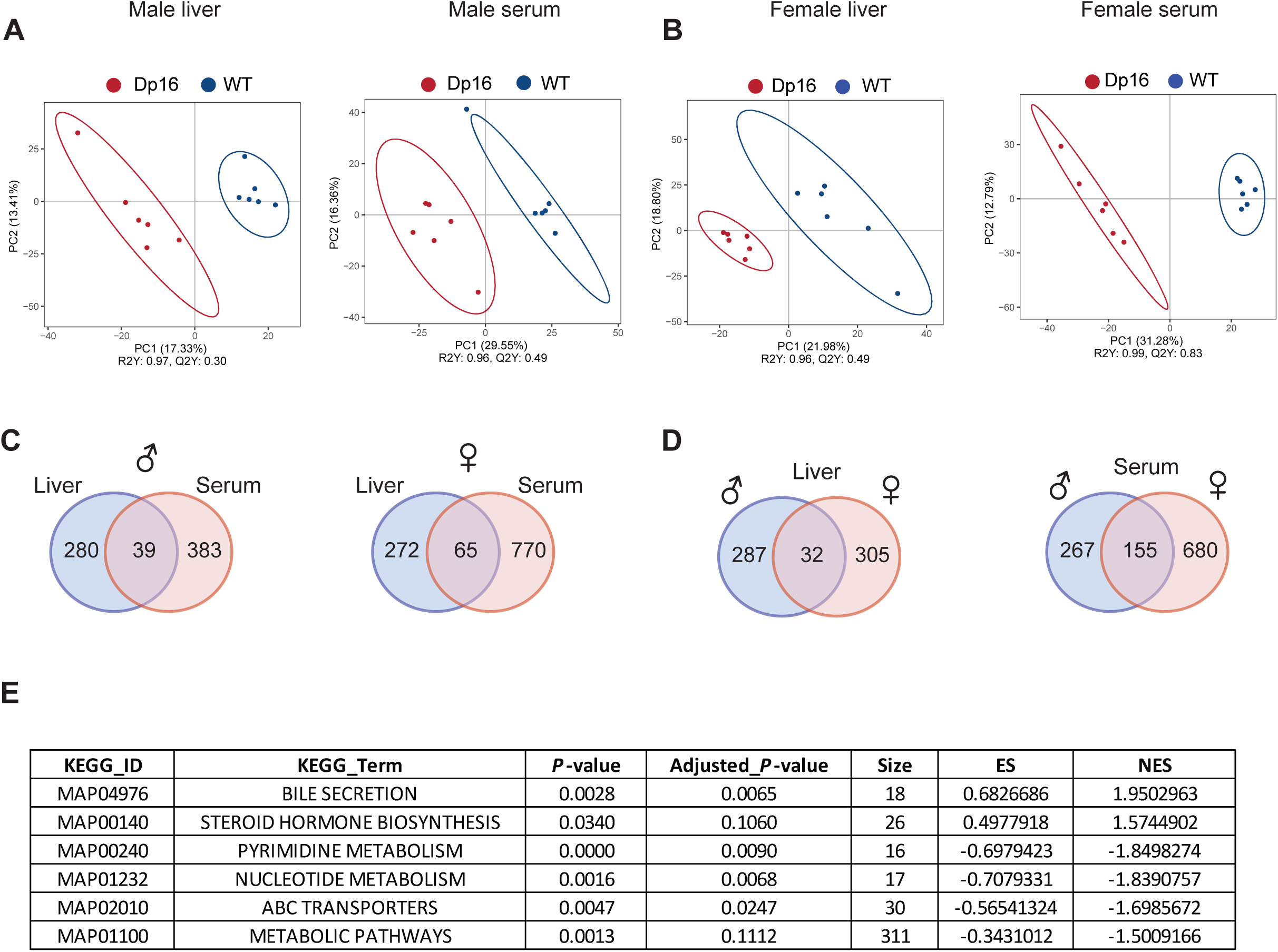
**Altered liver and serum metabolome in Dp16 male and female mice**. (A-B) Partial least squares discrimination analysis (PLS-DA) of liver and serum metabolites of Dp16 and WT males and females. *N* = 6 samples per genotype per sex. (C) Venn diagram of differential metabolites shared between liver and serum in Dp16 male or female mice. (D) Venn diagram of differential liver or serum metabolites shared between Dp16 males and females. (E) KEGG enrichment showing altered metabolic processes in Dp16 female serum. ES, enrichment score; NES, normalized enrichment score.

As shown by KEGG classification, differential metabolites related to global, as well as lipid and amino acid, metabolism accounted for the major differences seen in liver and serum of Dp16 males and females (**Fig. 4 – figure supplement 2 and 3**). In Dp16 female serum, KEGG enrichment analysis indicated altered metabolic pathways related to bile secretion, steroid hormone biosynthesis, nucleotide metabolism, and ABC transporters (**Fig. 4E**). In female and male liver, as well as male serum, KEGG enrichment analyses did not yield any pathways with a false discovery rate less than 0.05.

To provide greater detail, we highlighted some of the differential metabolites found in Dp16 male and female mice (**Table 1 and 2**). Consistent with recent findings (75), we also observed changes in hepatic bile acids content in both sexes, with most of the bile acids showing a reduced level in Dp16 mice. In contrast to the liver, we observed a more extensive changes in circulating bile acids, all of them except two were elevated in Dp16 mice of both sexes. Many bile acids serve as ligands for nuclear hormone receptors (e.g., FXR and TGR5) that control various aspects of glucose and lipid metabolism (76, 77), and extensive changes in circulating bile acids may potentially contribute, at least in part, to the systemic metabolic phenotypes in Dp16 mice. In addition to bile acids, circulating levels of many phospholipids (e.g., LysoPC, LysoPA, LysoPE), some with signaling roles (78), were also altered. In both liver and serum, multiple acyl-carnitines (intermediates in fat oxidation) were elevated in liver and serum, suggesting impaired fatty acid catabolism in Dp16 mice; this phenotype is further supported by our transcriptomic data indicating reduced expression of fat oxidation genes in liver and BAT (data are discussed further below).

**Table 1.**
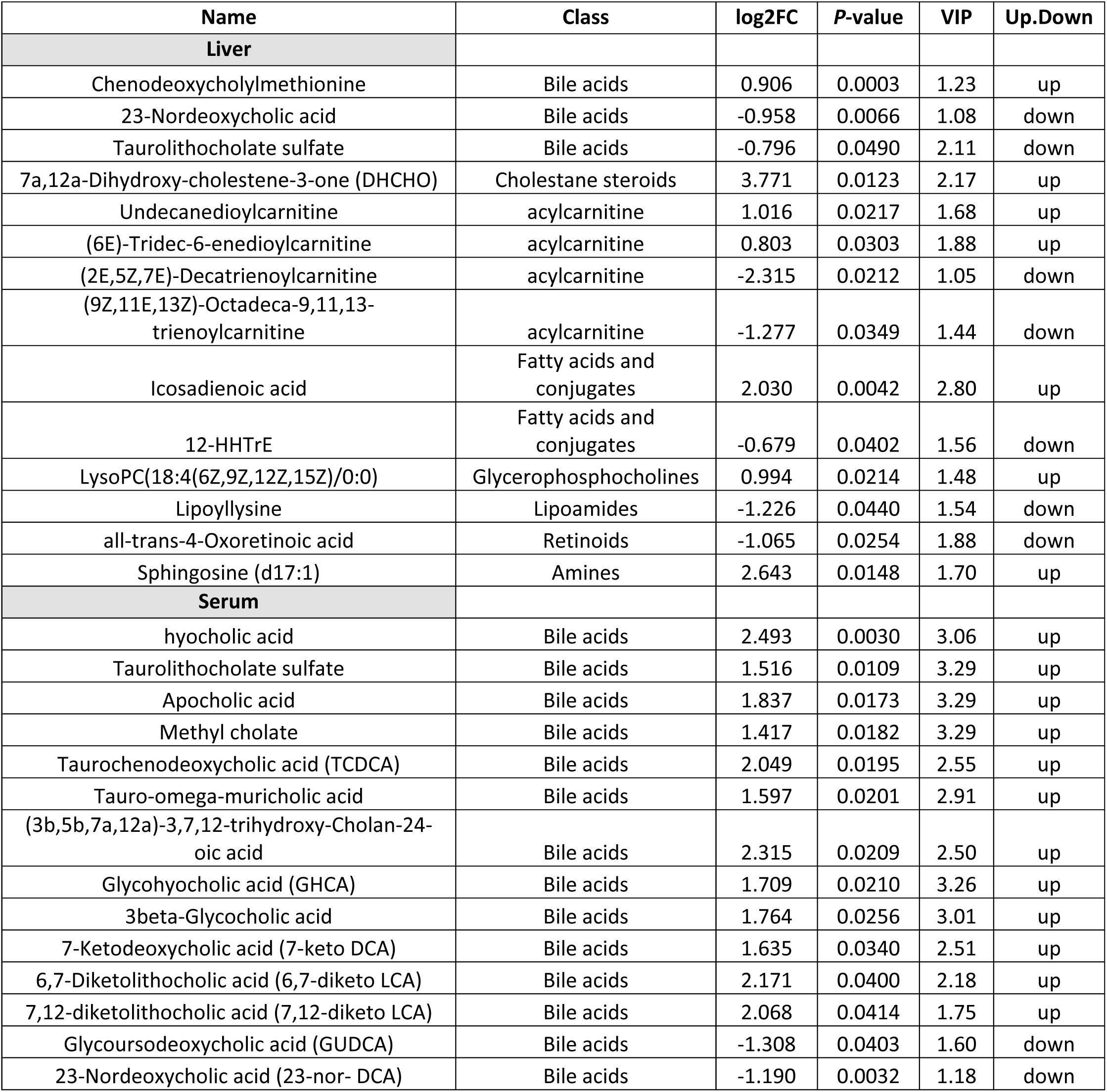
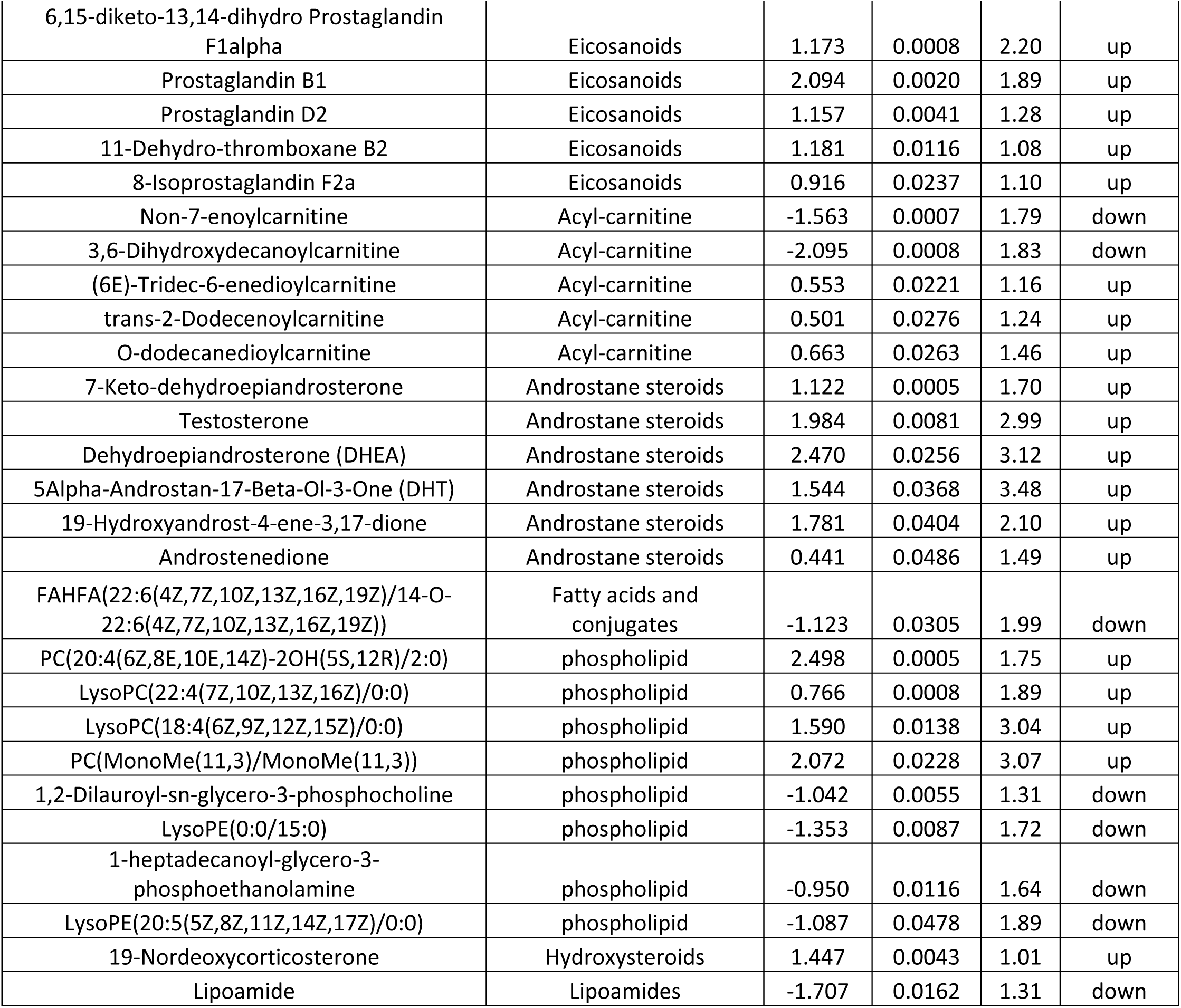
Selective differential metabolites in the liver and serum of Dp16 male mice. . Metabolites are considered significantly different if fold change (FC) > 1.2 or < 0.833, *P*-value < 0.05, and the variable importance in projection (VIP) score is > 1. Sample size: WT (*n* = 6) and Dp16 (*n* = 6)

**Table 2.**
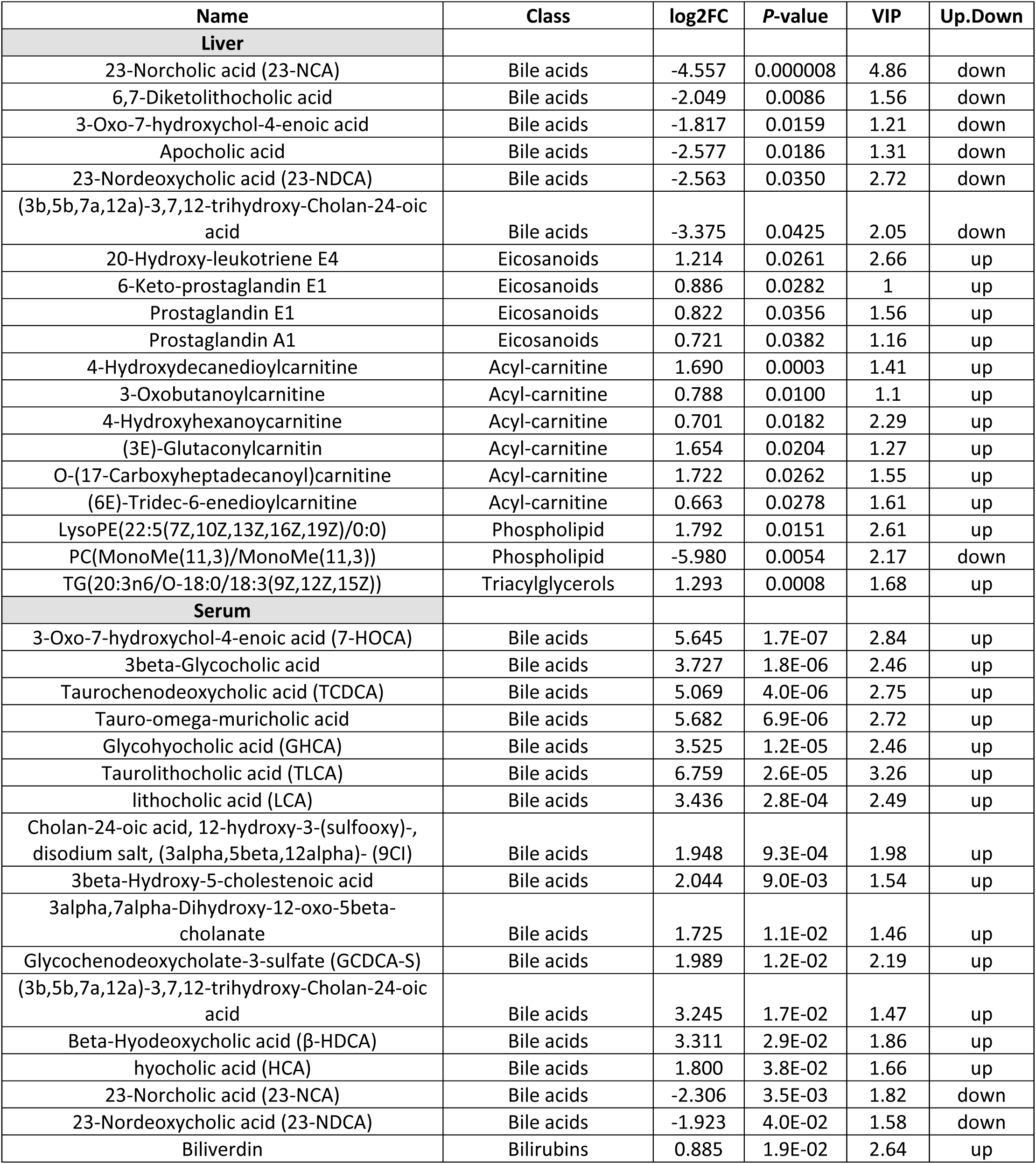
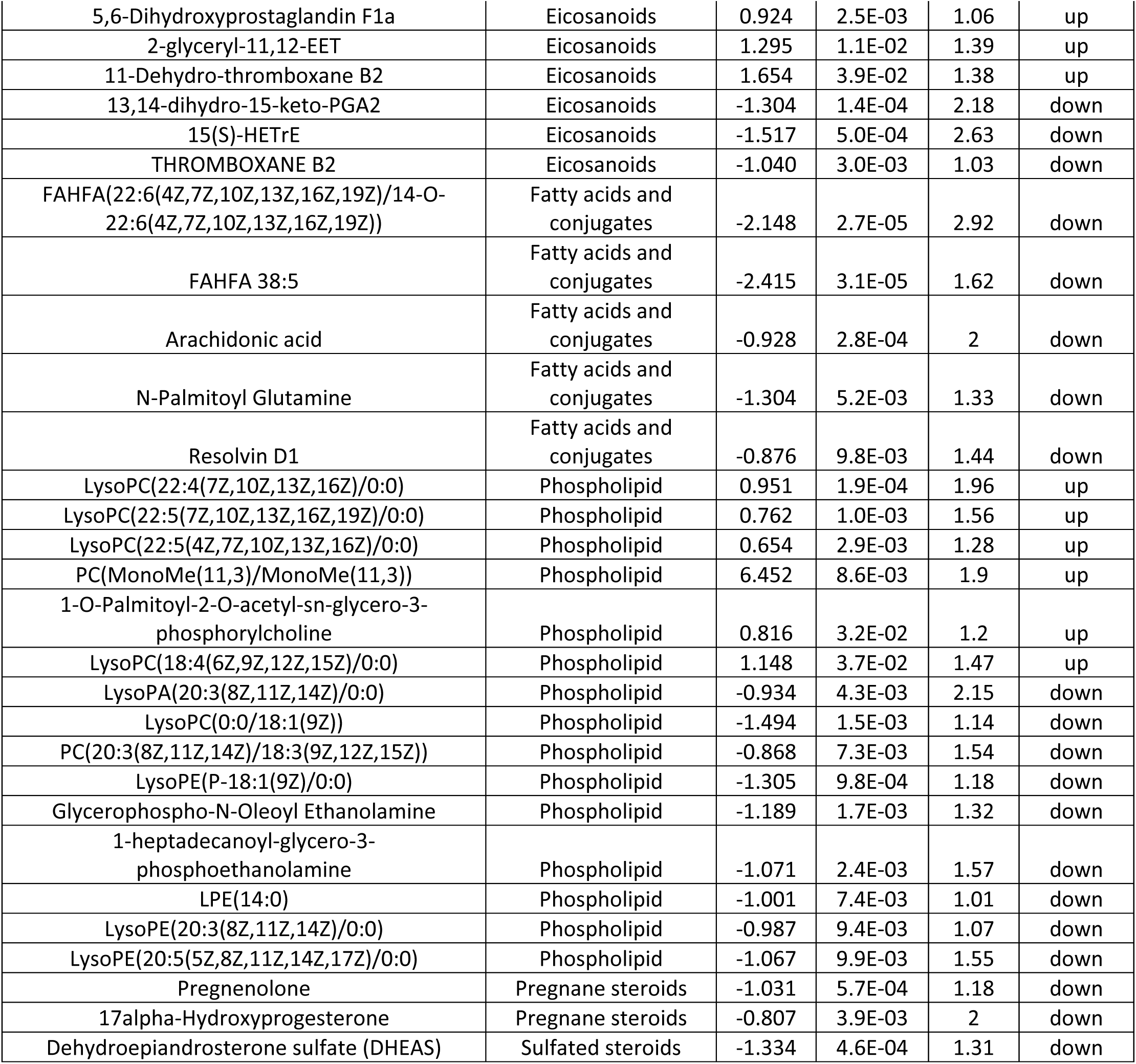
Selective differential metabolites in the liver and serum of Dp16 female mice. . Metabolites are considered significantly different if fold change (FC) > 1.2 or < 0.833, *P*-value < 0.05, and the variable importance in projection (VIP) score is > 1. Sample size: WT (*n* = 6) and Dp16 (*n* = 6)

The levels of multiple eicosanoids, a class of lipids with pro- and anti-inflammatory roles (79), were also changed in Dp16 mice. For example, several pro-inflammatory eicosanoids (e.g., 11-dehydro-thromboxane B2, prostaglandin B1, prostaglandin D2, 20-hydroxy-leukotriene E4) were elevated while some anti-inflammatory eicosanoids (e.g., 13,14-dihydro-15-keto-PGA2, 15(S)-HETrE) were reduced. In addition to eicosanoids, we observed lower levels of several fatty acids with anti-inflammatory and/or anti-diabetic roles (e.g., FAHFA, resolvin D1, all-trans-4-oxoretinoic acid) (80–82) and a concomitant increase in fatty acids with pro-inflammatory roles (e.g., icosadienoic acid, sphingosine) (83, 84). The general pro-inflammatory state in Dp16 mice, reflected by the metabolite data, parallel our transcriptomic data in BAT, liver, muscle, and hypothalamus showing a pro-inflammatory and heightened immune activation state (data are discussed further below).

We also would like to highlight that a crucial biomarker for oxidative stress, 8-Isoprostaglandin F2α (85), was significantly elevated in Dp16 male serum, whereas metabolites (e.g., lipoyllysine and lipoamide) that act as indirect antioxidants (86) to maintain mitochondrial health were reduced. Further, N-palmitoyl glutamine, a recently discovered acylated amino acid that stimulates mitochondrial biogenesis and efficiency (87), was also reduced in Dp16 female serum. Interestingly, 19-Nor-deoxycorticosterone (19-nor-DOC), a powerful mineralocorticoid hormone that increases blood pressure (88, 89), was elevated in Dp16 male serum, raising the possibility of altered blood pressure in these animals. In Dp16 female serum, we also observed elevated biliverdin (precursor of bilirubin), suggesting potential liver injury, which was confirmed by an increase in serum alanine transaminase (ALT) levels (**Figure 4 – figure supplement 4**).

We showed using an ELISA method that estradiol levels are significantly higher in females and testosterone levels in males exhibited high variations (Figure 2 – figure supplement 5). Our metabolite data, however, indicated that six androstane steroids, including testosterone, were significantly elevated in Dp16 male serum, whereas the precursors of female sex hormones (e.g., pregnenolone and 17α-hydroxyprogesterone) were reduced in female serum, presumably due to its greater conversion to estradiol. Given the pleiotropic systemic metabolic effects of testosterone and estradiol (90, 91), changes in circulating sex hormones in Dp16 mice are likely contributing, at least in part, to the sex differences in metabolic phenotypes. Taken together, our metabolomic analyses reveal major sex-dependent and independent changes in liver and serum metabolites likely contributing to the systemic metabolic deficits in Dp16 mice.

### Transcriptomic signatures of immune activation, fibrosis, ER stress, and impaired metabolic processes in Dp16 mice

To address the molecular underpinnings of the metabolic phenotypes seen in chow-fed Dp16 mice, we performed bulk RNA sequencing to assess global changes in the transcriptome and biological pathways across six metabolic tissues (**Fig. 5 – source data 1-24**). Except the liver, females have significantly more differentially expressed genes (DEGs) across tissues compared to males (**Fig. 5A**). In Dp16 females, gWAT, iWAT, and BAT together accounted for the majority of DEGs, with the least number of DEGs seen in skeletal muscle (**Fig. 5A**). In Dp16 males, BAT and liver have the highest number of DEGs, with skeletal muscle having the least DEGs. All tissues except gWAT have more upregulated than downregulated DEGs. Overlap analysis in males and females indicate shared DEGs across tissues, as well as DEGs that are seen only in males or females (**Fig. 5B**). BAT and liver have the highest number of shared DEGs across sex, with skeletal muscle having the least shared DEGs. iWAT, gWAT and BAT have the highest number of DEGs that are female-specific, whereas liver and BAT having the highest number of DEGs that are male-specific (**Fig. 5B**).

**Figure 5.**
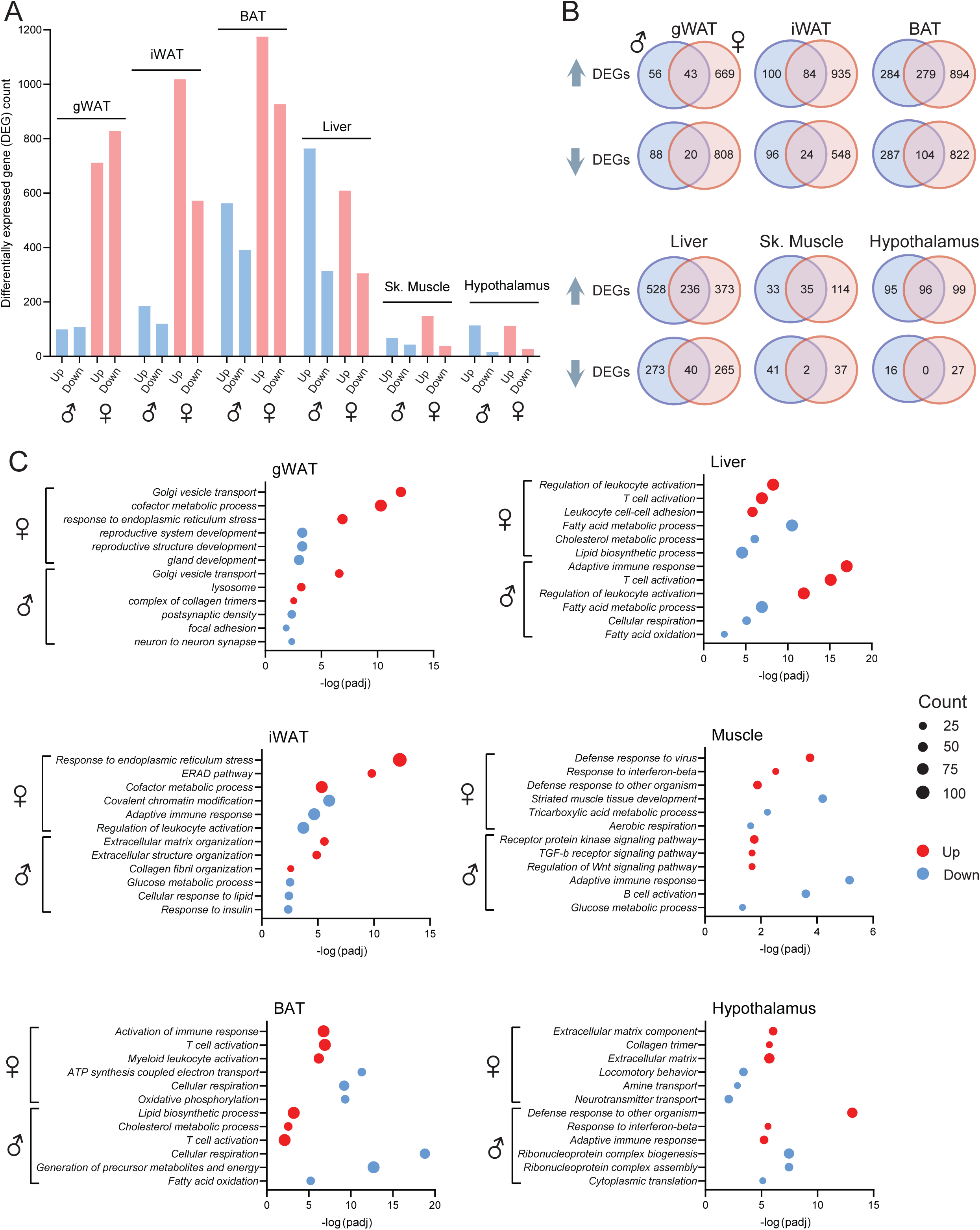
Transcriptomic changes and altered biological pathways across tissues in chow-fed Dp16 male and female mice. **A)** Number of differentially expressed genes (DEGs) that up or down regulated across six tissues in male and female Dp16 mice and their WT littermate controls. DEG is defined as any gene with log2(FC) > 0.5 and padj < 0.05. *N* = 6 per genotype per tissue. gWAT, gonadal white adipose tissue; iWAT, inguinal white adipose tissue; BAT, brown adipose tissue. **B)** Overlap analysis showing DEGs that are shared between males and females, as well as those DEGs found in males or females only, across six tissues. **C)** Gene ontology highlighting some of the top biological pathways altered across six tissues in male and female Dp16 mice.

We performed gene ontology (GO) analysis to reveal which major biological pathways are altered in Dp16 mice that could contribute to their metabolic phenotypes. Among the top biological pathways upregulated in female mice are ER stress (gWAT, iWAT), immune activation (iWAT, BAT, liver, and muscle), fibrosis (iWAT and hypothalamus), and the top downregulated pathways are ATP synthesis and cellular respiration (BAT and skeletal muscle) and lipid metabolism (liver) (**Fig. 5C**). In male mice, some of the top biological pathways upregulated include fibrosis (gWAT and iWAT), immune activation (BAT, liver, and hypothalamus), lipid metabolism (BAT), and receptor signaling (skeletal muscle), and the down-regulated pathways include glucose and lipid metabolism (iWAT, BAT, and liver) and cellular respiration (BAT and liver) (**Fig. 5C**). We highlighted some of the DEGs involved in immune activation, ER stress, fibrosis, glucose and lipid metabolism, fat oxidation, mitochondrial respiration, and signaling in iWAT, BAT, liver, skeletal muscle, and hypothalamus (**Figure 5 – figure supplement 1-5**). These transcriptomic changes parallel our metabolite data (**Table 1 and 2**); both sets of data highlighted impaired lipid metabolism, a pro-inflammatory state, reduced mitochondrial health, and oxidative stress. Altogether, these data underscore major sex differences in tissue transcriptomes, but also revealed common and shared biological pathways affected in Dp16 males and females that underpin their shared metabolic phenotypes. To confirm the biochemical correlates of our RNA-seq data, we examined markers of fibrosis (hydroxyproline) and oxidative stress (malondialdehyde) in liver, gWAT, and iWAT. Collagen content was significantly higher in Dp16 female liver and lower in male gWAT (**Figure 5 – figure supplement 6**). Oxidative stress was significantly higher in Dp16 male liver and lower in gWAT; in females it was lower in gWAT (**Figure 5 – figure supplement 7**). These results partly corroborate our transcriptomic and metabolomic data, and further indicate that fibrosis and oxidative stress occur in Dp16 mice in a sex- and tissue-dependent manner.

### Sexually dimorphic response of Dp16 mice to an obesogenic diet

Since the DS population is prone to developing obesity (11, 13), we challenged the Dp16 mice with an obesogenic high-fat diet (HFD) and assessed how they handle metabolic stress associated with chronic high-fat feeding. There were striking sex differences in the response of Dp16 mice to an HFD. For the first five weeks on HFD, Dp16 males gained a similar amount of weight as the WT controls. From six weeks onward, their body weight diverged, with the WT males gaining significantly more weight compared to Dp16 males (**Fig. 6A**), and this was reflected in much higher adiposity seen in the WT males (**Fig. 6B**). Absolute lean mass was not different between genotypes in males, but % lean mass (when normalized to body weight) was higher in Dp16 males. In contrast to males, Dp16 females gained weight rapidly in the first six weeks on HFD, but by seven weeks onward, the body weights of Dp16 females were no longer significantly different from WT controls (**Fig. 6C**). Body composition analysis revealed no difference in fat mass, but modestly higher lean mass, in Dp16 females after 16 weeks on HFD (**Fig. 6D**).

**Figure 6.**
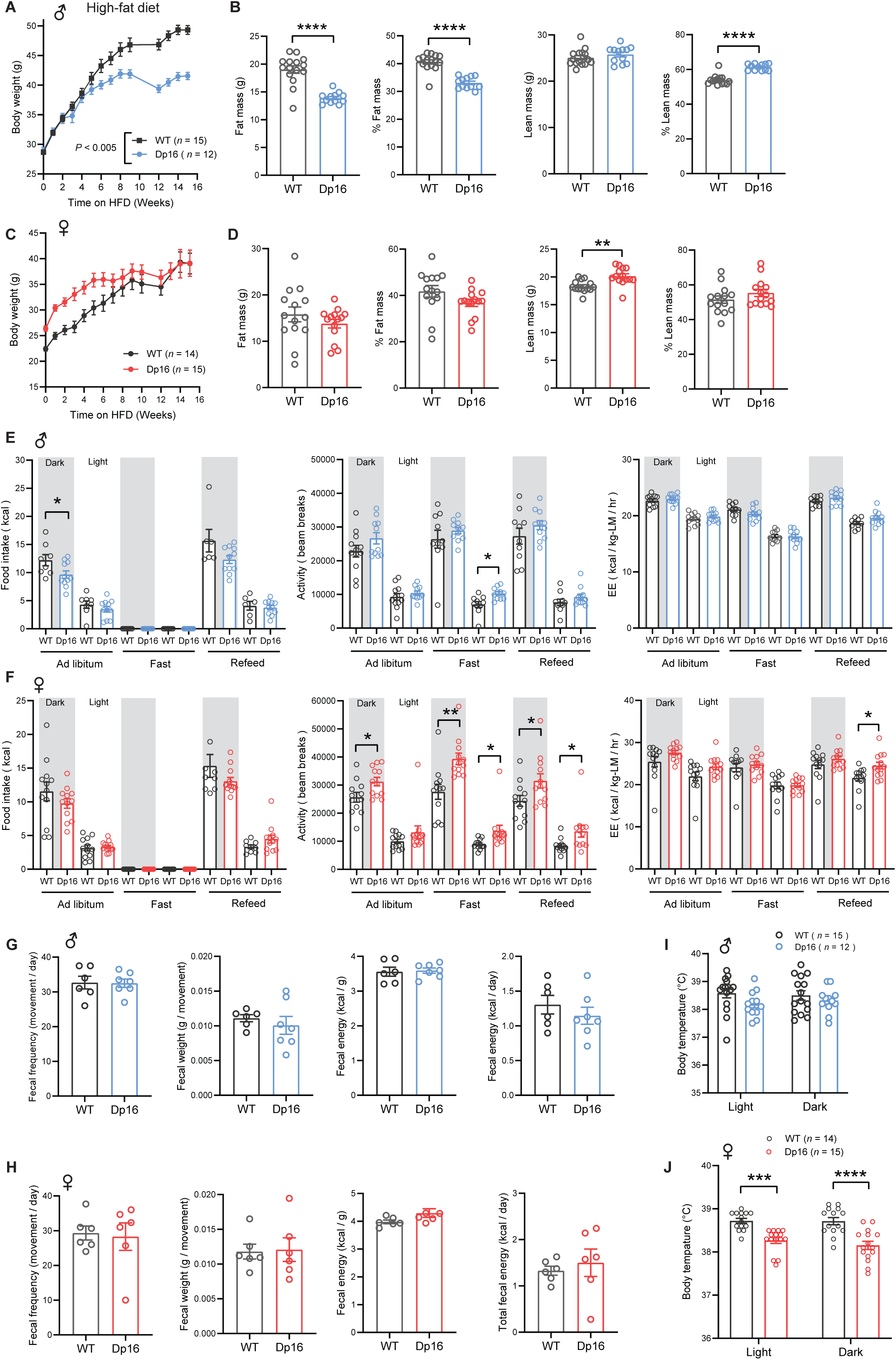
Sexually dimorphism in body weight, body temperature, food intake, and physical activity in Dp16 mice in response to a high-fat diet (HFD). **A)** Body weight of HFD-fed male Dp16 and WT mice over time. **B)** absolute and relative (% of body weight) fat and lean mass in male mice (WT = 15; Dp16 = 12). **C)** Body weight of HFD-fed female Dp16 and WT mice over time. **D)** absolute and relative (% of body weight) fat and lean mass in female mice (WT = 14; Dp16 = 14). **E-F)** Food intake, total physical activity level, and energy expenditure of male (E) and female (F) Dp16 and WT mice across the circadian cycle (light and dark) and metabolic states (*ad libitum* fed, fast, refeed). Sample size for male (WT = 8; Dp16 = 11) and female (WT = 12; Dp16 = 12) mice. **G-H)** fecal frequency, average fecal weight, and fecal energy content (per gram and total) in male (G) and female (H) Dp16 and WT mice on HFD. Sample size for male (WT = 6; Dp16 = 7) and female (WT = 6; Dp16 = 6) mice. **I-J)** Body temperature in the light and dark cycle of male (I) and female (J) Dp16 and WT mice on HFD. Sample size for male (WT = 15; Dp16 = 12) and female (WT = 14; Dp16 = 15) mice. All data are presented as mean ± SEM. * *P*<0.05; *** *P*<0.001; **** *P*<0.0001. For body weight over time, data were analyzed by 2-way ANOVA with Sidek post hoc tests.

We performed indirect calorimetry analysis after the mice were on HFD for 12 weeks. We observed lower *ad libitum* food intake in Dp16 males in the dark/active cycle (**Fig. 6E**), and this could contribute to lower weight gain over time. Dp16 males had modestly higher physical activity levels in the light cycle during fasting, but energy expenditure was not different from WT controls across the circadian cycles and metabolic states (fed, fasted, and refed) (**Fig. 6E**). In contrast to the males, food intake was not different between genotypes in females (**Fig. 6F**). However, Dp16 females had consistently higher physical activity levels in the dark cycle across different metabolic states (fed, fasted, refed). Although energy expenditure (normalized to lean mass) appeared to be slightly higher in Dp16 females, it was only significantly different in the light cycle during the refed period (**Fig. 6F**). To rule out potential overestimation of energy expenditure, we performed ANCOVA analyses (using lean mass as a covariate of energy expenditure) (57). ANCOVA analyses indicated no differences in energy expenditure between genotypes of either sex across the circadian cycles and metabolic states (**Figure 6 – figure supplement 1**).

Next, we determined whether there are any differences in nutrient absorption in the intestine by quantifying fecal output and frequency, as well as fecal energy content in mice on HFD. No differences in any of the fecal parameters were noted between genotypes of either sex (**Fig. 6G-H**). Because we observed differences in the body temperature of chow-fed mice, we again measured the body temperature of mice on HFD. In contrast to higher body temperature of chow-fed males, Dp16 males on HFD appeared to have slightly lower body temperature in the light cycle, though not significant (**Fig. 6I**). Like the chow-fed females, Dp16 females on HFD also had lower body temperature in both the light and dark cycles (**Fig. 6J**). We measured serum triiodothyronine (T3) to determine whether altered thyroid hormone level contributes to lower body temperature. Contrary to expectation, both Dp16 males and females had elevated serum T3 levels (**Figure 6 – figure supplement 2**). We also measured serum levels of sex and stress hormones, as these could contribute to our phenotypes. Corticosterone levels were not different between genotypes of either sex. Testosterone levels in males appeared lower but not significant; estradiol levels, however, were higher in Dp16 females (**Figure 6 – figure supplement 2**), possibly contributing to their lower body temperature and higher physical activity (71–74).

At termination of study, we assessed whether there are differences in tissue weight in Dp16 mice on HFD. Tissues were collected from male mice at 50 weeks of age when they were on HFD for 34.5 weeks. At the time of termination, body weights and tissue weights of gWAT, iWAT, and liver were significantly lower in Dp16 males relative to WT controls (Figure 6 – figure supplement 3). Although the absolute weights of heart and kidney were not different between genotypes, the relative weights (% of body weight) of heart and kidney were significantly higher in Dp16 males. For females, tissues were collected at 45 weeks of age (on HFD for 26 weeks). Although the body weights of Dp16 females on HFD were not different from WT controls at 35 weeks of age (on HFD for 16 weeks), there were significantly lower at 45 weeks of age (on HFD for 26 weeks) (**Figure 6 – figure supplement 3**). The absolute and relative weights of gWAT and iWAT were significantly lower in Dp16 females. The absolute and relative weights of heart and kidney, however, were higher in Dp16 females.

We also noted that a marker of fibrosis (hydroxyproline content) was significantly higher in Dp16 male liver, as well as in Dp16 female liver, gWAT, and iWAT. A marker of oxidative stress (malondialdehyde content) was not significant different between genotypes of either sex in liver, gWAT, and iWAT (**Figure 6 – figure supplement 4**). Taken together, these data indicate major sex differences in the physiological response of Dp16 mice to an obesogenic diet.

### High-fat diet exacerbates glucose intolerance and insulin resistance in Dp16 mice

Since Dp16 mice on a standard chow diet developed overt insulin resistance and dyslipidemia, we determined whether HFD would further exacerbate these phenotypes. In overnight (16 h) fasted males, serum insulin, triglyceride, cholesterol, NEFA, and β-hydroxybutyrate (ketone) levels were not different between genotypes (**Fig. 7A**), even though Dp16 males had significantly lower body weight and adiposity. Fasting blood glucose levels, however, were lower in Dp16 males, likely reflecting lower adiposity. The Dp16 females appeared to have higher fasting insulin levels, though not significant (**Fig. 7B**). Fasting blood glucose and serum triglyceride and cholesterol levels were not different between genotypes in female mice. Serum NEFA and β-hydroxybutyrate (ketone) levels were significantly lower in Dp16 females (**Fig. 7B**), suggesting lower fasting-induced lipolysis and hepatic fat oxidation.

**Figure 7.**
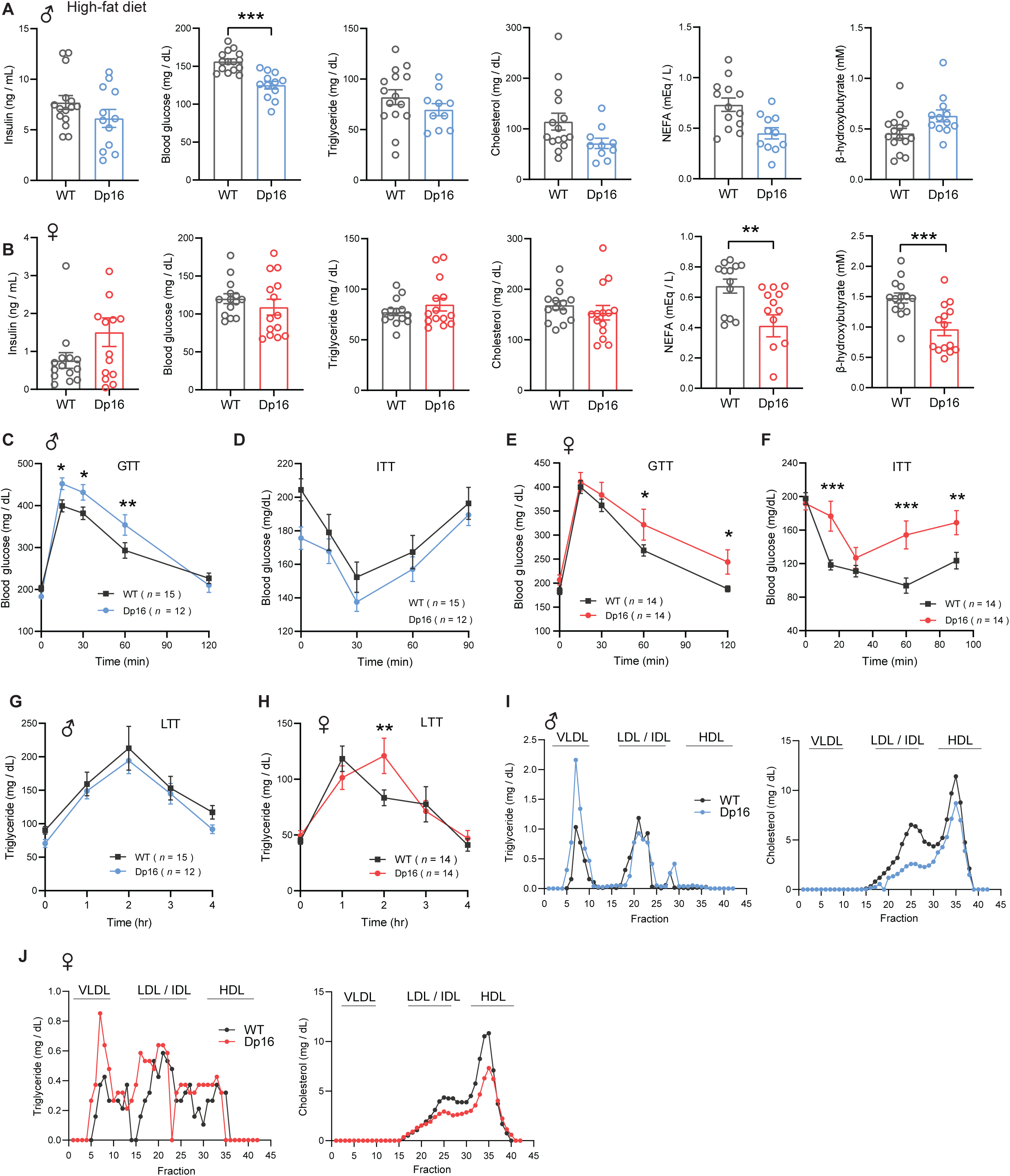
Exacerbated glucose intolerance and insulin resistance in Dp16 mice fed a high-fat diet (HFD). A-B) Overnight fasting insulin, blood glucose, serum triglyceride, cholesterol, non-esterified free fatty acids (NEFA), and β-hydroxybutyrate (ketone) in male (A) and female (B) Dp16 and WT mice on HFD. Sample size for male mice (WT = 15; Dp16 = 12) and female mice (WT = 14; Dp16 = 14). **C-F)** Exacerbated glucose intolerance as determined by the glucose tolerance test (GTT) in male (C) and female (E) Dp16 compared to WT controls on HFD. Exacerbated insulin resistance as determined by the insulin tolerance test (ITT) in male (D) and female (F) Dp16 compared to WT controls. Sample size for male mice (WT = 15; Dp16 = 12) and female mice (WT = 14; Dp16 = 14). **G-H)** The rate of triglyceride clearance in response to lipid gavage as determined by the lipid tolerance test (LTT) in male (G) and female (H) Dp16 and WT mice. Sample size for male mice (WT = 15; Dp16 = 12) and female mice (WT = 14; Dp16 = 14). **I-J)** Pooled mouse sera from male (I) and female (J) Dp16 and WT mice were fractionated by fast protein liquid chromatography (FPLC), and the triglyceride and cholesterol content of each fraction was quantified. Fractions corresponding to very-low density lipoprotein (VLDL), low-density lipoprotein (LDL), intermediate-density lipoprotein (IDL), and high-density lipoprotein (HDL) are indicated. All data are presented as mean ± SEM. * *P*<0.05; ** *P*<0.01; *** *P*<0.001; **** *P*<0.0001. For all tolerance tests, data were analyzed by 2-way ANOVA with Sidek post hoc tests.

Next, we subjected Dp16 mice on HFD to glucose tolerance test. Both Dp16 males and females showed exacerbated glucose intolerance compared to WT controls (**Fig. 7C and E**). Direct assessment of insulin sensitivity showed that Dp16 males and females have significantly reduced glucose clearance in response to insulin injection compared to WT controls, with Dp16 females exhibiting a more pronounced insulin resistance phenotype (**Fig. 7D and F**).

Given that Dp16 mice on a standard chow diet had marked deficit in lipid clearance, we again performed lipid tolerance tests to assess whether Dp16 mice on HFD have worsening lipid handling capacity. No differences were observed between genotypes in male mice, whereas Dp16 females had a modest impairment in lipid clearance compared to WT controls (**Fig. 7G-H**). Analysis of lipoprotein profiles showed that both Dp16 males and females have higher VLDL-TG and lower LDL-C compared to WT controls, with the effect more pronounced in males (**Fig. 7I-J**). Taken together, these data indicate that despite divergent weight gain in response to HFD, both Dp16 males and females show exacerbated insulin resistance and dyslipidemia.

## DISCUSSION

In this study, we show that triplication of Hsa21 gene orthologs in the Dp16 mouse model causes profound disruption in systemic metabolism. By combining deep phenotyping with multi-omics approaches, we show that gene dosage imbalance contributes to major changes in transcriptomes, metabolomes, and biological pathways that link to systemic insulin resistance, glucose intolerance, impaired lipid clearance, dyslipidemia, and exacerbate metabolic stress induced by an obesogenic diet. Our findings provide valuable insights and plausible mechanistic explanations for the prevalence of obesity, dyslipidemia, and diabetes in the DS population.

We show that most triplicated Hsa21 gene orthologs are expressed at the expected ∼1.5-fold or higher across six metabolic tissues, but in a striking sex- and tissue-specific manner. Variegated overexpression of Hsa21 genes has been previously noted in individuals with DS (92). In Dp16 mice, female adipose depots exhibit the highest number of differentially expressed triplicated genes, and this is associated with the substantial weight gain observed in Dp16 females. Males, in contrast, have relatively stable body weight and adiposity under standard chow diet despite having similar gene dosage imbalance. These sex differences underscore the complex interactions between triplicated genes and sex-dependent regulatory mechanisms that influences tissue transcriptomes (93), fat mass expansion and systemic energy metabolism (90, 91). Our data reinforces sex as an important biological determinant of metabolic risk in DS, consistent with documented sex differences in the susceptibility of individuals with DS to developing obesity and metabolic impairments (13, 23, 94–97).

Although their weight trajectories diverge, both male and female Dp16 mice exhibit hallmark features of metabolic dysfunction, including fasting hyperinsulinemia, glucose intolerance, insulin resistance, and impaired lipid clearance. These core phenotypes shared between the sexes arise in spite of divergent adiposity, indicating that dysregulated systemic metabolism is a primary consequence of triplicated gene dosage imbalance rather than a secondary effect of increased fat mass. The insulin resistance phenotype, as well as impaired triglyceride clearance and alterations in lipoprotein profiles are consistent with the propensity of individuals with DS to developing type 2 diabetes, dyslipidemia and altered fasting lipid profiles (10–19). These parallels strengthen the translational relevance of the Dp16 model. Future studies are needed to determine which aspects of lipid metabolism—hepatic lipid export, adipose tissue lipolysis, lipoprotein turnover—are dysregulated in DS. Nevertheless, our assessments of mitochondrial function and biochemical markers of fibrosis and oxidative stress, as well as our transcriptomic, metabolomic, and pathway enrichments analyses have provided some mechanistic insights. Our data point to a combination of changes—impaired glucose and lipid metabolism, mitochondrial function, ER and oxidative stress, fibrosis, and low-grade inflammation—all of which are known drivers of adverse metabolic outcomes (98–100). These collective changes act in concert across major metabolic tissues to disrupt local and systemic energy metabolism in Dp16 mice.

Our metabolomic analyses highlighted extensive remodeling of the metabolome of liver and serum. Strikingly, the majority of differential metabolites are not shared between the sexes. This once again underscores the unexpected and complex interactions of triplicated genes and biological sex in determining phenotypic outcomes. The differential metabolites are clustered around pathways related to lipid, amino acid, and bile acid metabolism; some of these pathways have been previously documented in DS (75, 101). Despite sex differences, we noted that several classes of metabolites—bile acids, acylcarnitine, eicosanoids, phospholipids, fatty acid conjugates—are shared by Dp16 males and females and that the directionality of change is also broadly consistent between the sexes. Altered bile acid pools, including both primary and conjugated bile acids made in the liver (e.g., taurochenodeoxycholic acid, chenodeoxycholymethionine, hyocholic acid, glycohyocholic acid) and secondary bile acids made by gut bacteria (e.g., lithocholic acid, taurolithocholate sulfate, glycoursodeoxycholid acid, keto and diketo lithocholic acid), suggest potential changes in gut–liver axis and bile-acid–regulated metabolic signaling (77, 102, 103) that can contribute to metabolic dysfunction. The directionality of change in multiple immuno-regulatory eicosanoids (79) and fatty acids (80–82) appears to promote a pro-inflammatory state in Dp16 mice, a phenotype that is also supported by our multi-tissue transcriptomic data. Interestingly, several hepatic and serum phospholipid species (e.g., Lyo-PC, Lyso-PA, Lyso-PA) are either elevated or reduced in Dp16 mice. Given the complex signaling roles for some of these phospholipids (78), we speculate that these changes may underline some aspects of the metabolic deficits in Dp16 mice. Thus, remodeling of tissue and serum metabolomes, in combination with major changes in transcriptomes across tissues, likely contributed to the pronounced metabolic dysfunction in Dp16 mice.

The introduction of an obesogenic diet helped us to reveal the complex interplay of gene and environment in the context of Hsa21 gene dosage imbalance. This becomes relevant as individuals with DS live increasingly longer lives, and can have varied lifestyles and diet. Despite opposite weight trajectories on HFD—males gaining less and females initially gaining more—both Dp16 male and female mice develop worsened glucose intolerance and insulin resistance. This dissociation between weight gain and metabolic impairment again point to gene dosage imbalance as the primary cause of metabolic dysfunction rather than a secondary effect of altered adiposity. The increased fibrosis observed in adipose tissue and liver in Dp16 mice on HFD suggests that chronic nutritional stress exacerbates extracellular matrix remodeling, a change in tissue architecture that is known to compromise adipose tissue and liver function (99, 104).

Although not the focus of the present study, we unexpectedly discovered that an obesogenic diet causes heart enlargement in Dp16 mice; this phenotype was not observed in Dp16 mice fed a standard chow. The cause of heart enlargement in response to a high-fat diet is presently unknown. Given that congenital heart defect is frequently seen in DS (105), prior studies in DS mouse models have been focused on the contribution of triplicated genes to developmental heart defects (e.g., atrial or ventricular septal defect) (69, 106). Our data suggests that the complex interactions of genetics and diet may predispose adult individuals with DS to cardiovascular complications independent of congenital heart abnormalities.

Among the 115 Hsa21 triplicated gene orthologs located on the syntenic region of mouse chromosome 16, several are known to affect metabolism, oxidative stress, inflammatory response, and/or fibrosis. These include *Dyrk1a* (107), *Dscr1*/*Rcan1* (108, 109), *AtpJ* (110), *Atp5o* (111), *Nrip1* (112), *Tiam1* (113, 114), *Prdm15* (115), *Ripk4* (116), *Znf295* (117), *Hmgn1* (118), *Cbr1* (119), *Bach1* (120), *Fam3b* (121), *Ets2* (122), *Adamts5* (123, 124), *Usp16* (125), *Runx1* (126), *Sim2* (127), and the interferon receptor gene locus (*Ifnar2*, *Il10rb*, *Ifnar1*, *Ifngr2*) (128, 129). Interestingly, some of these triplicated genes show sexually dimorphic expression in gWAT (*Il10rb*, *Cbr1*, *Ets2*, *Nrip1*), iWAT *(Il10rb*, *Runx1*, *Adamts5*, *Tiam1*, *Rcan1*, *Ripk4*, *Bach1*, *Nrip1*, *Dyrk1a*, *Fam3b*), BAT (*Cbr1*, *Usp16*, *Ets2*, *Atp5o*, *Nrip1*), Liver (*Tiam1*, *Rcan1*, *Bach1*, *Prdm15*), skeletal muscle (*Adamts5*, *Ifnar2*, *Rcan1*, *Ets2*, *Atp5o*, *Nrip1*), and hypothalamus (*Ripk4*, *Sim2*); these sex-biased expression patterns may contribute to the sex differences in Dp16 metabolic phenotypes. In striking contrast to other DS phenotypes, normalizing the expression dosage of the interferon receptor gene locus does not reverse the hepatic lipid metabolism profile (75). None of the other triplicated candidate genes were studied in the context of trisomy; thus, it is unknown whether normalizing these triplicated genes—individually or in combination—could reverse some or all of the metabolic phenotypes described. Systematic genetic dissection of dosage-sensitive genes in the context of trisomy—an approach successfully used for other DS phenotypes (69, 130–132)—are required to establish their necessity and sufficiency in promoting metabolic dysfunction. Given the complex metabolic phenotypes of Dp16, we anticipate that multiple dosage-sensitive genes are likely to work additively or synergistically to disrupt systemic glucose and lipid metabolism.

Several limitations of the study, however, are noted. First, while Dp16 mouse model contains ∼58% of the Hsa21 gene orthologs (55, 69), it does not contain the full complement of Hsa21 gene orthologs. Although no single mouse model fully recapitulates all DS phenotypes (133), in light of the impact and complex combinatorial effects of the triplicated genes on phenotypic outcomes and tissue transcriptomes (6, 134), it is worthwhile in future studies to comprehensively examine the metabolic phenotypes of the triple compound model (Dp16;Dp10;Dp17) carrying the full complement of the Hsa21 gene orthologs (135). The large-scale breeding, cost, and labor associated with generating the compound mice have been a major challenge; however, progress has recently been made to overcome this hurdle (136). Secondly, Dp16 is a segmental duplication model and not a trisomic model with an independently segregating chromosome. Recent studies have suggested that the presence of an extra chromosome (i.e., trisomy) can affect phenotypes and disomic gene expressions beyond the gene dosage effect of triplicated genes (137). Disentangling dosage-dependent versus chromosome-dependent effects will be an important future direction. Thirdly, although we observed major perturbations across tissue transcriptomes, not all changes in mRNAs would translate into corresponding changes in protein levels, and vice versa (138, 139). Future studies incorporating proteomics data would enhance and complement our transcriptomic results. Fourthly, while our studies were ongoing, Tolu and co-workers also reported the glucose intolerance and insulin resistance phenotype in Dp16 mice (140). The authors also showed reduced insulin content in the pancreatic β-cell without changes in β-cell mass. Notably, in our study, we did not measure pancreatic insulin content. We also did not include transcriptomic data from the pancreas, as none of the RNA samples passed the quality control needed for RNA sequencing.

In summary, our data show that the triplication of Hsa21 gene orthologs severely disrupts metabolic homeostasis through concerted perturbations in transcriptome, metabolome, tissue remodeling, mitochondrial function, and glucose and lipid metabolism. Our findings provide physiological contexts for ongoing studies aiming to understand the vulnerability of DS population to developing metabolic disorders. The striking and extensive sex differences uncovered here argue that metabolic studies and therapeutic strategies in DS should account for sex as an important biological variable. Our study highlighted both shared and sex-specific mechanisms of metabolic dysfunction in a DS mouse model, and the impact of gene dosage imbalance on altering whole-body metabolism. The wealth of molecular, biochemical, and physiological data help lay the crucial groundwork for genetic dissection of dosage-sensitive genes causally linked to metabolic dysfunction, and to inform efforts at identifying actionable therapeutic targets that can mitigate one or more aspects of metabolic deficits seen in individuals with DS.

## AUTHOR CONTRIBUTIONS

GWW: Conceptualization; FC, MS, DCS, SA, CMN, SA, MMS, GWW: Formal analysis; GWW, MMS: Funding acquisition; FC, MS, CMN, DCS, SA, GWW: Investigation; MMS, YEY: Methodology; GWW: Project administration; GWW, MMS: Supervision; FC, CMN, DCS, MMS, GWW: Visualization; GWW: Roles/Writing - original draft; FC, MS, DCS, CMN, SA, YEY, MMS: Writing - review & editing.

## Supporting information

All supplementary figures 1-20

## ACKNOWLEDGEMENTS

The work was funded, in part, by grants from the National Institute of Health (DK084171 to GWW, HL138193 and DK130640 to MMS, HD109750 and DC019735 to YEY). MS and DCS were supported by an NIH T32 training grant (HL007534). The FPLC/serum analyses were conducted by the Mouse metabolic Phenotyping Center (MMPC) at Baylor College of medicine, funded by NIH grants DK114356 and UM1HG006348. The fecal bomb calorimetry analysis was performed at the University of Michigan Animal Phenotyping Core, supported by center grants 1U2CDK135066-01 (Mi-MPMOD) and DK020572 (MDRC). The Metabolomics Workbench/National Metabolomics Data Repository (NMDR) is supported by the NIH (U2C-DK119886), Common Fund Data Ecosystem (CFDE) (3OT2OD030544) and Metabolomics Consortium Coordinating Center (M3C) (1U2C-DK119889)

## DATA AVAILABILITY

All RNA-seq data have been deposited in NCBI Sequence Read Archive (SRA), with the accession # PRJNA1160420. The metabolomics data is available at the NIH Common Fund’s National Metabolomics Data Repository (NMDR) website, the Metabolomics Workbench (https://www.metabolomicsworkbench.org) where it has been assigned Study ID (ST004905, ST004915, ST004916, and ST004918). The data can be accessed directly via it’s Project DOI: http://dx.doi.org/10.21228/M8SZ8Q).

## CONCLICT OF INTEREST

We declared that none of the authors has conflict of interest.

## SUPPLEMENTAL FIGURE FILES LEGENDS AND SOURCE DATA

Figure 2 **– figure supplement 1. Body and tissue weights of chow-fed male and female mice at termination of study.** Tissues were collected from chow-fed male and female mice at 27.5 weeks of age. Body weights and the absolute (A and C) and relative (B and D; % of body weight) weights of gWAT, iWAT, liver, and kidney in Dp16 and WT male (A-B) and female (C-D) mice. gWAT, gonadal white adipose tissue; iWAT, inguinal white adipose tissue. Sample size: WT male = 10; Dp16 male = 30; WT female = 15; Dp16 female = 10. All data are presented as mean ± SEM. * *P*<0.05; ** *P*<0.01; *** *P*<0.001.

Figure 2 – **figure supplement 2. ANCOVA analysis of energy expenditure in chow-fed mice where lean mass is used as a covariate.** ANCOVA analysis of WT and Dp16 male mice across the circadian cycle (dark and light) in *ad libitum* fed (A), fasted (B), and refed (C) states. ANCOVA analysis of WT and Dp16 female mice across the circadian cycle (dark and light) in *ad libitum* fed (D), fasted (E), and refed (F) states. Male Sample size: WT = 12; Dp16 = 12. Female sample size: WT = 15; Dp16 = 6.

Figure 2 – **figure supplement 3. Reduced mitochondrial activity in the brown adipose tissue (BAT) of Dp16 mice.** Mitochondrial respiration through complex I (CI), CII, and CIV in BAT of WT and Dp16 male and female mice fed a standard chow. (**A and D)** Average oxygen consumption rate (OCR) traces per group, normalized to mitochondrial content. Each group tracing represents the average trace of 10 WT and 9-10 Dp16 samples. Each tracing shows the entire process of the Seahorse-based respirometry assay with injection compounds listed at the time of introduction to the sample. The sequence is as follows: i) basal reads, ii) addition of NADH (activation of respiration through complex I), iii) addition of antimycin A (AA, inhibitor of complex III) and rotenone (Rot, inhibitor of complex I), iv) addition of TMPD and ascorbate (to activate complex IV via electron donation to cytochrome c), and finally v) addition of azide (inhibitor of complex IV). **(B and E)** The same information as presented in (A and D) conducted on the same samples, but the NADH injection step is replaced with the injection of succinate (to activate respiration through complex II) and rotenone (to inhibit complex I). **(C and F)** Average values of all data presented for BAT. Each data point represents the average of three technical replicates measured at three separate times. Both independent measurements of complex IV (CIV) were used to determine average CIV respiration. **** *p* < 0.0001 (two-way ANOVA with Sidak’s multiple comparison).

Figure 2 – **figure supplement 4. Serum Triiodothyronine (T3), sex and stress hormone levels in WT and Dp16 mice fed a standard chow.** (A) Serum T3 levels in male and female mice. (B) Serum testosterone levels in male mice. (C) Serum estradiol levels in female mice. (D) Serum corticosterone in male and female mice. Sample size: male WT = 8-10; male Dp16 = 25-30; female WT = 14-15; female Dp16 = 10.

Figure 3 – **figure supplement 1. Liver triacylglycerol (TAG), diacylglycerol (DAG), and cholesterol levels in chow-fed Dp16 mice.** Quantification of hepatic TAG and DAG (by TLC method), and cholesterol (by infinity assay kit) levels in chow-fed Dp16 male (A-C) and female mice (D-F) and their corresponding WT controls. Sample size: male WT = 10 and Dp16 = 30; female WT = 15 and Dp16 = 10.

Figure 3 – **figure supplement 2. Mitochondrial activity in the liver of Dp16 mice.** Mitochondrial respiration through complex I (CI), CII, and CIV in the liver of WT and Dp16 male and female mice fed a standard chow. (**A and D)** Average oxygen consumption rate (OCR) traces per group, normalized to mitochondrial content. Each group tracing represents the average trace of 10 WT and 10 Dp16 samples.

Each tracing shows the entire process of the Seahorse-based respirometry assay with injection compounds listed at the time of introduction to the sample. The sequence is as follows: i) basal reads, ii) addition of NADH (activation of respiration through complex I), iii) addition of antimycin A (AA, inhibitor of complex III) and rotenone (Rot, inhibitor of complex I), iv) addition of TMPD and ascorbate (to activate complex IV via electron donation to cytochrome c), and finally v) addition of azide (inhibitor of complex IV). **(B and E)** The same information as presented in (A and D) conducted on the same samples, but the NADH injection step is replaced with the injection of succinate (to activate respiration through complex II) and rotenone (to inhibit complex I). **(C and F)** Average values of all data presented for liver. Each data point represents the average of three technical replicates measured at three separate times. Both independent measurements of complex IV (CIV) were used to determine average CIV respiration.

Figure 4 – **Source data 1.** Supplemental table of differential metabolites in Dp16 male mouse liver vs WT controls. Differential metabolites criteria: VIP > 1.0, fold change (FC) > 1.2 or FC < 0.833 and *P*-value < 0.05. Sample name notation: male WT liver (M_WT_L), male WT serum (M_WT_S), male Dp16 liver (M_16_L), male Dp16 serum (M_16_S), female WT liver (F_WT_L), female WT serum (F_WT_L), female Dp16 liver (F_16_L), female Dp16 serum (F_16_S).

Figure 4 – **Source data 2.** Supplemental table of differential metabolites in Dp16 female mouse liver vs WT controls. Differential metabolites criteria: VIP > 1.0, fold change (FC) > 1.2 or FC < 0.833 and *P*-value < 0.05. Sample name notation: male WT liver (M_WT_L), male WT serum (M_WT_S), male Dp16 liver (M_16_L), male Dp16 serum (M_16_S), female WT liver (F_WT_L), female WT serum (F_WT_L), female Dp16 liver (F_16_L), female Dp16 serum (F_16_S).

Figure 4 – **Source data 3.** Supplemental table of differential metabolites in Dp16 male mouse serum vs WT controls. Differential metabolites criteria: VIP > 1.0, fold change (FC) > 1.2 or FC < 0.833 and *P*-value < 0.05. Sample name notation: male WT liver (M_WT_L), male WT serum (M_WT_S), male Dp16 liver (M_16_L), male Dp16 serum (M_16_S), female WT liver (F_WT_L), female WT serum (F_WT_L), female Dp16 liver (F_16_L), female Dp16 serum (F_16_S).

Figure 4 – **Source data 4.** Supplemental table of differentially expressed metabolites in Dp16 female mouse serum vs WT controls. Differential metabolites criteria: VIP > 1.0, fold change (FC) > 1.2 or FC < 0.833 and *P*-value < 0.05. Sample name notation: male WT liver (M_WT_L), male WT serum (M_WT_S), male Dp16 liver (M_16_L), male Dp16 serum (M_16_S), female WT liver (F_WT_L), female WT serum (F_WT_L), female Dp16 liver (F_16_L), female Dp16 serum (F_16_S).

Figure 4 – **Source data 5.** Supplemental table showing the shared and distinct differential metabolites in Dp16 male and female mouse liver and serum vs WT controls. Sample name notation: male WT liver (M_WT_L), male WT serum (M_WT_S), male Dp16 liver (M_16_L), male Dp16 serum (M_16_S), female WT liver (F_WT_L), female WT serum (F_WT_L), female Dp16 liver (F_16_L), female Dp16 serum (F_16_S).

Figure 4 – f**igure supplement 1. Differential metabolites found in the liver and serum of Dp16 male and female mice.** Volcano plots showing differential metabolites up- and down-regulated in Dp16 male liver (A), female liver (B), male serum (C), and female serum (D). *n* = 6 per genotype. VIP, Variable Importance in Projection. VIP scores provide a quantitative measure of a metabolite’s discriminatory power between different groups. Metabolites with a VIP score of 1.0 or greater are considered significant.

Figure 4 – **figure supplement 2. KEGG classification analysis of liver metabolites.** KEGG classification plots based on the differential metabolites from male (A) and female (B) Dp16 mouse liver vs WT control. The horizontal coordinates in the graph indicate the number of metabolites annotated under a particular KEGG pathway as a percentage of the number of all annotated metabolites, the vertical coordinates are KEGG pathway primary classifications on the right and KEGG pathway secondary classifications on the left.

Figure 4 – **figure supplement 3. KEGG classification analysis of serum metabolites.** KEGG classification plots based on the differential metabolites from male (A) and female (B) Dp16 mouse liver vs WT control. The horizontal coordinates in the graph indicate the number of metabolites annotated under a particular KEGG pathway as a percentage of the number of all annotated metabolites, the vertical coordinates are KEGG pathway primary classifications on the right and KEGG pathway secondary classifications on the left.

Figure 4 – f**igure supplement 4. Serum alanine aminotransferase (ALT) levels in WT and Dp16 mice fed a standard chow.** Serum ALT levels in male and female mice. Sample size: male WT = 10; male Dp16 = 27; female WT = 14; female Dp16 = 10.

Figure 5 **– Source data 1.** Supplemental table with differentially expressed genes (DEGs) upregulated in the gonadal white adipose tissue (gWAT) of chow-fed Dp16 male mice relative to WT controls.

Figure 5 – **Source data 2.** Supplemental table with differentially expressed genes (DEGs) down-regulated in the gonadal white adipose tissue (gWAT) of chow-fed Dp16 male mice relative to WT controls.

Figure 5 **– Source data 3.** Supplemental table with differentially expressed genes (DEGs) upregulated in the inguinal white adipose tissue (iWAT) of chow-fed Dp16 male mice relative to WT controls.

Figure 5 – **Source data 4.** Supplemental table with differentially expressed genes (DEGs) down-regulated in the inguinal white adipose tissue (iWAT) of chow-fed Dp16 male mice relative to WT controls.

Figure 5 – **Source data 5.** Supplemental table with differentially expressed genes (DEGs) upregulated in the brown adipose tissue (BAT) of chow-fed Dp16 male mice relative to WT controls.

Figure 5 – **Source data 6.** Supplemental table with differentially expressed genes (DEGs) down-regulated in the brown adipose tissue (BAT) of chow-fed Dp16 male mice relative to WT controls.

Figure 5 **– Source data 7.** Supplemental table with differentially expressed genes (DEGs) upregulated in the liver of chow-fed Dp16 male mice relative to WT controls.

Figure 5 – **Source data 8.** Supplemental table with differentially expressed genes (DEGs) down-regulated in the liver of chow-fed Dp16 male mice relative to WT controls.

Figure 5 – **Source data 9.** Supplemental table with differentially expressed genes (DEGs) upregulated in the skeletal muscle (gastrocnemius) of chow-fed Dp16 male mice relative to WT controls.

Figure 5 – S**ource data 10.** Supplemental table with differentially expressed genes (DEGs) down-regulated in the skeletal muscle (gastrocnemius) of chow-fed Dp16 male mice relative to WT controls.

Figure 5 – **Source data 11.** Supplemental table with differentially expressed genes (DEGs) upregulated in the hypothalamus of chow-fed Dp16 male mice relative to WT controls.

Figure 5 – **Source data 12.** Supplemental table with differentially expressed genes (DEGs) down-regulated in the hypothalamus of chow-fed Dp16 male mice relative to WT controls.

Figure 5 **– Source data 13.** Supplemental table with differentially expressed genes (DEGs) upregulated in the gonadal white adipose tissue (gWAT) of chow-fed Dp16 female mice relative to WT controls.

Figure 5 – **Source data 14.** Supplemental table with differentially expressed genes (DEGs) down-regulated in the gonadal white adipose tissue (gWAT) of chow-fed Dp16 female mice relative to WT controls.

Figure 5 – **Source data 15.** Supplemental table with differentially expressed genes (DEGs) upregulated in the inguinal white adipose tissue (iWAT) of chow-fed Dp16 female mice relative to WT controls.

Figure 5 – **Source data 16.** Supplemental table with differentially expressed genes (DEGs) down-regulated in the inguinal white adipose tissue (iWAT) of chow-fed Dp16 female mice relative to WT controls.

Figure 5 **– Source data 17.** Supplemental table with differentially expressed genes (DEGs) upregulated in the brown adipose tissue (BAT) of chow-fed Dp16 female mice relative to WT controls.

Figure 5 **– Source data 18.** Supplemental table with differentially expressed genes (DEGs) down-regulated in the brown adipose tissue (BAT) of chow-fed Dp16 female mice relative to WT controls.

Figure 5 – **Source data 19.** Supplemental table with differentially expressed genes (DEGs) upregulated in the liver of chow-fed Dp16 female mice relative to WT controls.

Figure 5 – **Source data 20.** Supplemental table with differentially expressed genes (DEGs) down-regulated in the liver of chow-fed Dp16 female mice relative to WT controls.

Figure 5 – **Source data 21.** Supplemental table with differentially expressed genes (DEGs) upregulated in the skeletal muscle (gastrocnemius) of chow-fed Dp16 female mice relative to WT controls.

Figure 5 – **Source data 22.** Supplemental table with differentially expressed genes (DEGs) down-regulated in the skeletal muscle (gastrocnemius) of chow-fed Dp16 female mice relative to WT controls.

Figure 5 – **Source data 23.** Supplemental table with differentially expressed genes (DEGs) upregulated in the hypothalamus of chow-fed Dp16 female mice relative to WT controls.

Figure 5 – **Source data 24.** Supplemental table with differentially expressed genes (DEGs) down-regulated in the hypothalamus of chow-fed Dp16 female mice relative to WT controls.

Figure 5 – **figure supplement 1.** Differentially expressed genes (DEGs) involved in ER stress, fibrosis, glucose and lipid metabolism that are up- or down-regulated in the inguinal white adipose tissue (iWAT) of Dp16 mice.

Figure 5 – **figure supplement 2.** Differentially expressed genes (DEGs) involved in immune activation, lipid metabolism, and mitochondrial respiration that are up- or down-regulated in the brown adipose tissue (BAT) of Dp16 mice.

Figure 5 – **figure supplement 3.** Differentially expressed genes (DEGs) involved in immune activation, lipid metabolism, and mitochondrial respiration that are up- or down-regulated in the liver of Dp16 mice.

Figure 5 – **figure supplement 4.** Differentially expressed genes (DEGs) involved in immune response, metabolism, mitochondrial respiration, and Wnt signaling that are up- or down-regulated in the skeletal muscle (gastrocnemius) of Dp16 mice.

Figure 5 – **figure supplement 5.** Differentially expressed genes (DEGs) involved in immune response and extracellular matrix that are upregulated in the hypothalamus of Dp16 mice.

Figure 5 – **figure supplement 6. Hydroxyproline (marker of fibrosis) and malondialdehyde (marker of oxidative stress) levels in the liver, gWAT, and iWAT of chow-fed Dp16 mice. (A-F)** Quantification of hydroxyproline content in the liver, gWAT, and iWAT of Dp16 male (A-C) and female (D-F) mice and their corresponding WT controls. gWAT, gonadal white adipose tissue; iWAT, inguinal white adipose tissue. Sample size: male WT = 7-10 and Dp16 = 27-29; female WT = 10-13 and Dp16 = 7-10. **(G-L)** Quantification of malondialdehyde (MDA) levels in the liver, gWAT, and iWAT of Dp16 male (G-I) and female (J-L) mice and their corresponding WT controls. gWAT, gonadal white adipose tissue; iWAT, inguinal white adipose tissue. Sample size: male WT = 6-10 and Dp16 = 25-30; female WT = 8-13 and Dp16 = 6-10. All data are presented as mean ± SEM. * *P*<0.05; ** *P*<0.01

Figure 6 – **figure supplement 1. ANCOVA analysis of energy expenditure in HFD-fed mice where lean mass is used as a covariate.** ANCOVA analysis of WT and Dp16 male mice across the circadian cycle (dark and light) in *ad libitum* fed (A), fasted (B), and refed (C) states. ANCOVA analysis of WT and Dp16 female mice across the circadian cycle (dark and light) in *ad libitum* fed (D), fasted (E), and refed (F) states. Male Sample size: WT = 12; Dp16 = 12. Female sample size: WT = 14; Dp16 = 14.

Figure 6 – f**igure supplement 2. Serum Triiodothyronine (T3), sex and stress hormone levels in WT and Dp16 mice fed a high-fat diet.** (A) Serum T3 levels in male and female mice. (B) Serum testosterone levels in male mice. (C) Serum estradiol levels in female mice. (D) Serum corticosterone in male and female mice. Sample size: male WT = 9-13; male Dp16 = 11-12; female WT = 14-15; female Dp16 = 14.

Figure 6 – **figure supplement 3. Body and tissue weights of HFD-fed male and female mice at termination of study.** Tissues were collected from male mice (50 weeks old) after they had been fed a high-fat diet for 34.5 weeks. Body weights and the absolute (A) and relative (B; % of body weight) weights of gWAT, iWAT, liver, and kidney in Dp16 and WT male mice. Female tissues (45 weeks old) were mice had been fed a high-fat diet for 26 weeks. Body weights and the absolute (A) and relative (B; % of body weight) weights of gWAT, iWAT, liver, heart, and kidney in Dp16 and WT female mice. gWAT, gonadal white adipose tissue; iWAT, inguinal white adipose tissue. Sample size: WT male = 14; Dp16 male = 12; WT female = 14; Dp16 female = 12. All data are presented as mean ± SEM. * *P*<0.05; ** *P*<0.01; *** *P*<0.001; **** *P*<0.0001.

Figure 6 – **figure supplement 4. Hydroxyproline (marker of fibrosis) and nalondialdehyde (marker of oxidative stress) levels in the liver, gWAT, and iWAT of Dp16 mice on HFD. (A-F)** Quantification of hydroxyproline content in the liver, gWAT, and iWAT of Dp16 male (A-C) and female (D-F) mice and their corresponding WT controls. gWAT, gonadal white adipose tissue; iWAT, inguinal white adipose tissue. Sample size: male WT = 13-14 and Dp16 = 10-12; female WT = 10-14 and Dp16 = 12-14. **(G-L)** Quantification of malondialdehyde (MDA) levels in the liver, gWAT, and iWAT of Dp16 male (G-I) and female (J-L) mice and their corresponding WT controls. gWAT, gonadal white adipose tissue; iWAT, inguinal white adipose tissue. Sample size: male WT = 13-14 and Dp16 = 12; female WT = 14 and Dp16 = 13-14. All data are presented as mean ± SEM. * *P*<0.05; ** *P*<0.01

