## Supplementary material for "Gene dosage imbalance disrupts systemic metabolism in the Dp16 Down syndrome mouse model": All supplementary figures 1-20

Fig. 2 - figure supplement 1

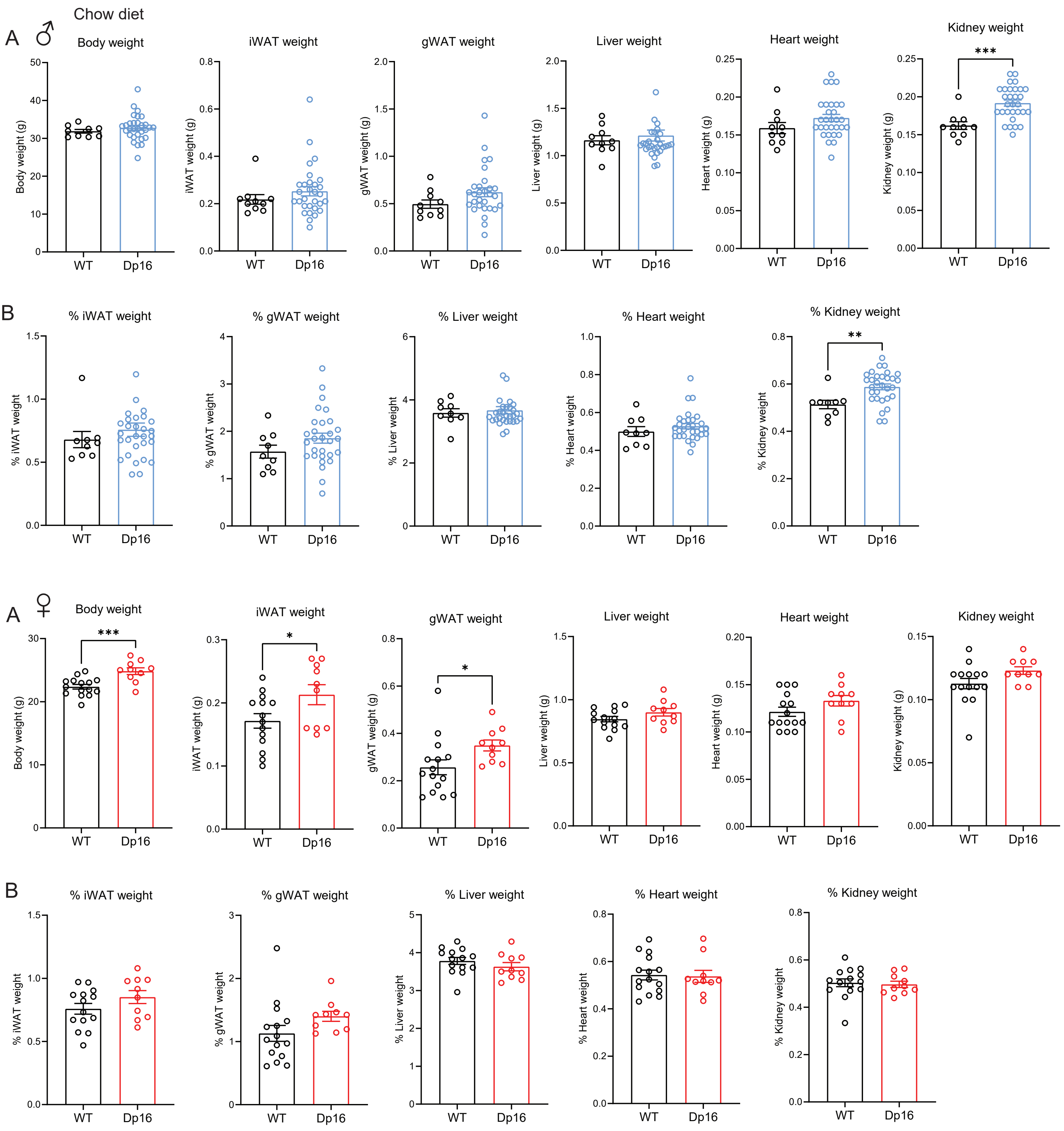

Fig. 2 - figure supplement 2

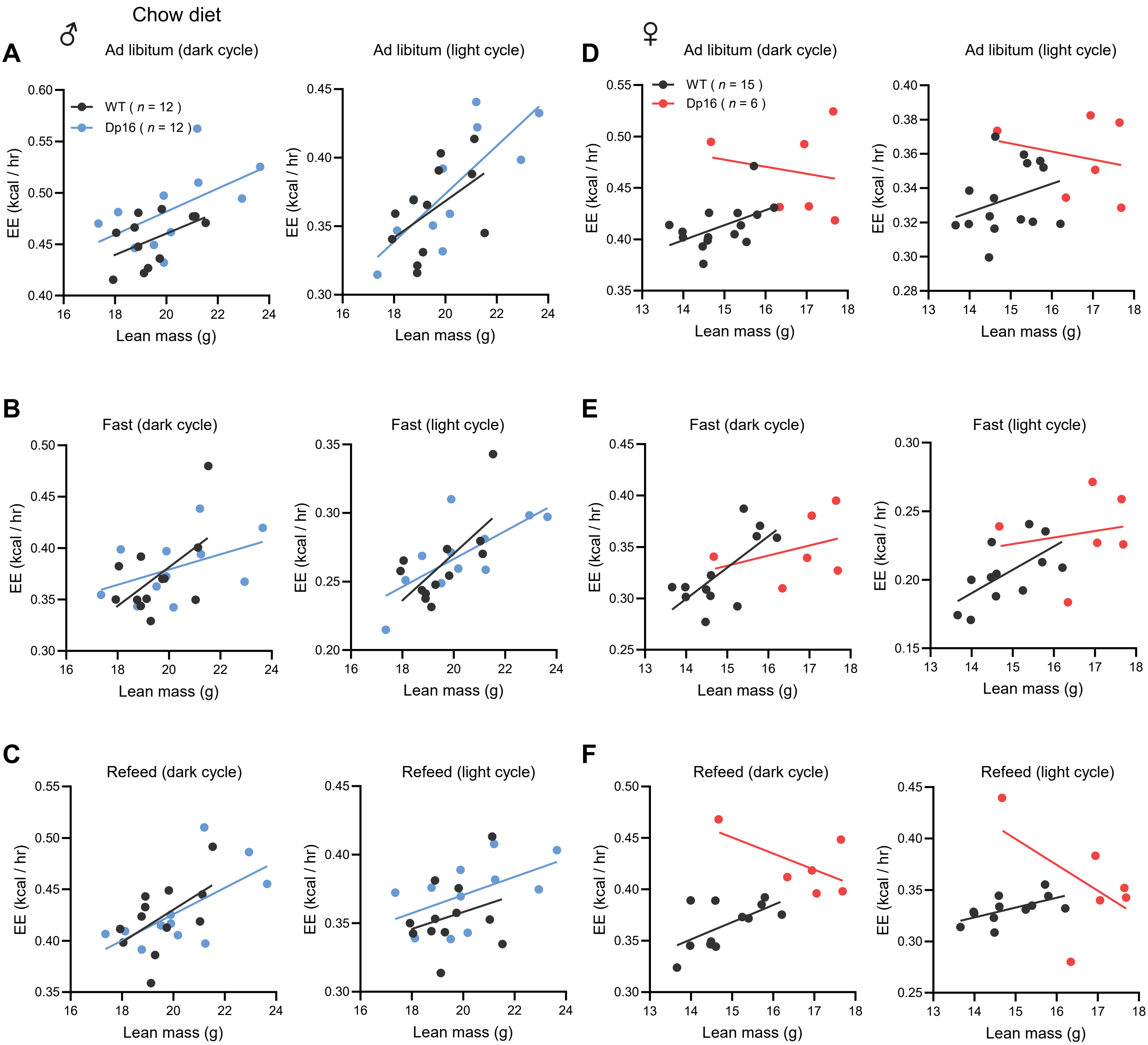

Fig. 2 - figure supplement 3

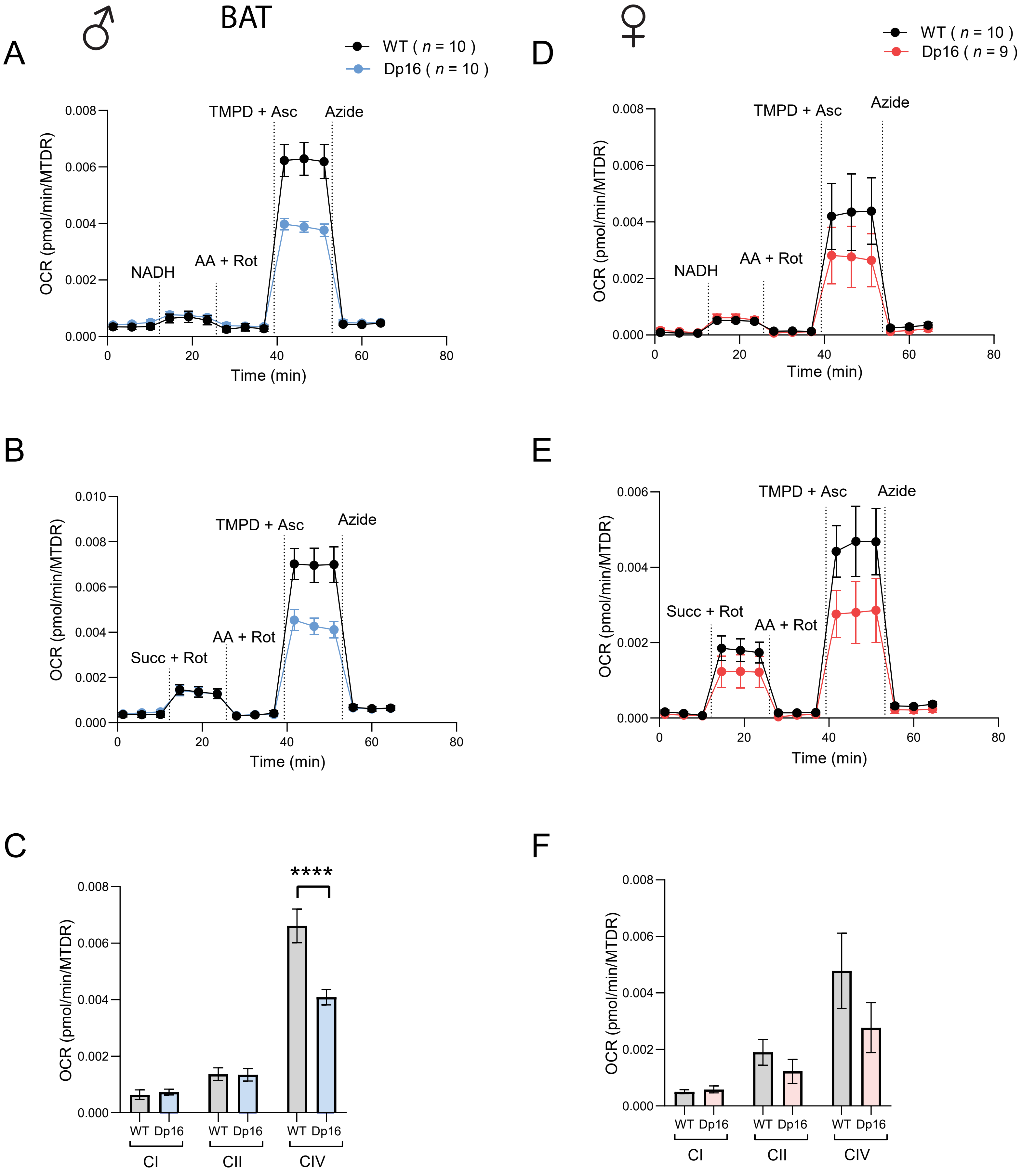

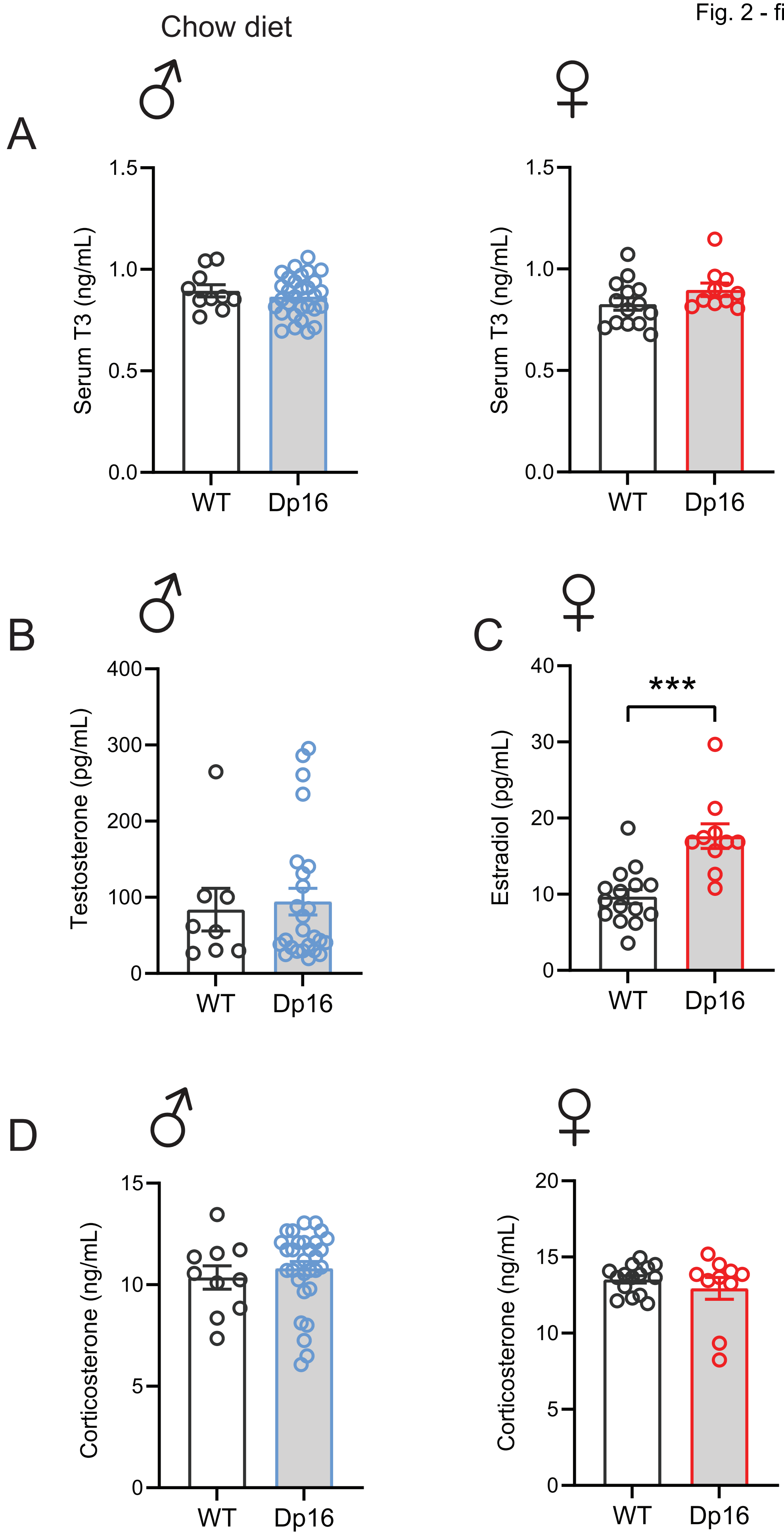
Chow diet
♂
♀
A
Serum T3 (ng/mL)
1.5
1.0
0.5
0.0
WT
Dp16
B
Testosterone (pg/mL)
400
300
200
100
0
C
Estradiol (pg/mL)
40
30
20
10
***
D
Corticosterone (ng/mL)
15
5
20

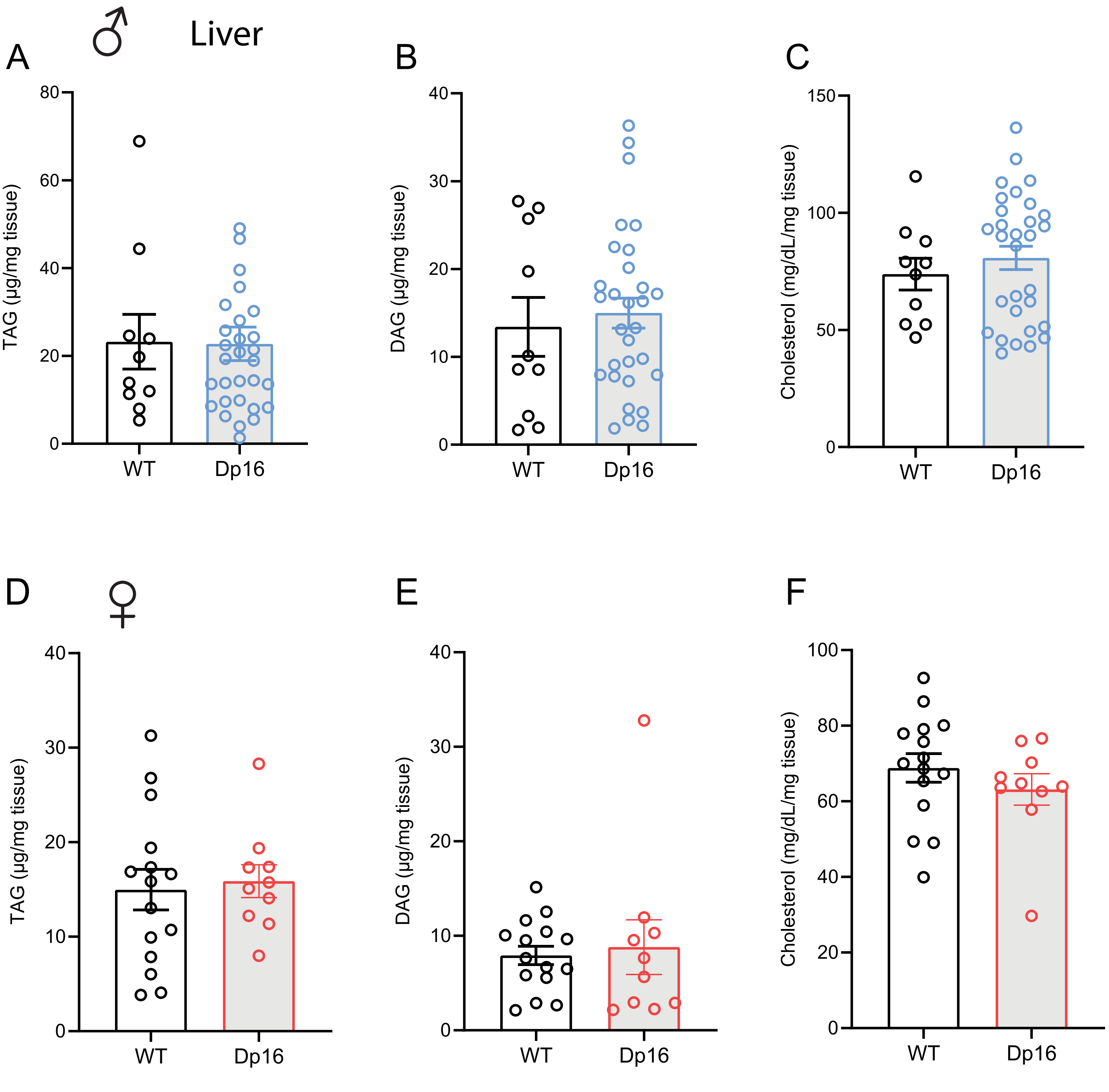

♂
Liver
A
B
C
D
♀
E
F
TAG (µg/mg tissue)
DAG (µg/mg tissue)
Cholesterol (mg/dL/mg tissue)
0
20
40
60
80
10
30
50
100
150
WT
Dp16

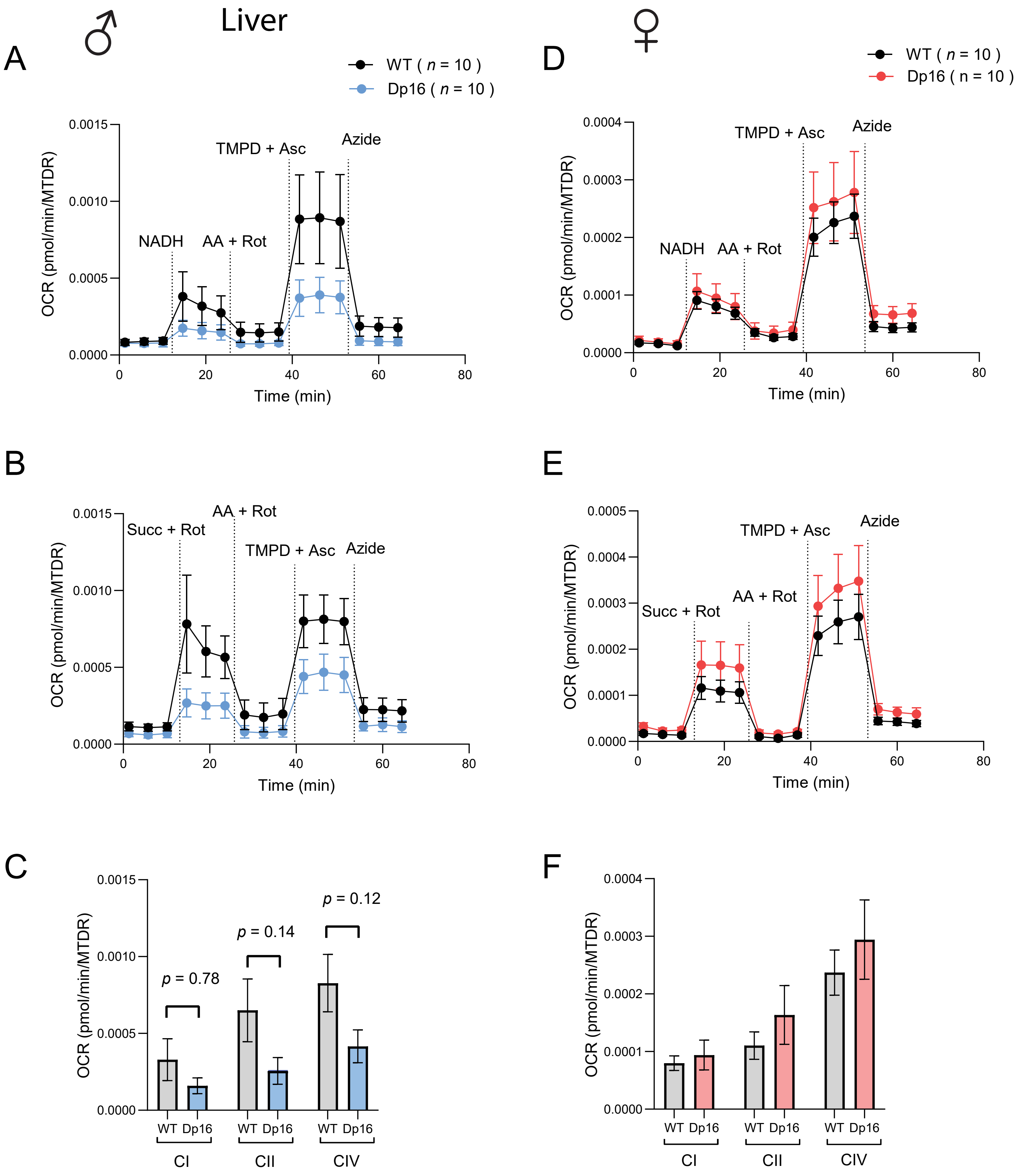
Liver
♂
♀
A
WT ( n = 10 )
Dp16 ( n = 10 )
OCR (pmol/min/MTDR)
Time (min)
NADH
AA + Rot
TMPD + Asc
Azide
B
Succ + Rot
C
p = 0.78
p = 0.14
p = 0.12
WT Dp16
CI
CII
CIV
D
E
F

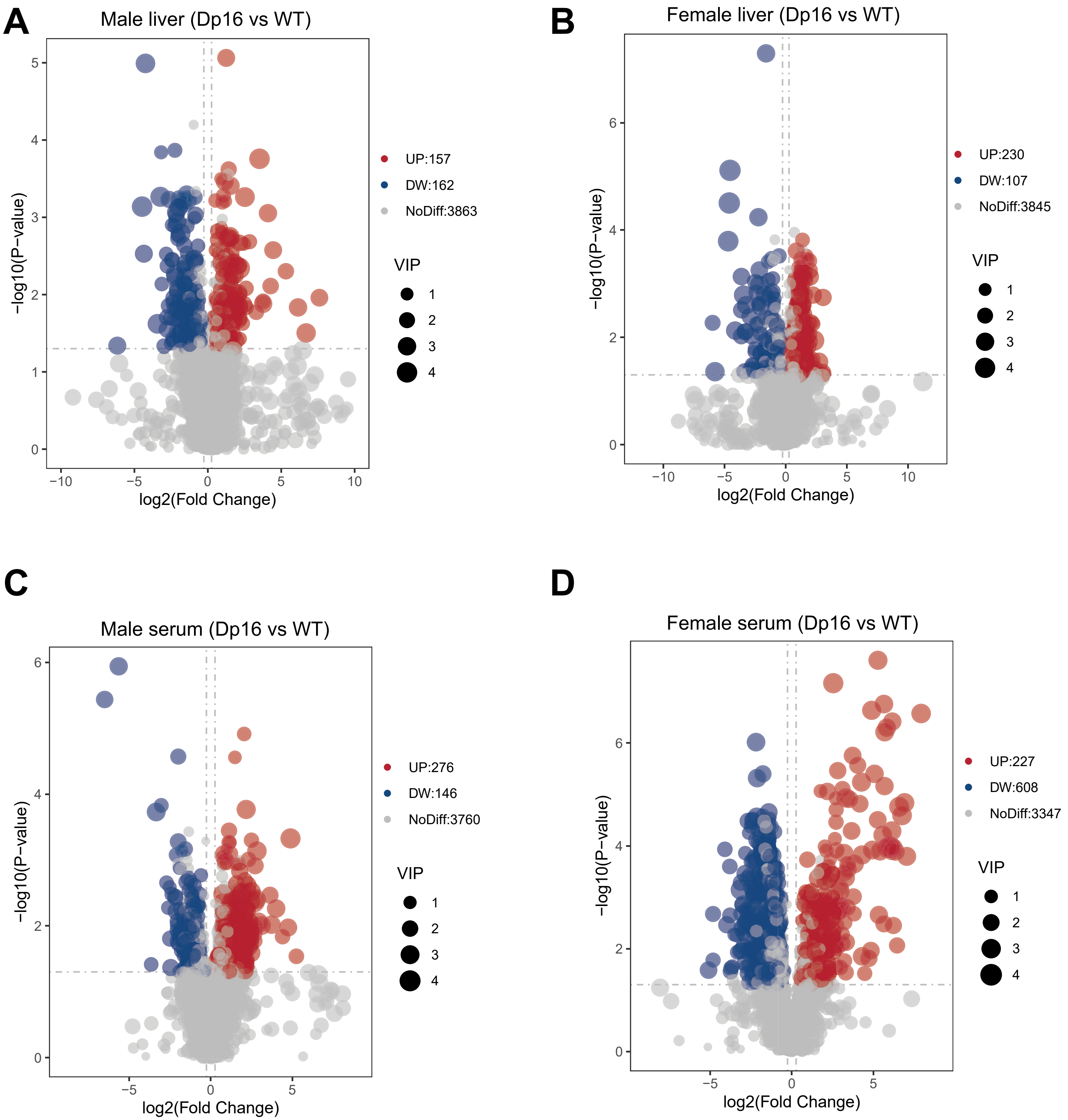
A
Male liver (Dp16 vs WT)
−log10(P−value)
log2(Fold Change)
UP:157
DW:162
NoDiff:3863
VIP
1
2
3
4
B
Female liver (Dp16 vs WT)
−log10(P−value)
log2(Fold Change)
UP:230
DW:107
NoDiff:3845
VIP
C
Male serum (Dp16 vs WT)
−log10(P−value)
log2(Fold Change)
UP:276
DW:146
NoDiff:3760
VIP
D
Female serum (Dp16 vs WT)
−log10(P−value)
log2(Fold Change)
UP:227
DW:608
NoDiff:3347
VIP

**A** Male liver (Dp16 vs WT)

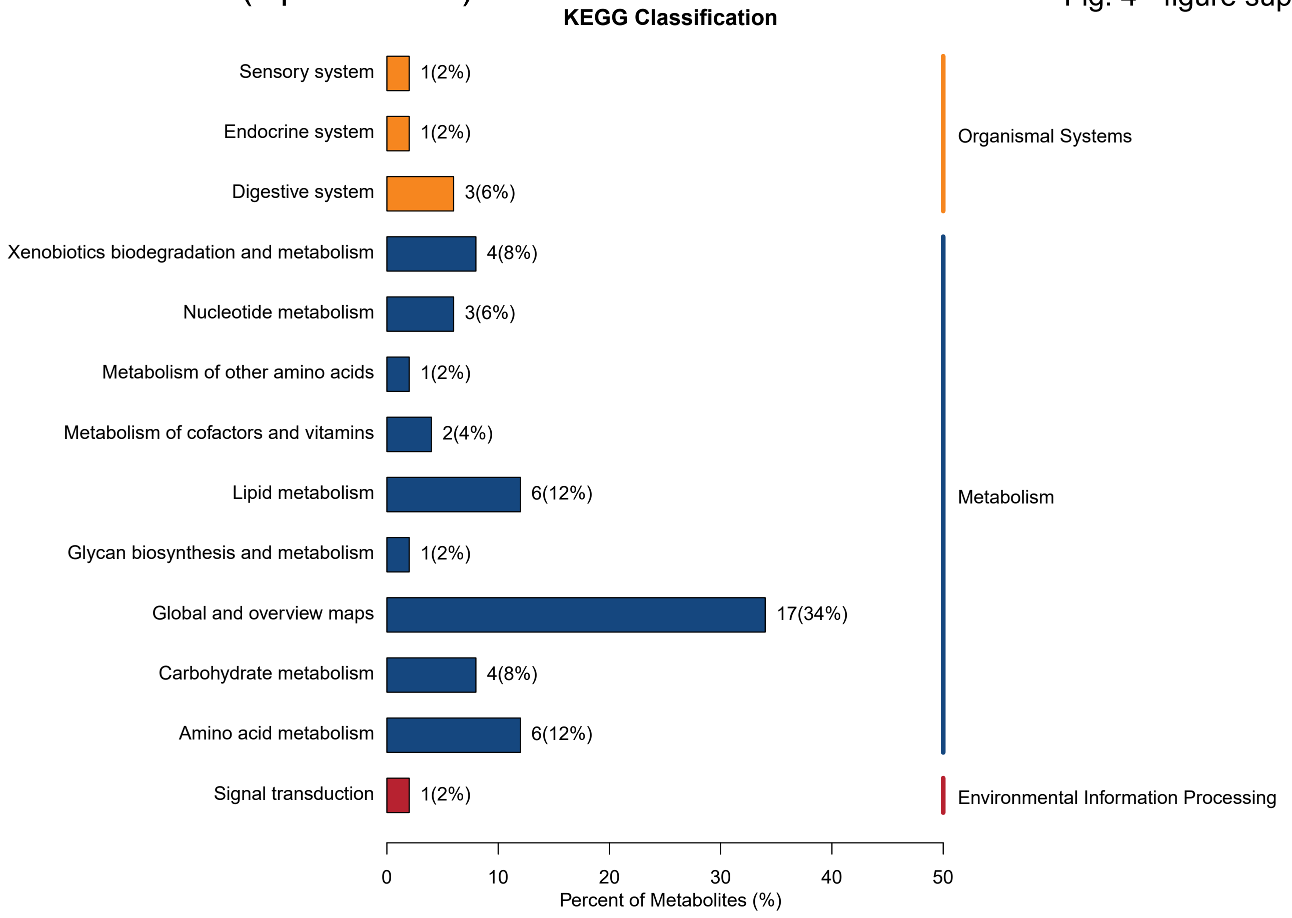


**B** Female liver (Dp16 vs WT)

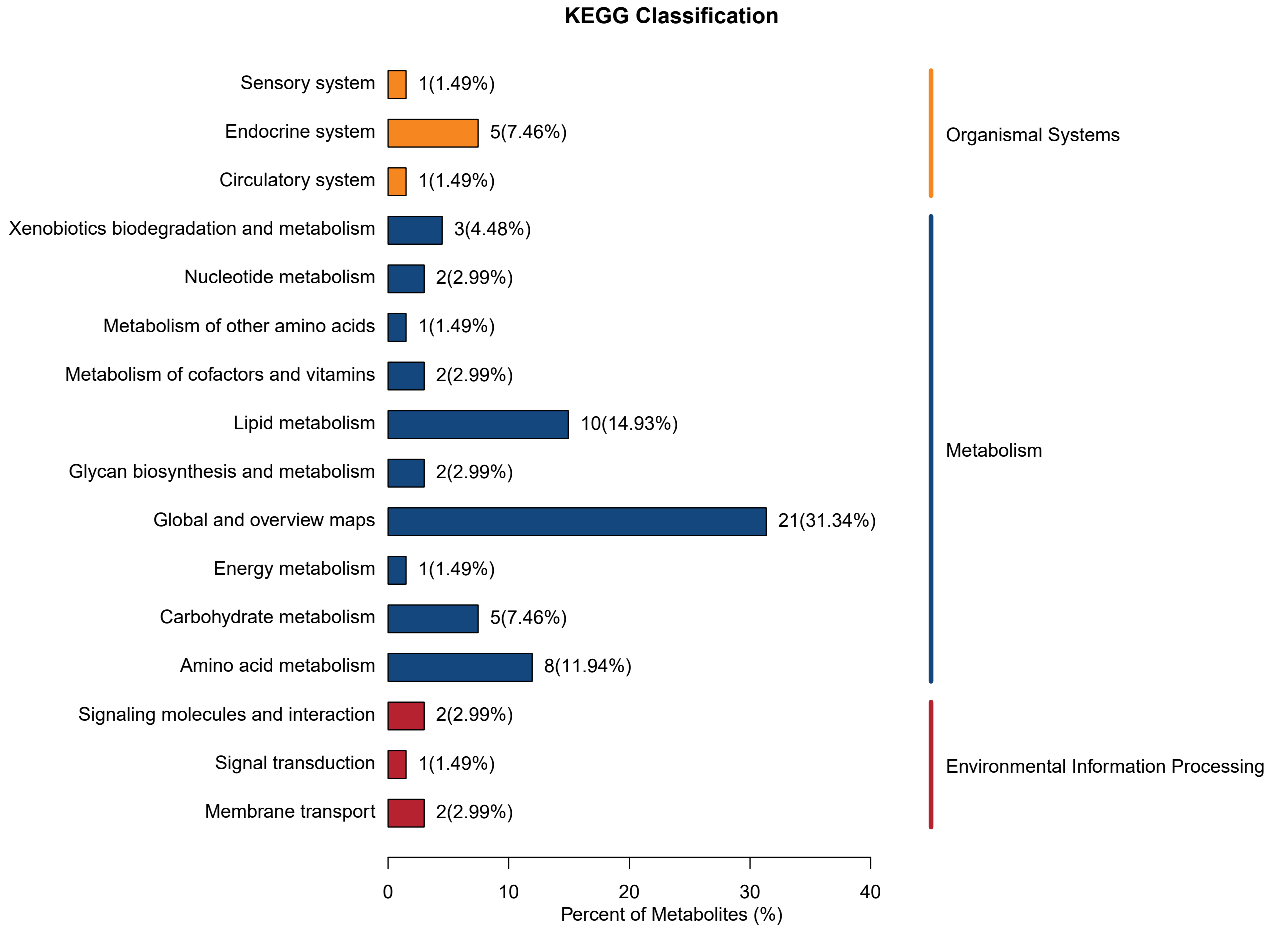

Fig. 4 - figure supplement 3

**A** Male serum (Dp16 vs WT)

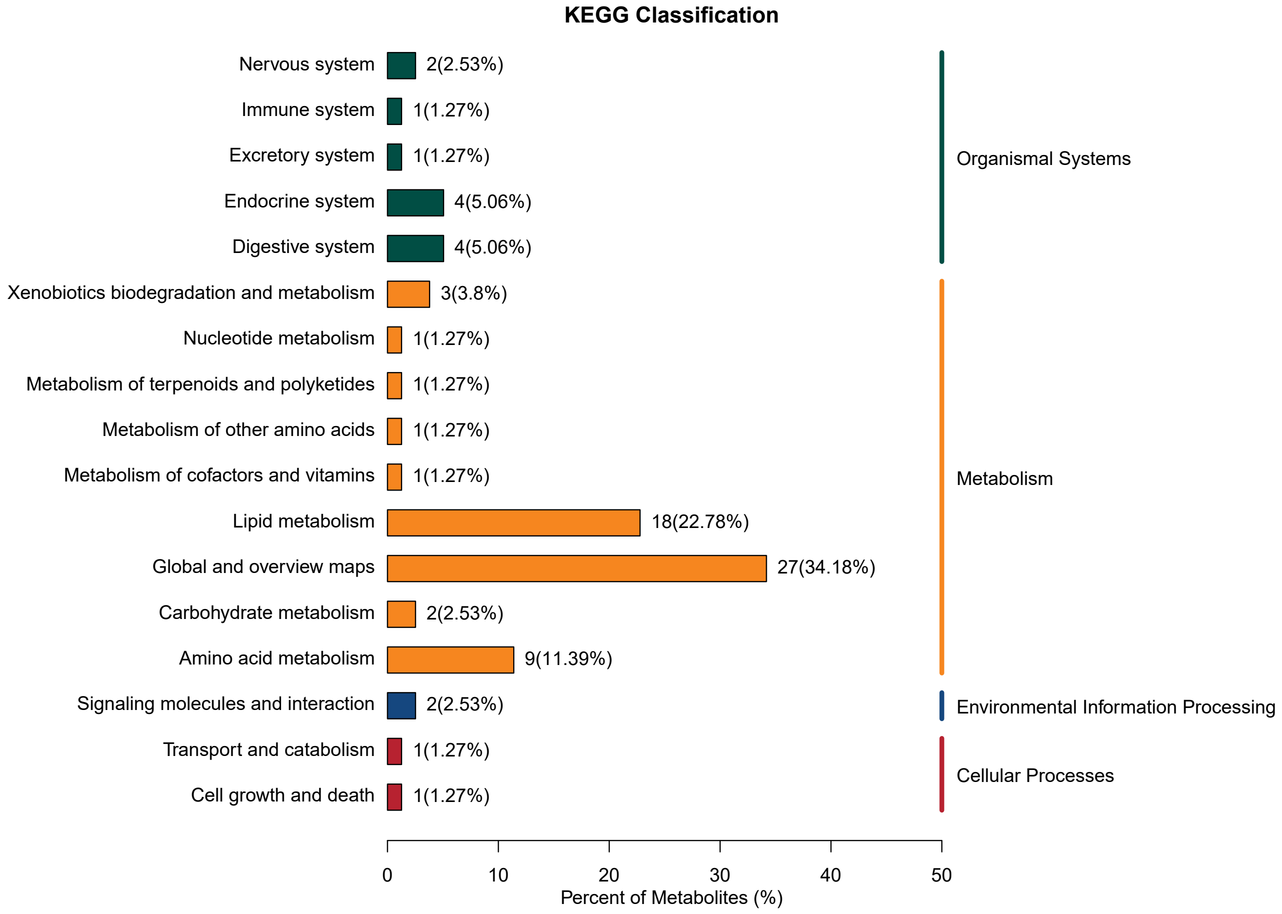


**B** Female serum (Dp16 vs WT)

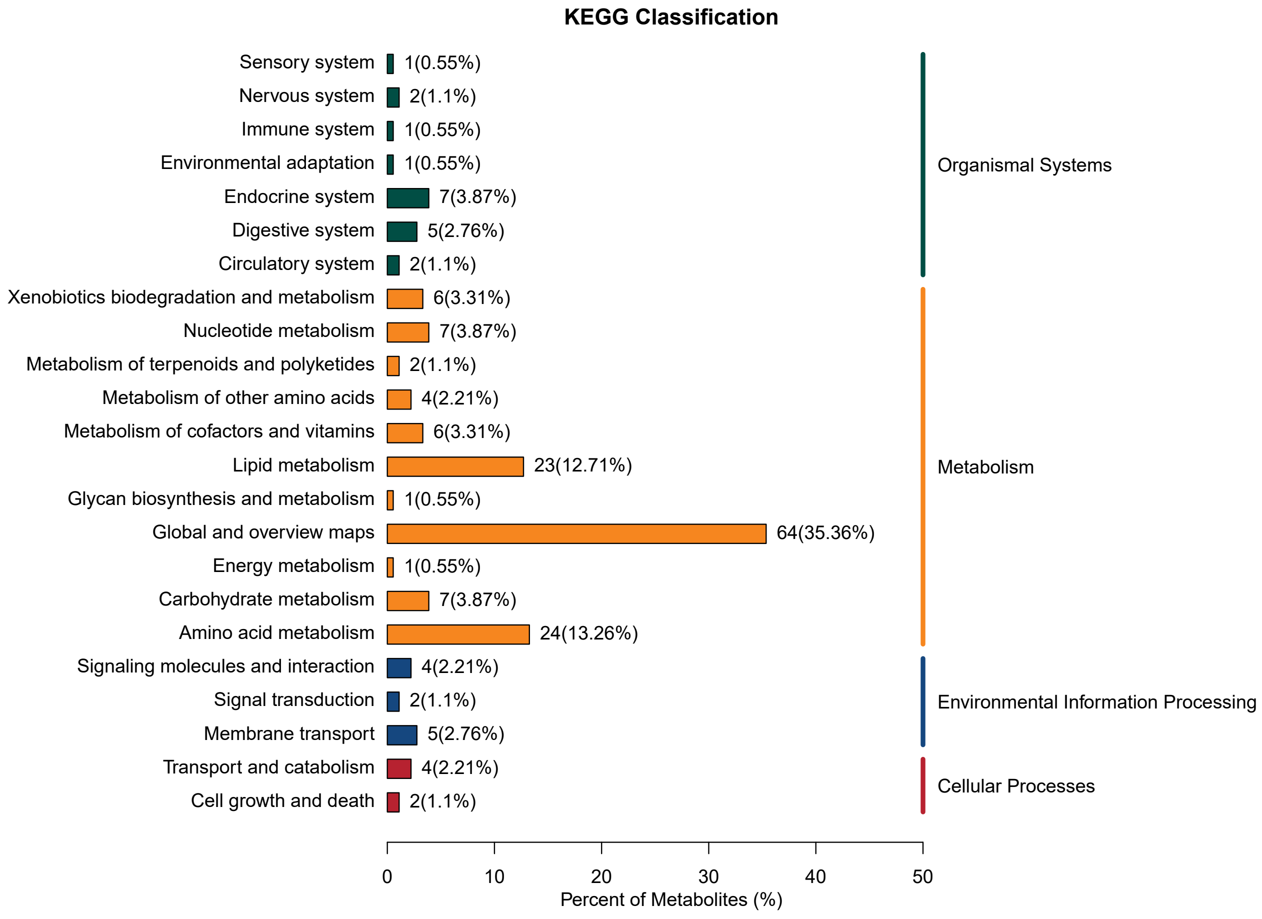

Fig. 4 - figure supplement 4

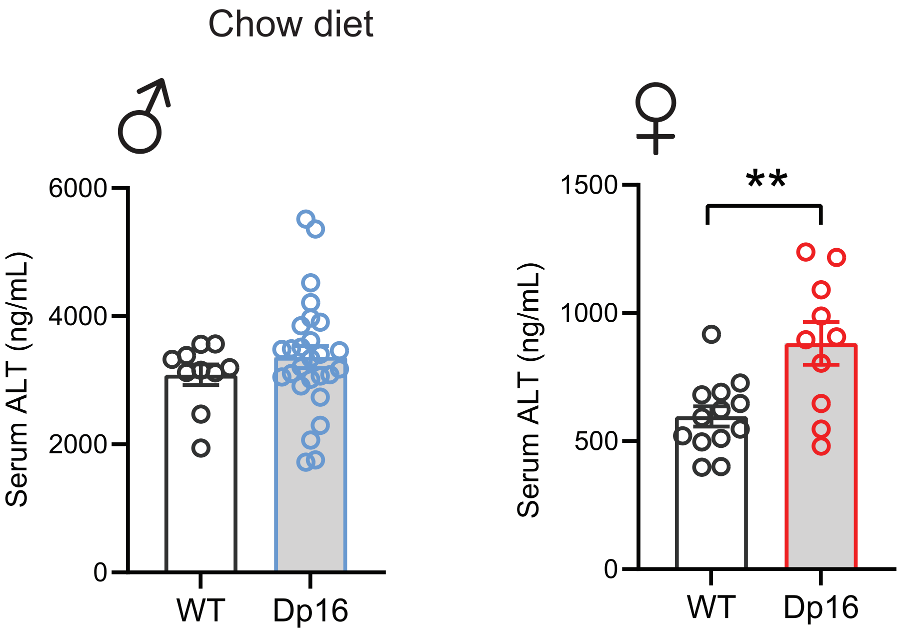

Fig. 5 - figure supplement 1

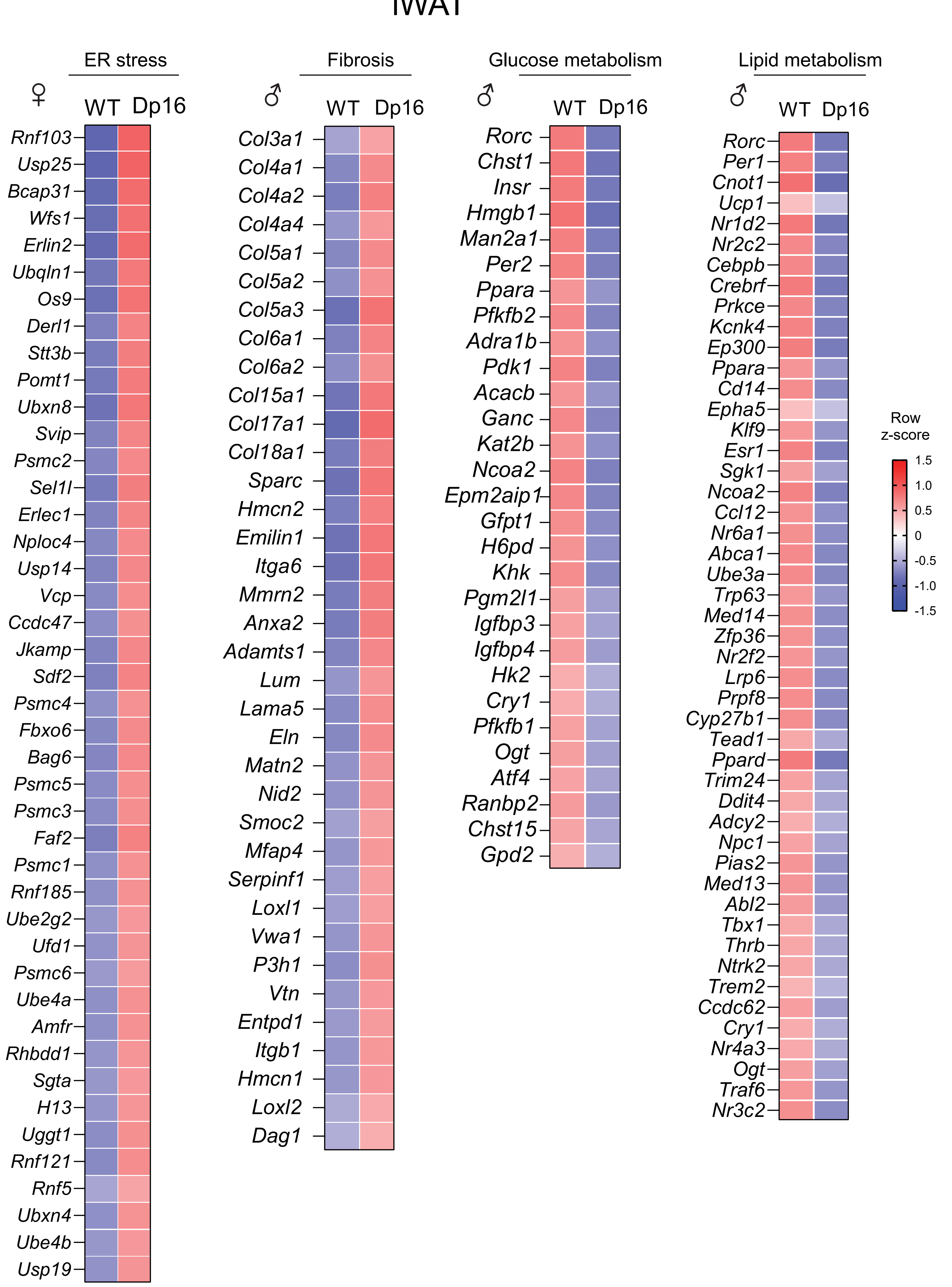

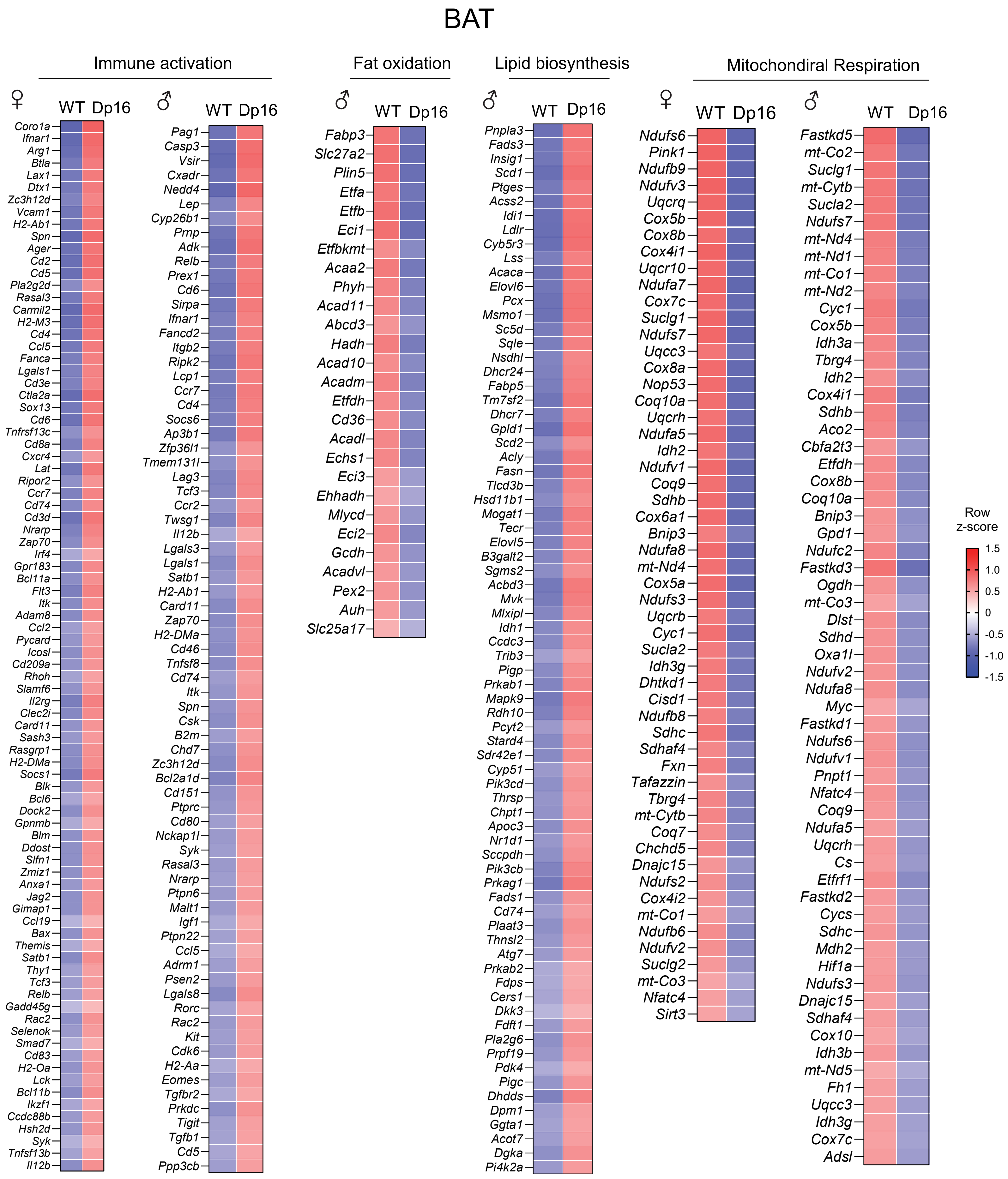
BAT
Immune activation
Fat oxidation
Lipid biosynthesis
Mitochondiral Respiration
♀
♂
WT
Dp16
Coro1a Ifnar1 Arg1 Btla Lax1 Dtx1 Zc3h12d Vcam1 H2-Ab1 Spn Ager Cd2 Cd5 Pla2g2d Rasal3 Carmil2 H2-M3 Cd4 Ccl5 Fanca Lgals1 Cd3e Ctla2a Sox13 Cd6 Tnfrsf13c Cd8a Cxcr4 Lat Ripor2 Ccr7 Cd74 Cd3d Nrarp Zap70 Irf4 Gpr183 Bcl11a Flt3 Itk Adam8 Ccl2 Pycard Icosl Cd209a Rhoh Slamf6 Il2rg Clec2i Card11 Sash3 Rasgrp1 H2-DMa Socs1 Blk Bcl6 Dock2 Gpnmb Blm Ddost Slfn1 Zmiz1 Anxa1 Jag2 Gimap1 Ccl19 Bax Themis Satb1 Thy1 Tcf3 Relb Gadd45g Rac2 Selenok Smad7 Cd83 H2-Oa Lck Bcl11b Ikzf1 Ccdc88b Hsh2d Syk Tnfsf13b Il12b
Pag1 Casp3 Vsir Cxadr Nedd4 Lep Cyp26b1 Prnp Adk Relb Prex1 Cd6 Sirpa Ifnar1 Fancd2 Itgb2 Ripk2 Lcp1 Ccr7 Cd4 Socs6 Ap3b1 Zfp36l1 Tmem131l Lag3 Tcf3 Ccr2 Twsg1 Il12b Lgals3 Lgals1 Satb1 H2-Ab1 Card11 Zap70 H2-DMa Cd46 Tnfsf8 Cd74 Itk Spn Csk B2m Chd7 Zc3h12d Bcl2a1d Cd151 Ptprc Cd80 Nckap1l Syk Rasal3 Nrarp Ptpn6 Malt1 Igf1 Ptpn22 Ccl5 Adrm1 Psen2 Lgals8 Rorc Rac2 Kit Cdk6 H2-Aa Eomes Tgfbr2 Prkdc Tigit Tgfb1 Cd5 Ppp3cb
Fabp3 Slc27a2 Plin5 Etfa Etfb Eci1 Etfbkmt Acaa2 Phyh Acad11 Abcd3 Hadh Acad10 Acadm Etfdh Cd36 Acadl Echs1 Eci3 Ehhadh Mlycd Eci2 Gcdh Acadvl Pex2 Auh Slc25a17
Pnpla3 Fads3 Insig1 Scd1 Ptges Acss2 Idi1 Ldlr Cyb5r3 Lss Acaca Elovl6 Pcx Msmo1 Sc5d Sqle Nsdhl Dhcr24 Fabp5 Tm7sf2 Dhcr7 Gpld1 Scd2 Acly Fasn Tlcd3b Hsd11b1 Mogat1 Tecr Elovl5 B3galt2 Sgms2 Acbd3 Mvk Mlxipl Idh1 Ccdc3 Trib3 Pigp Prkab1 Mapk9 Rdh10 Pcyt2 Stard4 Sdr42e1 Cyp51 Pik3cd Thrsp Chpt1 Apoc3 Nr1d1 Sccpdh Pik3cb Prkag1 Fads1 Cd74 Plaat3 Thnsl2 Atg7 Prkab2 Fdps Cers1 Dkk3 Fdft1 Pla2g6 Prpf19 Pdk4 Pigc Dhdds Dpm1 Ggta1 Acot7 Dgka Pi4k2a
Ndufs6 Pink1 Ndufb9 Ndufv3 Uqcrq Cox5b Cox8b Cox4i1 Uqcr10 Ndufa7 Cox7c Suclg1 Ndufs7 Uqcc3 Cox8a Nop53 Coq10a Uqcrh Ndufa5 Idh2 Ndufv1 Coq9 Sdhb Cox6a1 Bnip3 Ndufa8 mt-Nd4 Cox5a Ndufs3 Uqcrb Cyc1 Sucla2 Idh3g Dhtkd1 Cisd1 Ndufb8 Sdhc Sdhaf4 Fxn Tafazzin Tbrg4 mt-Cytb Coq7 Chchd5 Dnajc15 Ndufs2 Cox4i2 mt-Co1 Ndufb6 Ndufv2 Suclg2 mt-Co3 Nfatc4 Sirt3
Fastkd5 mt-Co2 Suclg1 mt-Cytb Sucla2 Ndufs7 mt-Nd4 mt-Nd1 mt-Co1 mt-Nd2 Cyc1 Cox5b Idh3a Tbrg4 Idh2 Cox4i1 Sdhb Aco2 Cbfa2t3 Etfdh Cox8b Coq10a Bnip3 Gpd1 Ndufc2 Fastkd3 Ogdh mt-Co3 Dlst Sdhd Oxa1l Ndufv2 Ndufa8 Myc Fastkd1 Ndufs6 Ndufv1 Pnpt1 Nfatc4 Coq9 Ndufa5 Uqcrh Cs Etfrf1 Fastkd2 Cycs Sdhc Mdh2 Hif1a Ndufs3 Dnajc15 Sdhaf4 Cox10 Idh3b mt-Nd5 Fh1 Uqcc3 Idh3g Cox7c Adsl
Row z-score
1.5
1.0
0.5
0
-0.5
-1.0
-1.5

Fig. 5 - figure supplement 3

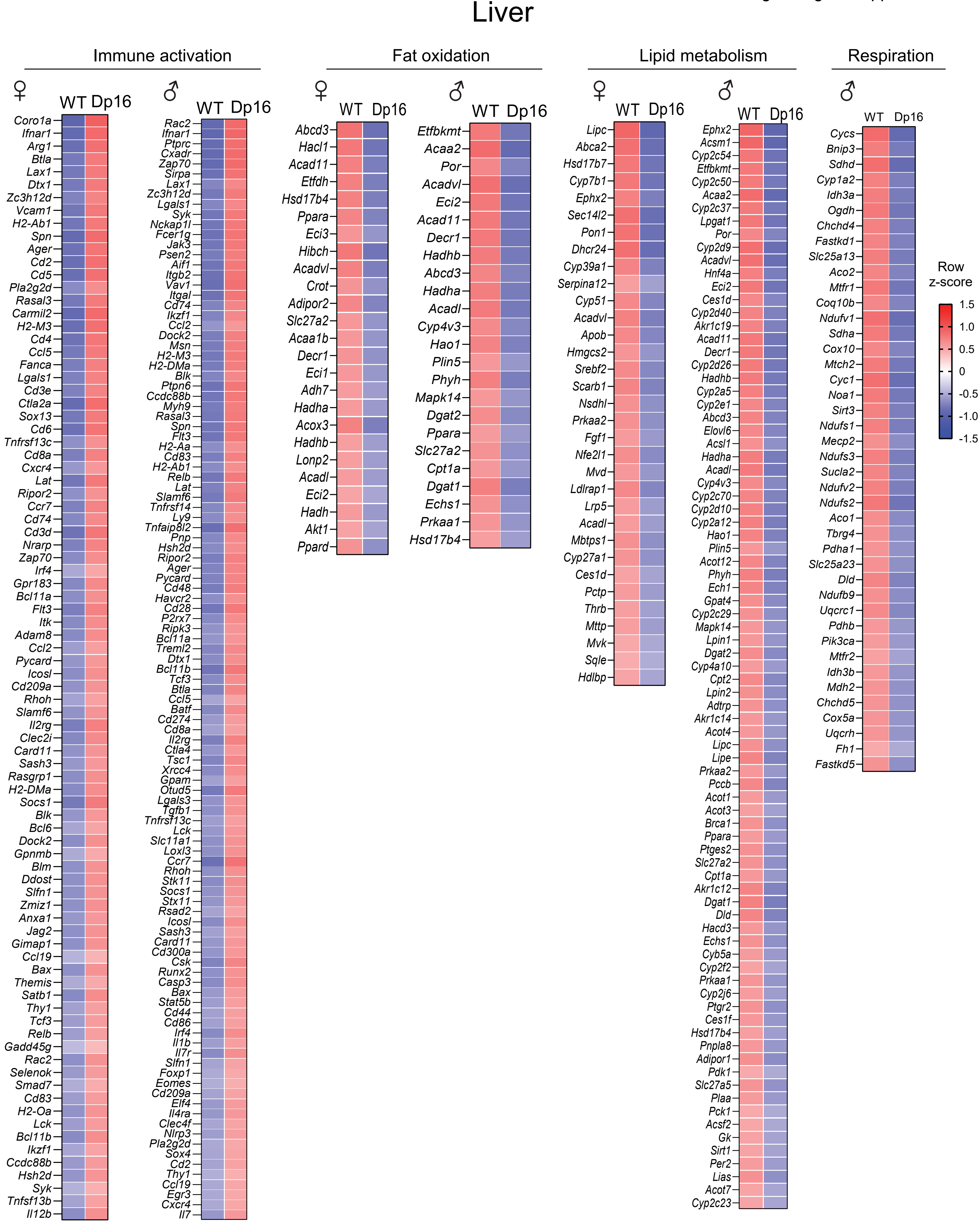

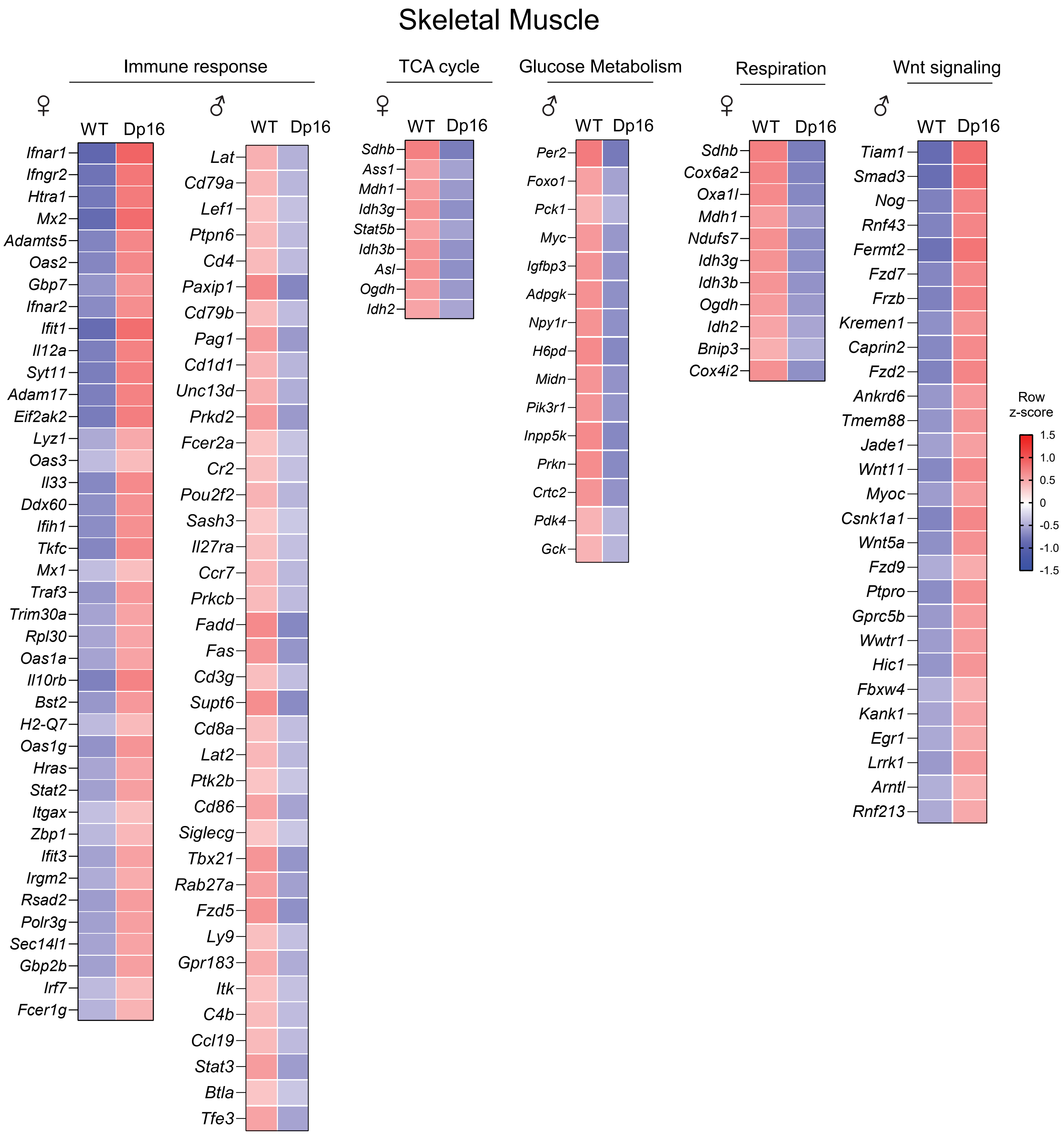
Skeletal Muscle
Immune response
TCA cycle
Glucose Metabolism
Respiration
Wnt signaling
♀
♂
WT
Dp16
Ifnar1
Ifngr2
Htra1
Mx2
Adamts5
Oas2
Gbp7
Ifnar2
Ifit1
Il12a
Syt11
Adam17
Eif2ak2
Lyz1
Oas3
Il33
Ddx60
Ifih1
Tkfc
Mx1
Traf3
Trim30a
Rpl30
Oas1a
Il10rb
Bst2
H2-Q7
Oas1g
Hras
Stat2
Itgax
Zbp1
Ifit3
Irgm2
Rsad2
Polr3g
Sec14l1
Gbp2b
Irf7
Fcer1g
Lat
Cd79a
Lef1
Ptpn6
Cd4
Paxip1
Cd79b
Pag1
Cd1d1
Unc13d
Prkd2
Fcer2a
Cr2
Pou2f2
Sash3
Il27ra
Ccr7
Prkcb
Fadd
Fas
Cd3g
Supt6
Cd8a
Lat2
Ptk2b
Cd86
Siglecg
Tbx21
Rab27a
Fzd5
Ly9
Gpr183
Itk
C4b
Ccl19
Stat3
Btla
Tfe3
Sdhb
Ass1
Mdh1
Idh3g
Stat5b
Idh3b
Asl
Ogdh
Idh2
Per2
Foxo1
Pck1
Myc
Igfbp3
Adpgk
Npy1r
H6pd
Midn
Pik3r1
Inpp5k
Prkn
Crtc2
Pdk4
Gck
Sdhb
Cox6a2
Oxa1l
Mdh1
Ndufs7
Idh3g
Idh3b
Ogdh
Idh2
Bnip3
Cox4i2
Tiam1
Smad3
Nog
Rnf43
Fermt2
Fzd7
Frzb
Kremen1
Caprin2
Fzd2
Ankrd6
Tmem88
Jade1
Wnt11
Myoc
Csnk1a1
Wnt5a
Fzd9
Ptpro
Gprc5b
Wwtr1
Hic1
Fbxw4
Kank1
Egr1
Lrrk1
Arntl
Rnf213
Row z-score
1.5
1.0
0.5
0
-0.5
-1.0
-1.5

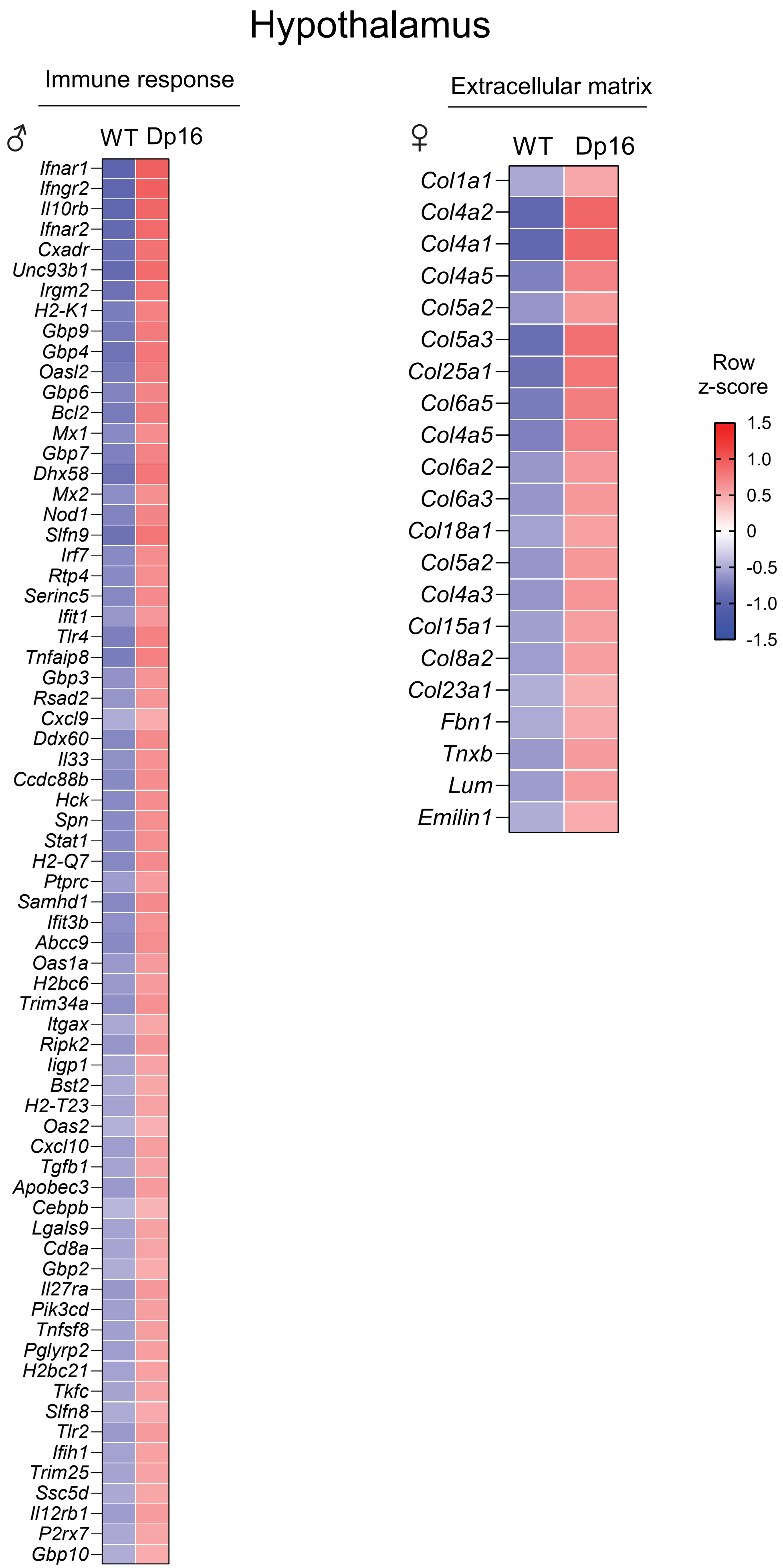
Hypothalamus
Immune response
♂
WT
Dp16
Ifnar1
Ifngr2
Il10rb
Ifnar2
Cxadr
Unc93b1
Irgm2
H2-K1
Gbp9
Gbp4
Oasl2
Gbp6
Bcl2
Mx1
Gbp7
Dhx58
Mx2
Nod1
Slfn9
Irf7
Rtp4
Serinc5
Ifit1
Tlr4
Tnfaip8
Gbp3
Rsad2
Cxcl9
Ddx60
Il33
Ccdc88b
Hck
Spn
Stat1
H2-Q7
Ptprc
Samhd1
Ifit3b
Abcc9
Oas1a
H2bc6
Trim34a
Itgax
Ripk2
Iigp1
Bst2
H2-T23
Oas2
Cxcl10
Tgfb1
Apobec3
Cebpb
Lgals9
Cd8a
Gbp2
Il27ra
Pik3cd
Tnfsf8
Pglyrp2
H2bc21
Tkfc
Slfn8
Tlr2
Ifih1
Trim25
Ssc5d
Il12rb1
P2rx7
Gbp10
Extracellular matrix
♀
WT
Dp16
Col1a1
Col4a2
Col4a1
Col4a5
Col5a2
Col5a3
Col25a1
Col6a5
Col4a5
Col6a2
Col6a3
Col18a1
Col5a2
Col4a3
Col15a1
Col8a2
Col23a1
Fbn1
Tnxb
Lum
Emilin1
Row
z-score
1.5
1.0
0.5
0
-0.5
-1.0
-1.5

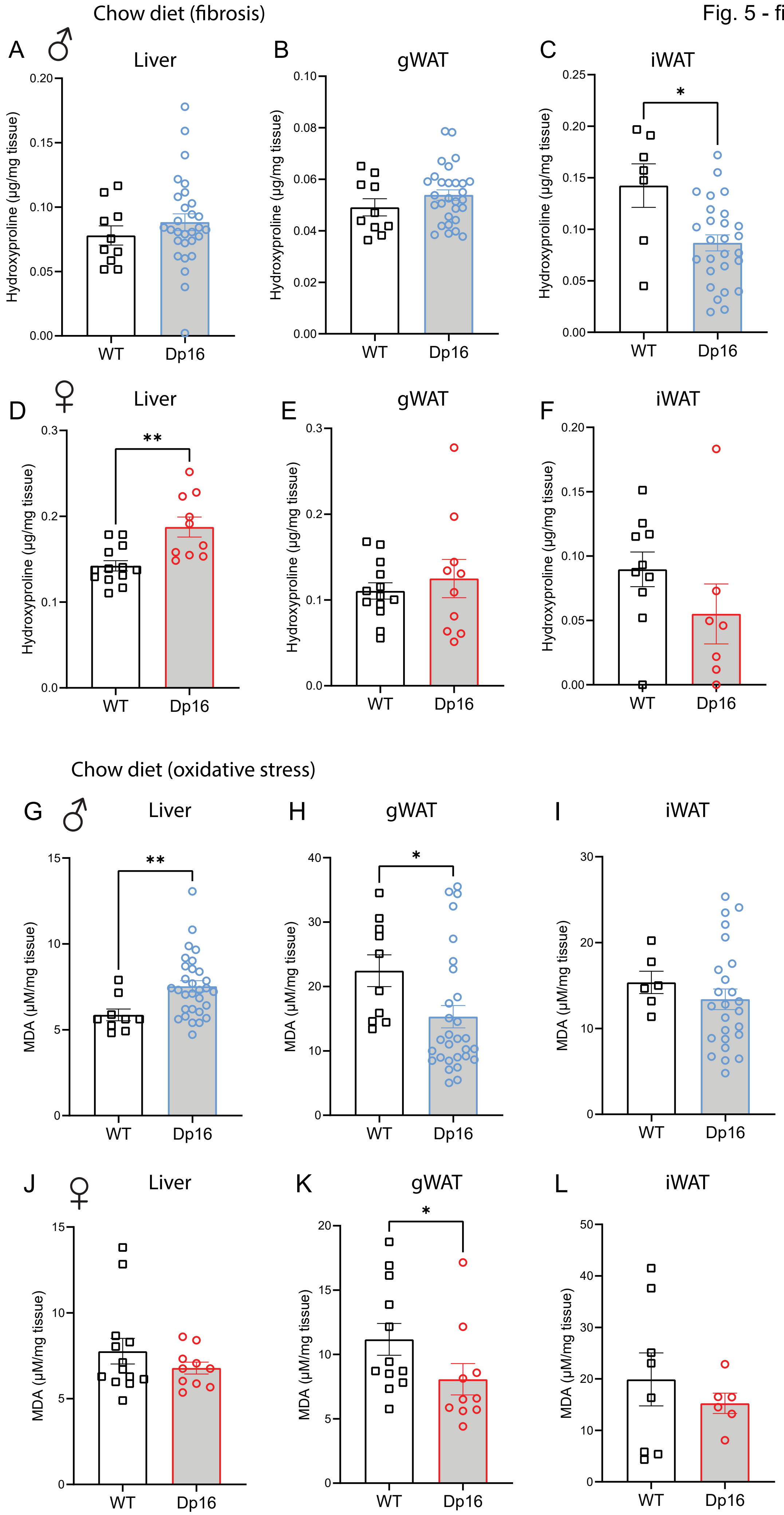
Chow diet (fibrosis)
A ♂ Liver
B gWAT
C iWAT
D ♀ Liver
E gWAT
F iWAT
Hydroxyproline (µg/mg tissue)
WT
Dp16
*
**
Chow diet (oxidative stress)
G ♂ Liver
H gWAT
I iWAT
J ♀ Liver
K gWAT
L iWAT
MDA (µM/mg tissue)

Fig. 6 - figure supplement 1

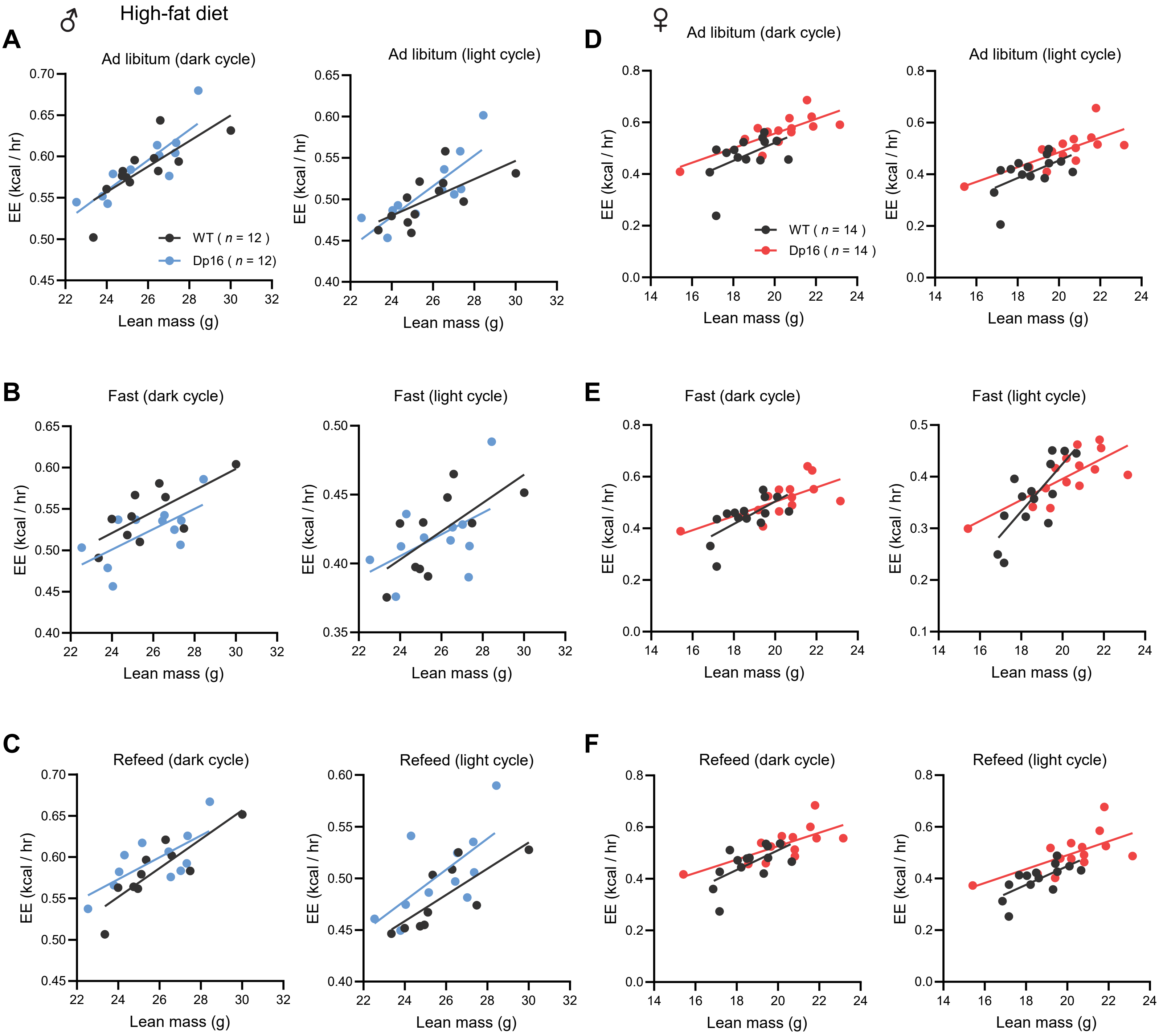

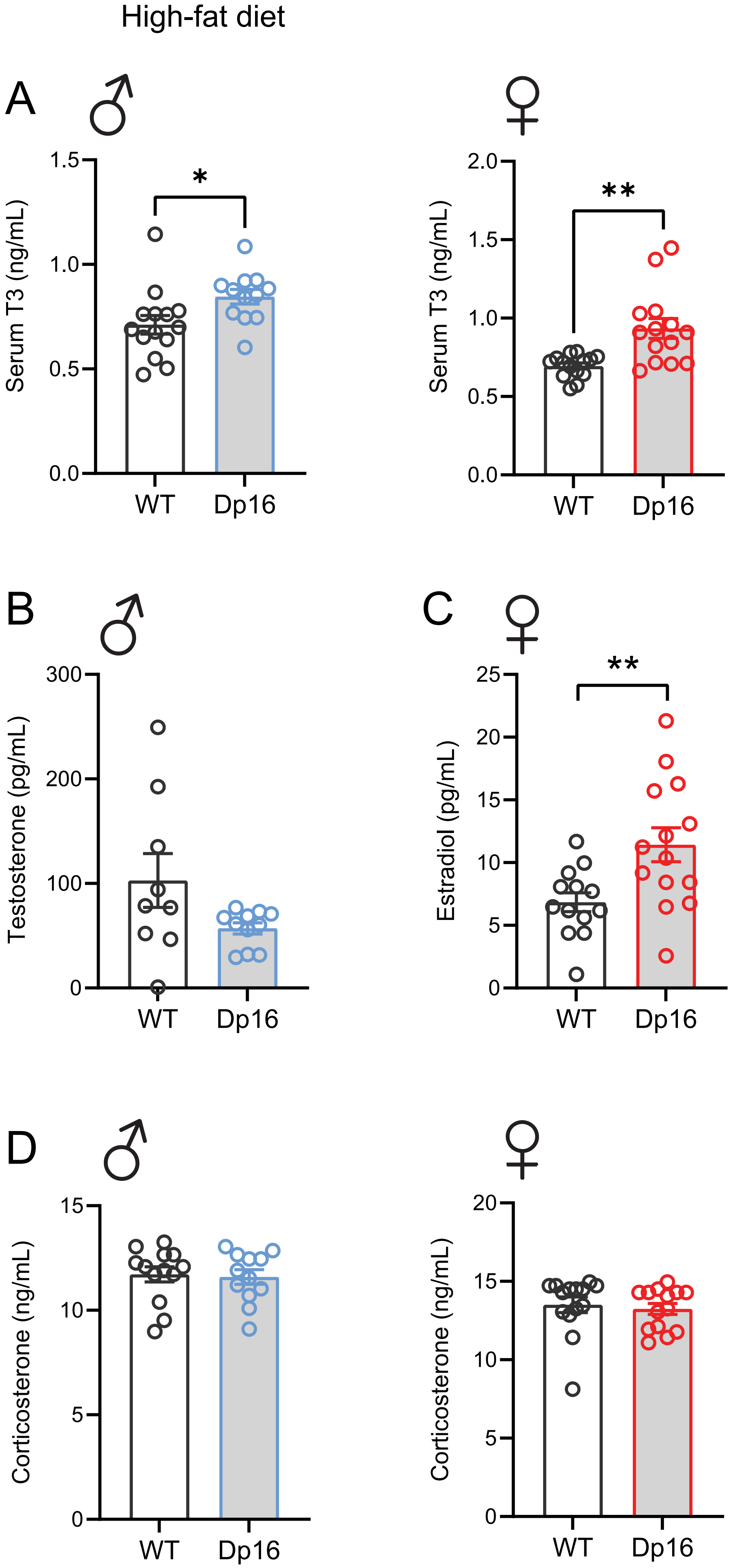
High-fat diet
A
♂
♀
Serum T3 (ng/mL)
1.5
1.0
0.5
0.0
*
WT
Dp16
Serum T3 (ng/mL)
2.0
1.5
1.0
0.5
0.0
**
WT
Dp16
B
♂
Testosterone (pg/mL)
300
200
100
0
WT
Dp16
C
♀
Estradiol (pg/mL)
25
20
15
10
5
0
**
WT
Dp16
D
♂
♀
Corticosterone (ng/mL)
15
10
5
0
WT
Dp16
Corticosterone (ng/mL)
20
15
10
5
0
WT
Dp16

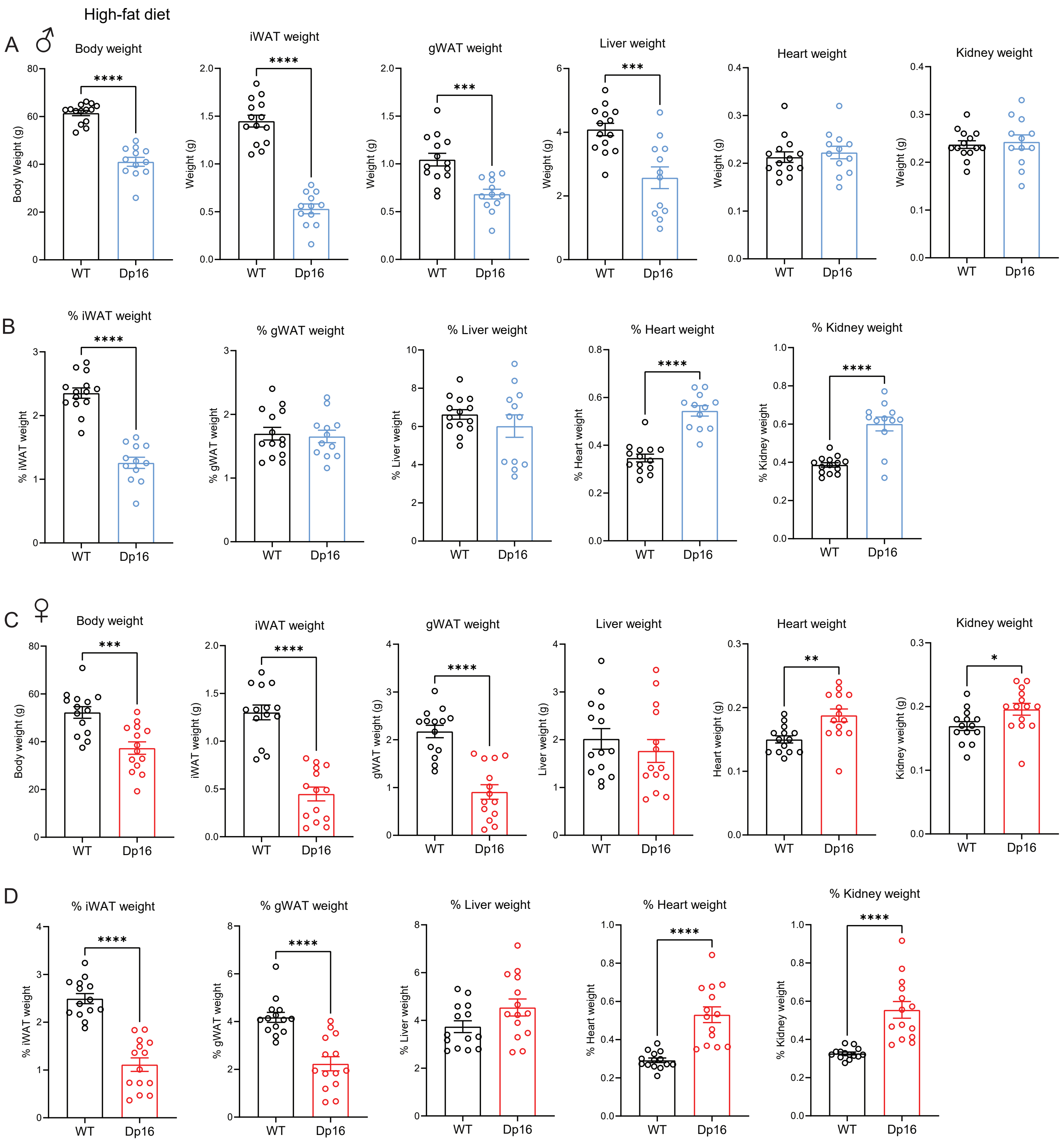
High-fat diet
A ♂
Body weight
iWAT weight
gWAT weight
Liver weight
Heart weight
Kidney weight
Body Weight (g)
Weight (g)
****
***
WT
Dp16
B
% iWAT weight
% gWAT weight
% Liver weight
% Heart weight
% Kidney weight
C ♀
Body weight
iWAT weight
gWAT weight
Liver weight
Heart weight
Kidney weight
Body weight (g)
iWAT weight (g)
gWAT weight (g)
Liver weight (g)
Heart weight (g)
Kidney weight (g)
**
*
D
% iWAT weight
% gWAT weight
% Liver weight
% Heart weight
% Kidney weight

Fig. 6 - figure supplement 4

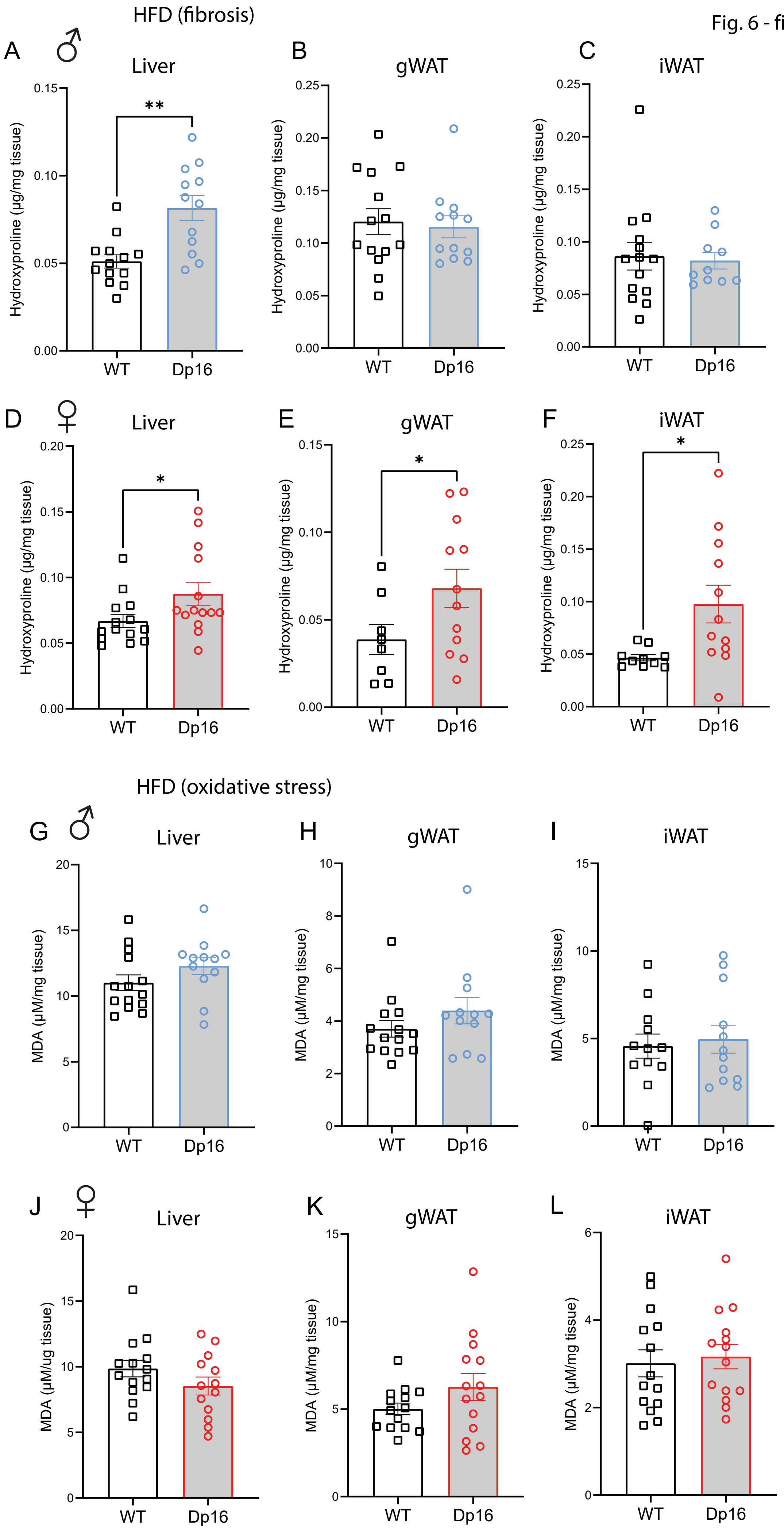
